# Punctuated and continuous structural diversity of S-layers across the prokaryotic tree of life

**DOI:** 10.1101/2024.05.28.596244

**Authors:** Edward Johnston, Buse Isbilir, Vikram Alva, Tanmay A.M. Bharat, Jonathan P. K. Doye

## Abstract

Surface layers (S-layers) are two-dimensional (2D) crystalline lattices that frequently coat prokaryotic cells, playing a crucial role in protection, maintaining cellular integrity, and mediating environmental interactions. However, the molecular landscape of these abundant proteins has remained underexplored due to a lack of structural data. By employing AlphaFold2multimer together with planar symmetry constraints in a workflow validated by electron cryomicroscopy structure determination, we have elucidated the lattice structures of over 150 S-layers from diverse archaea and bacteria. Our findings unveil a multifaceted evolutionary landscape for S-layer proteins, highlighting key differences in the evolution of bacterial and archaeal S-layers. Our study allows us to discover underlying patterns in S-layer structure, organisa-tion, and cell anchoring mechanisms across the prokaryotic tree of life, deepening our understanding of the intricately complex microbial cell surfaces, which appear to have evolved proteinaceous S-layers independently on multiple occasions. This work will open avenues for rational manipulation of prokaryotic cellular interactions in multicellular microbiomes, as well as for innovative 2D biomaterial design.

## Introduction

In numerous biological contexts, ranging from components of eukaryotic cells such as coated vesicles and microtubules (*1*), to the capsids of viruses (*2*) and the surface layers (S-layers) of prokaryotes (*3*), many proteins are known to form highly ordered two-dimensional (2D) lattice assemblies. These structured assemblies fulfil important biological functions that their monomeric forms cannot. Although recent advances in structure prediction have enabled the accurate modeling of monomeric proteins (*4*), predicting the arrangements of these proteins in ordered extended assemblies remains a major challenge.

Among these assemblies, the S-layers of prokaryotes are notable for their highly organised, paracrystalline 2D lattices that envelop entire cells (*3, 5–7*). S-layers, composed of repeating units of S-layer proteins (SLPs) (*8,9*), are found across archaea (*10–12*) and both Gram-negative (diderm) (*13*) and Gram-positive (monoderm) bacteria (*14–17*). Remarkably, SLPs are often the proteins with the highest copy number in these cells (*3*), leading to the proposition that SLPs are among the most abundant classes of proteins in nature (*3, 18*).

The production and maintenance of an S-layer on a prokaryotic cell requires precise orchestration of cellular machinery (*19, 20*), imposing considerable energetic demands on the cell (*21, 22*). Therefore, it is not surprising that S-layers in bacteria and archaea fulfill many important functions, including maintenance of cell shape, protection from predators (*23*), regula-tion of nutrient uptake (*24*), and pathogenicity (*25*). While early studies reported low-resolution electron crystallography structures of S-layers several decades ago (*26–28*), the characterisation of S-layers at the atomic level has only recently become possible using advanced structural biology techniques (*14–17, 29–32*). Yet, experimental structural studies of S-layers are labor-intensive and time-consuming, usually requiring a range of methods to accurately determine and validate structures (*33*). As a result, the structural and evolutionary landscapes of S-layers remain largely uncharted. Furthermore, while imaging studies have detected S-layers in several species, the specific SLPs forming the S-layer frequently remain unidentified. In addition, for numerous taxonomic groups, the presence of S-layers is yet to be confirmed. This means that the diversity, lattice arrangements, and the cell-anchoring mechanisms of S-layers remain poorly understood. Given that S-layers represent the outermost cellular organelle in many prokaryotes (*3, 7*), the paucity of S-layer structures significantly limits our understanding of how microorganisms interact with and inhabit their specific ecological niches on Earth.

To bridge this gap in our fundamental understanding, we leverage the power of contemporary bioinformatics and protein structure prediction techniques, specifically AlphaFold-multimer (*34*), to identify and predict the 2D crystal structures of over 150 S-layers across the prokaryotic tree of life. Our analyses reveal significant diversity in the structures of S-layers across all evaluated dimensions, including symmetry, porosity, and mechanisms of cell anchoring, as well as in their secondary, tertiary, and quarternary protein structures. Notably, our findings demonstrate that the structural and sequence landscapes of archaeal S-layers are predominantly evolutionarily continuous, when compared with those of bacterial S-layers, which display substantial discontinuity, indicating multiple independent evolutionary origins for bacterial S-layers.

## Results

AlphaFold-multimer cannot directly predict the structure of a crystal, but only the structure of a discrete complex. To extend this method to 2D crystals, our approach assumes that the structural predictions for *n*-mers can provide a comprehensive insight into the structure of the S-layer around symmetry sites with *n*-fold rotational symmetry. Hence, we run calculations for complexes of the SLP, including fragments thereof, in configurations ranging from homodimers (2 monomers) to homoheptamers (7 monomers). In assessing our results, we rely on the confidence metrics of AlphaFold, which have been shown to be impressively robust (*35*), both to rank the predictions for a particular *n*-mer and to identify which cluster sizes are most stable.

We illustrate our method in Fig. 1 (also see Figs. S1-S4) for the SLP of *Corynebacterium glutamicum* (*36*), a Gram-positive bacterium widely used for industrial-scale synthesis of amino acids. Unlike the majority of known S-layer lattices, which are predominantly composed of *β*-strand-rich folds (*14, 16, 37*), the SLP of *C. glutamicum*, referred to as PS2, is entirely *α*helical. Notably, its homologs are found exclusively in the Corynebacterium genus, including both pathogenic and non-pathogenic species integral to the human microbiota. From the confidence-metric-based comparison of different oligomers, it becomes evident that interactions towards the C-terminus of PS2 strongly stabilize a hexamer, whereas interactions closer to the N-terminus favor the formation of a trimeric assembly. The structures of the *C*_6_ hexamer and *C*_3_ trimer are shown in Fig. 1B. In the hexamer, the C-terminal *α*-helices wrap around each other in a coiled coil that extends downward, away from the assembly plane of the S-layer. The N-terminal segment of each monomer projects outward, aligning with the plane of the Slayer and providing two distinct interfaces for engaging with the N-terminal segments of two monomers from adjacent hexamers, thereby establishing a trimeric interface. Moreover, PS2 features a predicted transmembrane helix at its C-terminus, which likely anchors the S-layer to the cell membrane.

**Figure 1:**
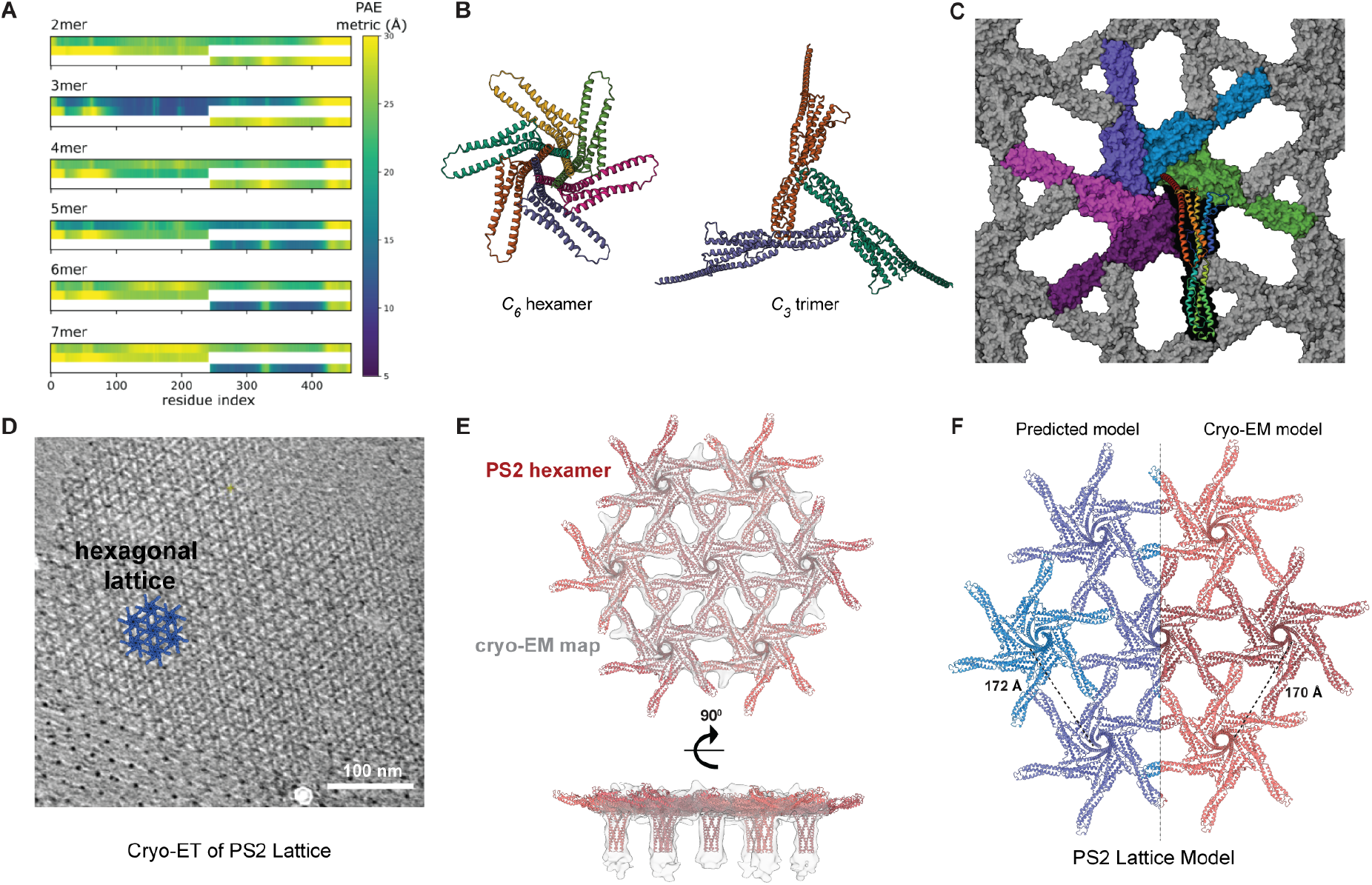
The S-layer of Corynebacterium *glutamicum*. (A) Confidence-metric-based comparison of the predictions for different sized multimers of the whole SLP and sub-proteins consisting of the N- and C-terminal halves. The plots allow both the stable complexes and the parts of the protein involved in interactions to be identified. Low values of the metric correspond to high confidence in inter-residue distances. (B) Identified high-confidence complexes: C-terminal 6-mer and N-terminal 3-mer. (C) Space-filling representation of the predicted 2D lattice with the 6 monomers in the primitive cell represented in colour (one monomer is additionally shown as a ribbon diagram). (D) A tomographic slice through a purified *C. glutamicum* cell envelope showing a hexagonal lattice. (E) Cryo-EM density of the S-layer (in light gray), fitted with *C. glutamicum* S-layer protein PS2 hexamers; top (top view) and bottom (side view). (F) Computationally predicted PS2 lattice with 172 Å lattice constant and PS2 lattice produced by fitting the hexameric PS2 into cryo-EM map with 170 Å lattice constant compared side by side.

To build a crystal model from these complexes, we first generate a symmetrized average model of the sub-proteins: a *C*_3_ symmetrized N-terminal half and a *C*_6_ symmetrized C-terminal half of PS2. We then superimpose the two overlapping sections of the two sub-proteins in the two complexes. This typically leads to a structure wherein the symmetry axes are not perfectly parallel, as they would be in a planar crystal. For the current example, the two axes deviate by an angle of 6°, likely because the C-terminal *α*-helices in the 2D crystal bend further compared to a hypothetical monomer to maximize their interactions in the coiled coil. Therefore, we apply a small distortion to the overlapping section to rotate the symmetry axes to be co-parallel, thus generating the final 2D crystal model of the S-layer (Fig. 1C and S4).

The agreement of our crystal model with previously reported low-resolution atomic force microscopy (AFM) images of the *C. glutamicum* S-layer (*36*) lends strong support to our approach. To further substantiate the validity of our approach, we investigated the structure of the S-layer using electron cryomicroscopy (cryo-EM) and electron cryotomography (cryo-ET) (Figs. 1D-F and S9, Tables S3-S4 and Movie S1). First, we purified S-layers from *C. glutamicum* cells using previously described methods (*11, 36*). Cryo-ET analysis of purified S-layers confirmed the hexagonal arrangement of the S-layer (Fig. 1D), matching closely with the reported results from AFM (*36*). Using a cryo-EM single-particle approach, we obtained a 16.8 Å -resolution reconstruction of the S-layer, which allowed us to unequivocally fit multiple PS2 hexamers into the map (Fig. 1E). The reconstructed map distinctly reveals the overall protein arrangement, hexameric coiled coils, and both hexameric and trimeric pores, with the predicted crystal structure fitting almost perfectly into the cryo-EM map. Notably, the measured lattice constant from our predicted structure (172 Å) matches the experimental value (170 Å) (Fig. 1F), and the dimensions of the trimeric pores in the experimental and predicted sheet structures, which arise directly through formation of the lattice, exhibit remarkable agreement (22.6 Å vs 22.8 Å). Lastly, the congruence in the positioning of supersecondary structural ele-ments, such as the hexameric coiled coil, further validates our approach (Fig. 1E-F).

After confirming the validity of our approach on the S-layer of the bacterium *C. glutamicum*, which contains a primarily *α*-helical SLP that had not been structurally resolved before, we proceeded to test our approach on an archaeal S-layer from *Haloferax volcanii*, which contains mainly *β*-strands and has a known atomic structure (*11*). This S-layer is structurally distinct from the S-layer of *C. glutamicum*, consisting of a series of immunoglobulin (Ig)-like folds. The lattice models for this S-layer, predicted using our methodology (Table S1), showed remarkable agreement with the experimentally determined structure (*11*). Specifically, previous studies reported the experimental lattice constant for the *H. volcanii* S-layer as 168 Å, which closely aligns with our prediction of 164 Å (Figs. S2-S3 and Table S1).

Following the rigorous validation of our approach through examples of experimentally determined archaeal and bacterial S-layer structures, which are both evolutionarily and structurally diverse, we applied the method to study a broad range of S-layers across the prokaryotic tree of life. We began our exploration of the structural diversity of S-layers by considering organisms where the SLP had been identified experimentally. We then extended our analysis to homologs of these proteins (Figs. S5-S6), particularly targeting those organisms whose S-layers have been ultrastructurally characterized. These two sets provided many examples where our predictions matched existing low-resolution reconstructions of the S-layer structures (Fig. 2 and Tables S1-2), further increasing our confidence in the reliability of the approach. This consistency allowed the definitive identification of the SLPs in many of these organisms for the first time. For example, in the thermophilic, biosurfactant-producing Gram-positive bacterium *Aneurinibacillus thermoaerophilus*, the SLP had been previously identified (*38*), but little was known about the structure of the S-layer beyond its *p*4 symmetry and lattice constant (*39*). In fact, this protein has a large number of homologs in the *Bacillota* phyla, including a series of organ-isms for which the similarity of their S-layers in electron microscopy reconstructions had been noted (*39*); this group includes the much-displayed *p*4 S-layer of *Desulfotomaculum nigrificans* (Fig. 2). Moreover, our homology searches initiated with the SLP of the evolutionarily deep-branching extremophile *Deinococcus radiodurans* uncovered divergent SLPs in closely related extremophiles, such as *Deinococcus geothermalis* and *Thermus thermophilus*, simultaneously resolving the previously ambiguous identity of the SLP in *Thermus thermophilus* (*40*).

**Figure 2:**
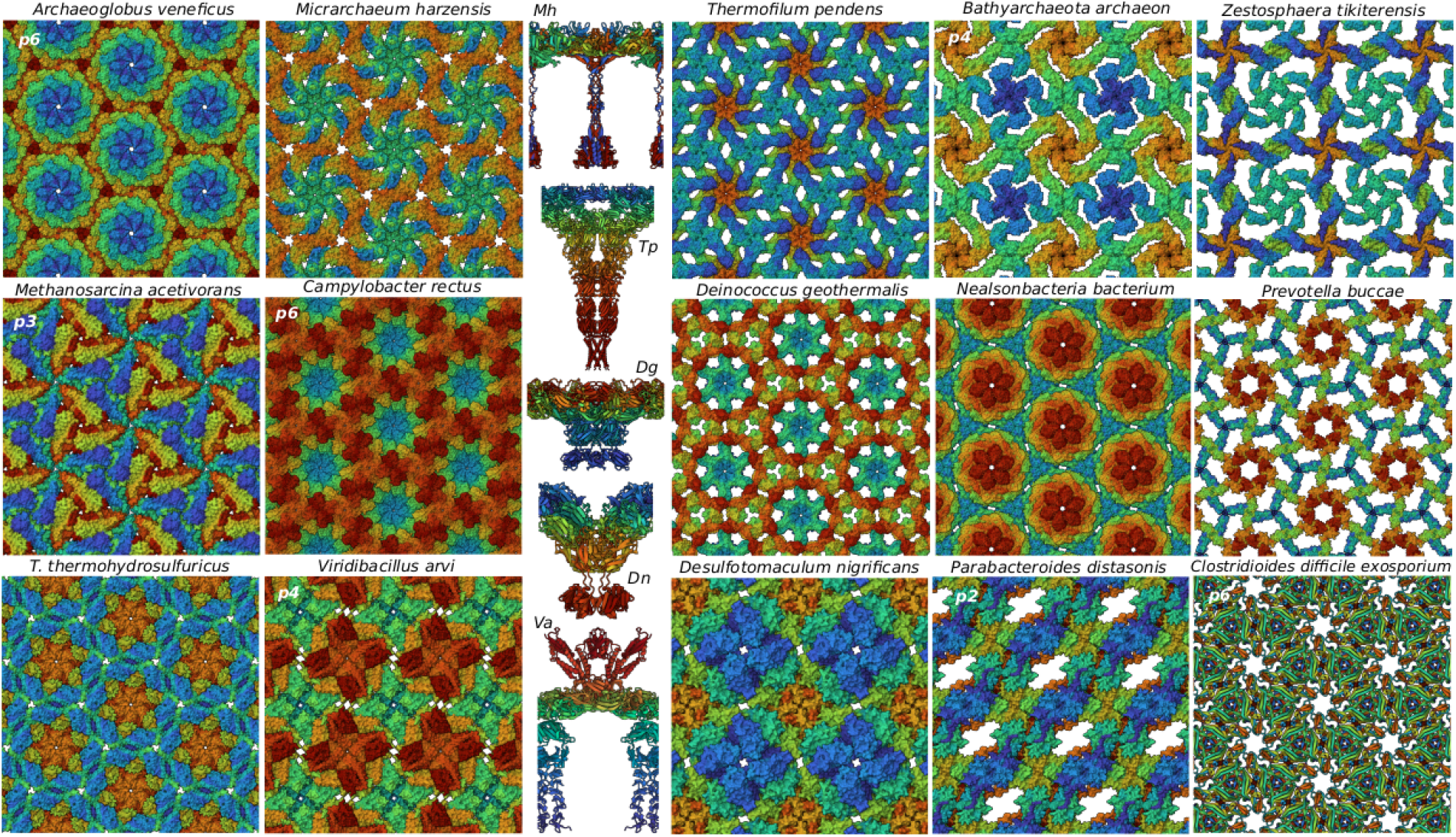
Example 2D protein lattice predictions. The first six are archaeal S-layers; the next eight are bacterial S-layers and the last is a bacterial exosporium crystal. In addition, there are five side views shown of a single S-layer unit cell highlighting the three-dimensionality of the S-layer crystals, including inward-facing cell-anchoring and outward-facing exterior domains. The top views shown are from the exterior of the cell (i.e., looking at the cell from the outside) and the colouring reflects the position along the sequence with the N-terminus dark blue and the C-terminus red.

Successful predictions were then iteratively used as stepping stones into increasingly distant taxonomic regions from the starting set of SLPs (Figs. S5-S6). This approach allowed us to detect SLPs in several largely uncultured taxonomic lineages within Archaea (Figs. S10-16). Among these were species within the phylum Hydrothermarchaeota, which includes hyperthermophilic species discovered in hydrothermal vents, and the class Bathyarchaeia, representing one of the most abundant microbial groups on Earth. Additionally, we found SLPs in the class Methanocella as well as in the phyla Micrarchaeota, Nanoarchaeota, and Nanohaloarchaeota, all of which comprise symbionts. In Bacteria (Figs. S17-S19), we identified SLPs within the mostly uncultured phylum Patescibacteria, which encompasses diverse Gram-negative bacteria characterized by reduced genomes and a symbiotic lifestyle. Furthermore, for organisms known to possess S-layers but lacking candidate proteins with obvious homology to known SLPs, we scanned their genomes for putative SLPs. The search criteria included several relevant target features, such as the presence of signal peptides, potential sequence motifs or domains for cell anchoring, and other characteristic structural features of SLPs (e.g., large size of the protein or the presence of multiple Ig-like domains). This approach facilitated the identification of completely novel classes of SLPs, notably within the bacterial orders Tepidiformales (e.g., *Tepidiforma thermophila*), Paceibacterales (Candidatus Nealsonbacteria bacterium), and Bacteroidales (*Prevotella buccae*), as well as the archaeal order Thermofilales (*Thermofilum pendens*). Additionally, we identified SLPs in taxonomic groups previously unknown to harbor S-layers, including the archaeal phyla Aenigmatarchaeota, Altiarchaeota, and Undinarchaeota, and the bacterial phylum Thermodesulfobiota.

Overall, our analysis has elucidated the structural diversity of S-layers across a wide range of bacterial and archaeal phyla (Figs. 3-4), covering a broad spectrum of the prokaryotic tree of life. Furthermore, our analysis reveals that S-layers are more prevalent and widespread among prokaryotic organisms than previously recognized, extending to lesser-studied groups and environments. Within archaea, S-layers are prevalent in many phyla (Fig. 3), with notable exceptions in certain taxonomic groups such as the genera Methanobrevibacter (e.g., *Methanobrevibacter smithii*), Thermoplasma (*Thermoplasma acidophilum*), and Ignicoccus (*Ignicoccus hospitalis*). The absence of an S-layer in some of these groups is likely due to the presence of alternative cell wall structures; for instance, Methanobrevibacter species have a pseudomurein layer and Ignicoccus species have a double membrane system. Intriguingly, for the Asgardarchaeota phylum, which contains prokaryotic species most closely related to eukaryotes, our method, when applied to both previously predicted putative SLPs (*41*) and those identified through our homology searches, failed to produce a lattice model. This leaves the existence of S-layers in this phylum uncertain. In contrast to archaea, the distribution of S-layers among bacteria is sporadic, with their presence or absence varying significantly even within individual genera. While several major phyla such as Actinomycetota, Bacillota, Bacteroidota, Chloroflexota, Cyanobacteriota, and Pseudomonadota show a substantial occurrence of S-layers, their existence in others, such as Acidobacteriota, remains ambiguous (Fig. 4). Notably, the evolutionarily deep-branching phylum Deinococcota, which includes hyperextremophiles like *D. radiodurans* and *T. thermophilus*, is one of the only bacterial phyla with an extremely widespread presence of S-layers. Furthermore, S-layers are also prevalent in several other deep-branching lineages, including Verrucomicrobiota, Patescibacteria, Chloroflexota, and Cyanobacteriota, suggesting that S-layers may represent an ancestral feature in bacteria.

**Figure 3:**
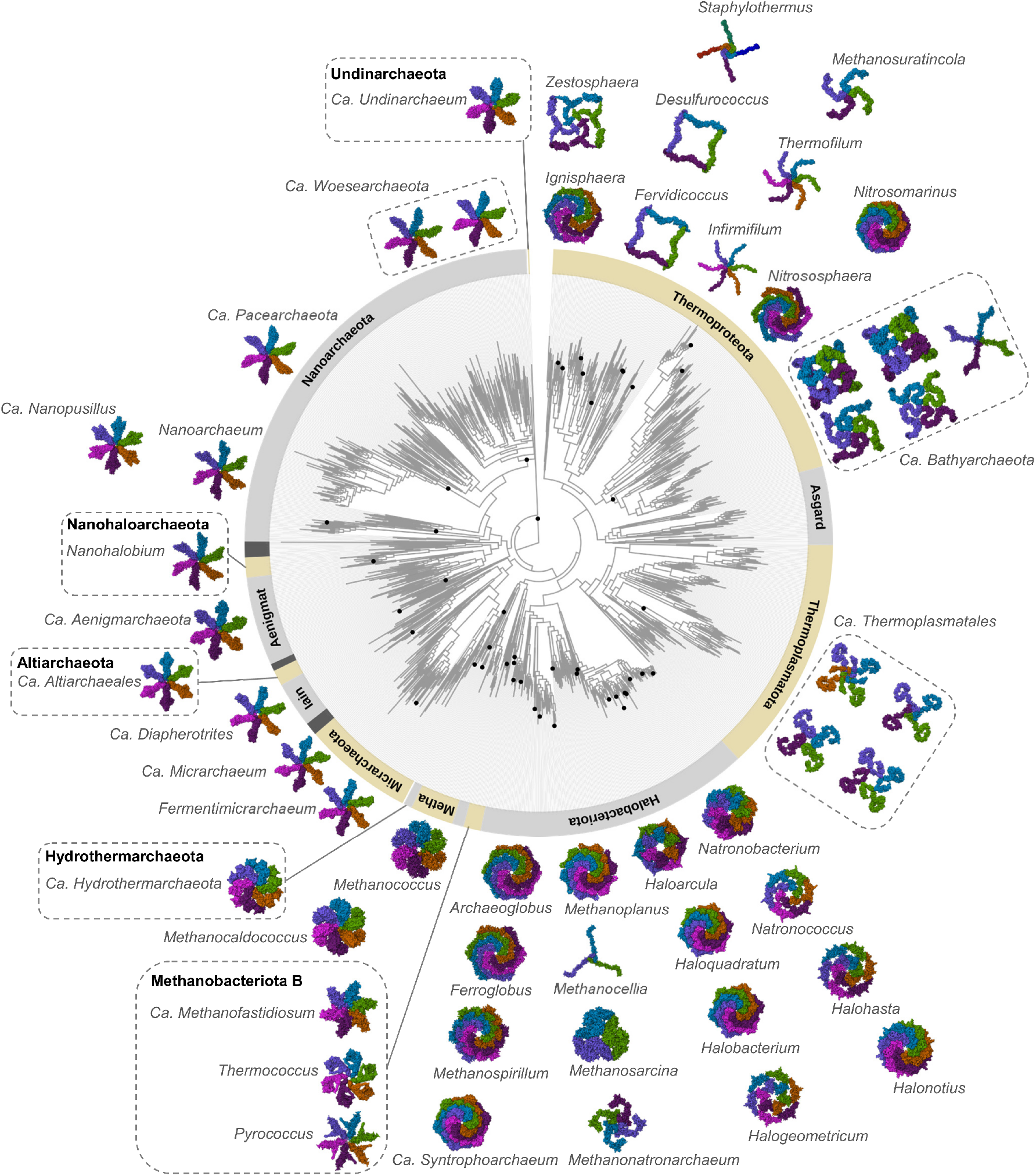
Phylogenetic distribution of S-layer unit cells in Archaea. A phylogenetic tree at the phylum level for Archaea, based on the Genome Taxonomy Database (GTDB) classification. Representative S-layer unit cells are shown and labeled by genus names. The outer ring denotes the designated phyla according to GTDB. Major phyla are annotated, and each dot represents the genus corresponding to the displayed S-layer structures. Where species are not classified within GTDB or the genus designation remains unresolved, assignment to the next higher taxonomic rank is indicated. Abbreviations: ‘Metha’ for Methanobacteriota and ‘Iain’ for Iainarchaeota. Accession details and domain composition data for each S-layer can be found in Table S1 and Figs. 5 and S7.

**Figure 4:**
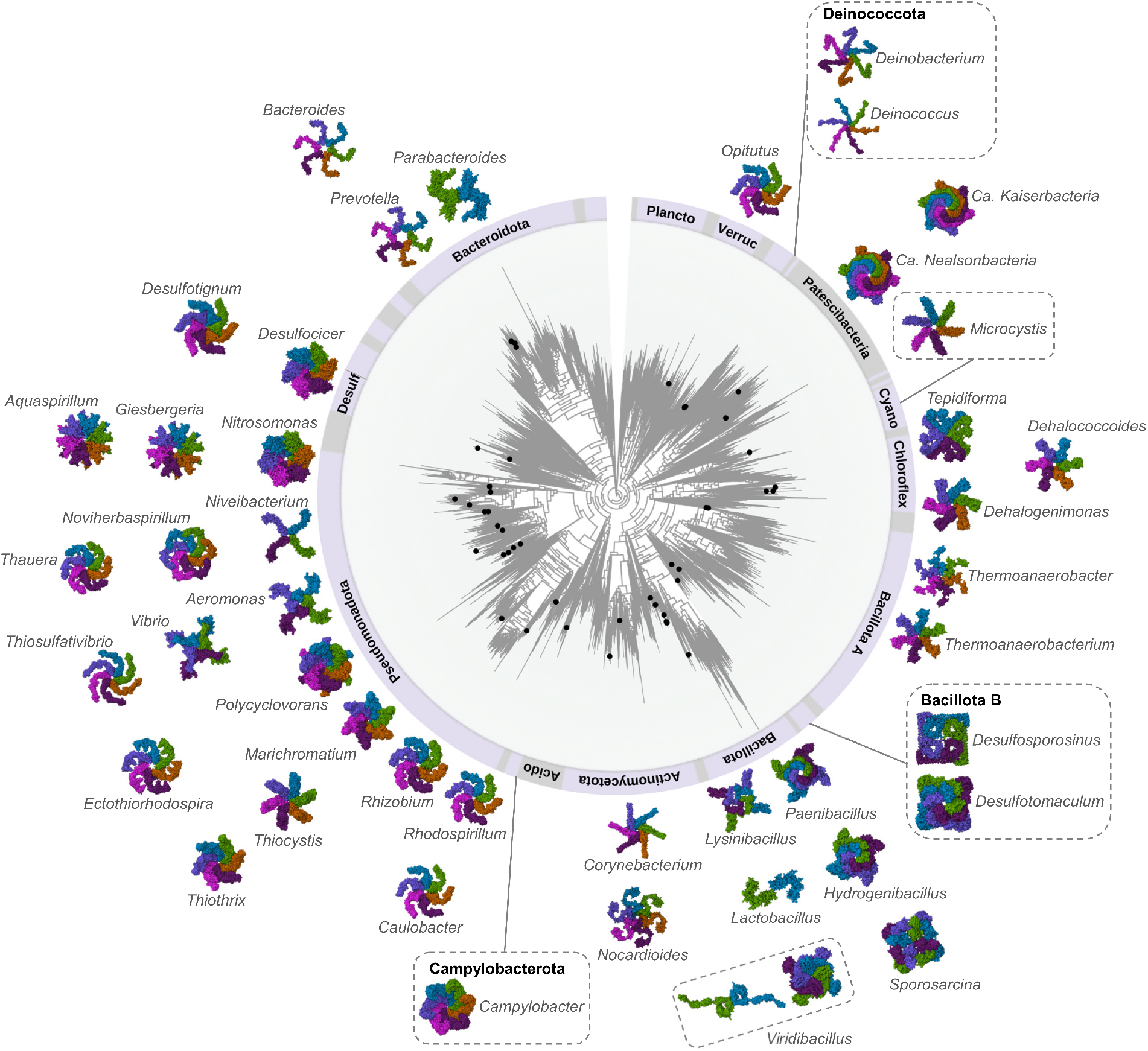
Phylogenetic distribution of S-layer unit cells in Bacteria. Following the same scheme used in Fig. 3, this figure illustrates a GTDB phylogenetic tree for Bacteria at the phylum level. The outer ring denotes the designated phyla and each dot represents the genus corresponding to the displayed S-layer structures. Abbreviations: ‘Plancto’ for Planctomycetota, ‘Verruc’ for Verrucomicrobiota, ‘Cyano’ for Cyanobacteriota, ‘Chloroflex’ for Chloroflexota, ‘Acido’ for Acidobacteriota, and ‘Desulf’ for Desulfobacterota. Accession details and domain composition data for each S-layer can be found in Table S2 and Figs. 5 and S8.

At the lattice level, our analysis shows that S-layers are characterized by astonishing structural diversity (Figs. 2 and S10-S20). Although *p*6 planar symmetry is the most commonly observed symmetry in S-layer lattices, all possible planar symmetries for proteins, including *p*4, *p*3, *p*2, and *p*1, are also observed, occasionally even within a single phylum, such as Bacillota (Fig. 4). The lattice constants of the S-layers range from *∼*37 Å in *Lactobacillus acidophilus* to *∼*270 Å in *Odoribacter splanchnicus*, with S-layer thicknesses ranging from *∼*40 Å in *L. acidophilus* to *∼*300 Å in *Lysinibacillus sphaericus*. At the protein level, SLPs primarily consist of multiple tandemly repeated domains with a single fold type that form the bulk of the 2D sheet. Additionally, they include domains or segments that contribute to the formation of stalks extending from the sheet towards the cell membrane, which are involved in membrane anchoring. Although certain folds, such as the Ig-like and other *β*-sandwich folds, are predominantly observed, a variety of other folds are also present (Figs. 5 and S7-S8). These include the OB-fold seen in *Thermanaerobacter kivui, β*-helix fold in *C. vibroides*, the *α*-helical bundle in *C. glutamicum*, the *β*-trefoil fold in *Parabacteroides distasonis*, and the *β*-propeller in *Dehalococcoides mccartyi*.

**Figure 5:**
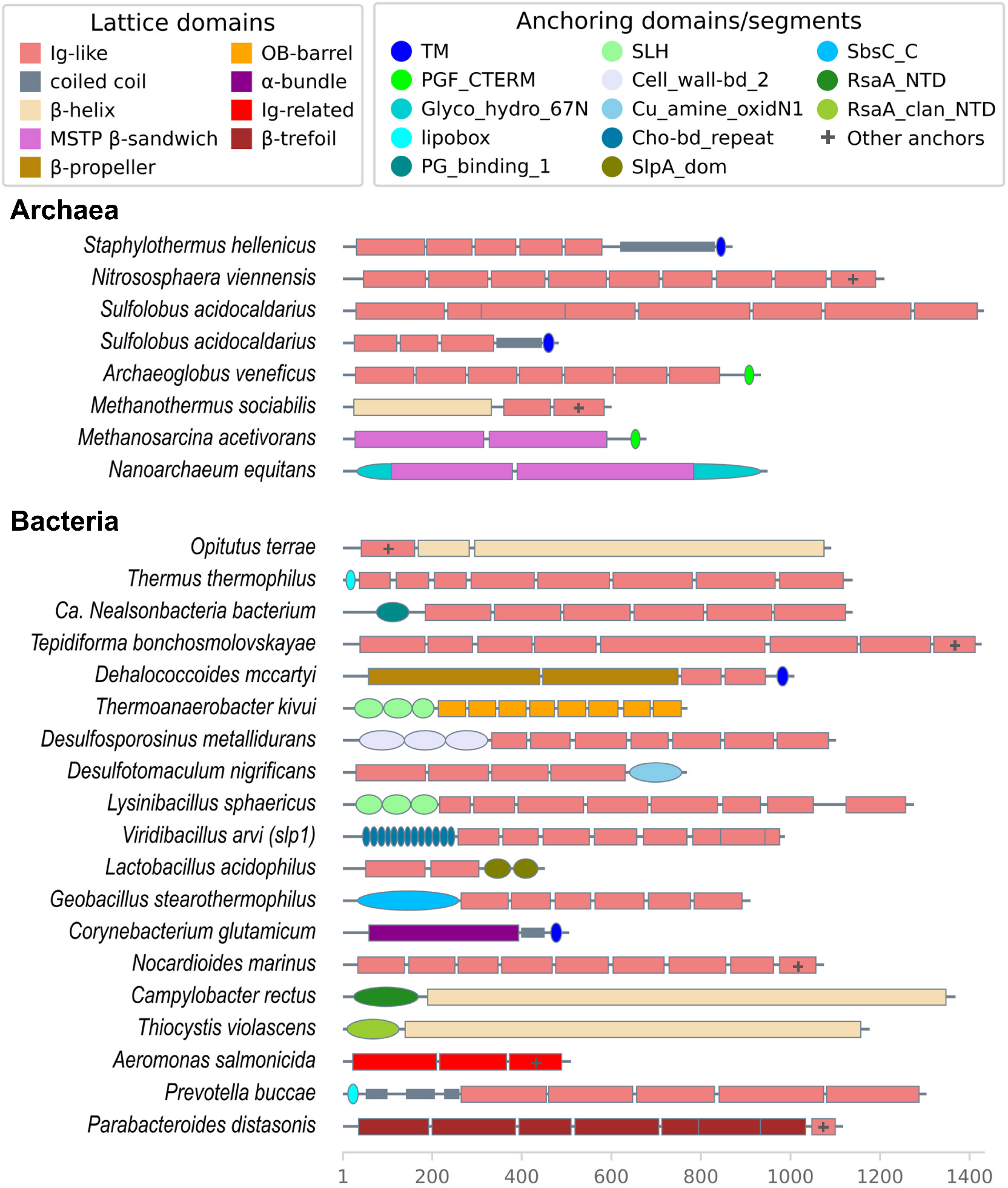
Diversity of SLPs in Archaea and Bacteria. The domain composition and anchoring mechanisms of SLPs in representative archaea and bacteria are shown as cartoon block diagrams. For each organism listed, the domain architecture is represented by colored blocks, with each color corresponding to a specific type of lattice domain or anchoring segment, as defined in the key (top). Putative anchoring domains that share a fold also observed in lattice domains are marked with a ‘+’. Compared to archaea, bacteria display a greater diversity of both lattice and anchoring domains, indicative of their diverse cell surfaces, needed to inhabit diverse environments. The length of the proteins, in amino acids, is indicated by the scale at the bottom. For detailed domain architectures in archaea and bacteria, please refer to Figs. S7-S8.

The taxonomic distribution of SLPs and the resulting S-layer lattices, as illustrated in Figs. 3-4, highlights a notably higher level of structural diversity among bacteria compared to archaea. Within each archaeal phylum, there is a greater conservation in fold composition as well as the overall architecture of S-layers (Fig. 5), despite significant variations in the details of individual S-layers, such as those observed in the phylum Halobacteriota (Fig. 3). Archaeal Slayers can be categorized into three main structural classes. These include the hexagonal spiral pyramids formed of Ig-like domains seen in Halobacteriota; the p4 lattice structure composed of Ig-like domains seen in Thermoproteota, and the hexagonal Methanosarcinales S-layer tile protein (MSTP)-like *β*-sandwich configuration observed in class I methanogen phyla such as Methanobacteriota A and Methanobacteriota B, as well as DPANN superphylum phyla such as Altiarchaeota, Aenigmatarchaeota, Nanoarchaeota, and Micrarchaeota (Fig. 3). Sequence analysis indicates that SLPs displaying the MSTP-like *β*-sandwich fold are orthologous, whereas those containing Ig-like domains demonstrate extreme divergence, making their evolutionary relationships challenging to substantiate beyond a specific phylum (Figs. S5-S6). While species of the phylum Halobacteriota generally possess SLPs with Ig-like domains, species of one of its genera, Methanosarcina, possess SLPs comprising MSTP-like *β*-sandwich domains, distantly homologous to those observed in DPANN and class I methanogen SLPs, suggesting a likely acquisition by horizontal gene transfer (Figs. 5 and S7).

In contrast, the bacterial SLPs and their arrangements in the lattice exhibit substantial vari-ability, even within individual phyla (Fig. 4). For example, in the phyla Pseudomonadota and Bacillota, there is remarkable sequence variability in SLPs, evident in symmetries, arrangements, and structures (Fig. 4). The predominant class of SLPs, conserved in both sequence and structure among bacteria, is the hexagonal lattice structure characterized by a *β*-helix fold, observed across the phyla Pseudomonadota, Campylobacterota, and Desulfobacterota, as well as in the more distant phyla Cyanobacteria and Verrucomicrobiota (Figs. 4 and S8). However, it remains unclear whether the presence of *β*-helix fold SLPs in these distant phyla is due to divergence from a shared ancestral SLP or horizontal gene transfer. Furthermore, while Ig-like domains are also frequently observed in bacterial SLPs, they are extremely divergent at the sequence level, making it challenging to evaluate their evolutionary relationships beyond a specific bacterial taxa or to archaeal SLPs containing Ig-like domains. Collectively, these findings suggest that while archaeal SLPs exhibit a degree of evolutionary conservation, bacterial SLPs likely have had multiple independent evolutionary origins.

This significant difference in the evolution of bacterial and archaeal S-layers is further supported by our bioinformatic prediction of their cell-anchoring segments (Figs. 5 and S7-S8). Our analysis indicates that, compared to archaeal S-layers, bacterial S-layers exhibit remarkable diversity in their cell-anchoring mechanisms, which is congruent with the diversity observed in their lattice structures (Figs. 3-4). In most archaea, S-layers are directly anchored to the cell membrane through lipidation, transmembrane helices, membrane-binding domains, or interactions with co-expressed accessory proteins. In contrast, bacterial S-layers can be anchored to various components of the cell envelope, including the cell membrane, cell wall, or lipopolysaccharide. Specifically, bacterial SLPs employ a variety of anchoring mechanisms, including lipidation (*14*), PG-binding, or secondary cell wall polysaccharide (SCWP)-binding domains (*30, 42*), and transmembrane *α*-helices. These mechanisms in bacteria vary significantly, even within a single phylum, such as in Bacillota and Pseudomonadota, as do their S-layer structures (Fig. 4). Notably, several SLP types in bacteria exhibit a modular architecture, where conserved lattice-forming domains can be paired with diverse anchoring domains, resulting in a variety of S-layers with different anchoring mechanisms. For instance, the *β*-helix domain in Pseudomonadota and Verrucomicrobiota and the Ig-like domain in Bacillota are associated with a range of anchoring domains. A distinctive feature of some S-layers is a relatively long stalk region that separates the lattice from the anchoring segment; for example, the S-layer of the bacterium *C. glutamicum* and many hyperthermophilic archaea within the phylum Thermoproteota. While we identified several known anchoring domains like the SLH domain (Fig. S27), we also discovered many novel ones, including those also found in non-SLP cell-surface proteins, such as the potential C-terminal pseudomurein-binding domain in *Methanothermus fervidus* (Fig. S28).

## Discussion

The ubiquity and relative conservation of S-layers in archaea, contrasted with their sporadic occurrence and enormous diversity in bacteria, have intriguing connotations. One plausible explanation for this difference could be the essential mechanical and protective roles S-layers serve in archaea, where they are a major component of the cell envelope. In bacteria, however, S-layers are a part of a more complex cell envelope architecture that includes peptidoglycan, outer membranes, capsules, and glycocalyces. This suggests a less critical structural role for S-layers, as these additional surface components may provide sufficient capabilities, including protection and adhesion, thereby reducing the reliance on S-layers for survival. This might lead to scenarios where S-layers could be lost or reduced without significantly compromising the bacterium’s fitness. Additionally, the emergence of novel S-layers in certain bacterial lineages indicates a specialization for fulfilling specific niche requirements.

Another factor contributing to the observed diversity of bacterial S-layers could be their evo-lutionary history, which has implications for understanding the evolution of bacteria. A growing body of evidence suggests that the last common ancestor of all bacteria was a diderm bacterium (possessing two membranes), with monoderm bacteria (possessing a single membrane) evolving subsequently through multiple loss events of the outer membrane (*43–45*). These losses are proposed to have occurred in the Terrabacteria group, which includes Bacillota, as indicated by phylogenetic studies (*46, 47*). This suggests that monoderm bacteria within the various Bacillota phyla are not monophyletic and may have independently evolved cell envelope components, including S-layers, contributing to their remarkable diversity.

Cell surface molecules can be regarded as ‘living fossils’, each reflecting the unique environmental conditions an organism has faced throughout its evolutionary history. In light of this perspective, our results provide an intriguing view on the overall evolution of prokaryotes. We observe a comparatively continuous (or “gradual” in the evolutionary sense) diversity in the structure of archaeal cell surfaces, enabling their specialisation in various challenging habitats, including those with high acidity, salinity, and temperature. In stark contrast, the cell surfaces of bacteria show markedly increased variability, likely driven by their colonisation of a wide variety of ecological niches and interactions with a diverse array of species and microenvironments. From a molecular standpoint, this increased variability appears ‘punctuated’ in structural space. Collectively, a key observation across both bacterial and archaeal S-layer lineages is their remarkable structural diversity. This diversity suggests that S-layers did not evolve from a single common ancestor. Instead, it appears that the vast assortment of S-layers has arisen through multiple independent evolutionary events.

More broadly, this study extends the capabilities of protein structure prediction to 2D arrays, which are ubiquitous in biology. We also applied our approach to lattice-forming proteins beyond prokaryotic S-layers, including bacterial exosporia (Fig. 2), bacterial microcompartments, prokaryotic sheath proteins, and viral capsids (Figs. S24-S25). These predictions will be extended further in the future to understand how diverse protein crystals form in various biological contexts.

We anticipate that our findings on the diverse lattice structures of prokaryotic S-layers, characterized by variations in lattice constants, porosity, and thickness, will open new avenues for biomaterial design, especially using the hyperthermostable S-layers of extremophiles. Additionally, our advancements in 2D crystal structure prediction will aid in detection of SLPs in previously unannotated genomes, even in organisms yet to be discovered. The integration of S-layer structure prediction with genomic information will help produce a molecular barcode for each organism based on its cell surface, which would contribute to the understanding of their interactions within multicellular communities and microbiomes, providing opportunities for rational re-design of these communities. The rich dataset we provide in this study will therefore serve as a solid theoretical basis for understanding the cell surfaces of many prokaryotic organisms, offering insights into how prokaryotes fit into their particular ecological niches in the biosphere.

## Supporting information

Supplementary Movie 1: Cryo-ET of the C. glutamicum S-layer

## Acknowledgments

T.A.M.B. would like to thank Andriko von Kügelgen, Ido Caspy, and Suzanne Letham for helpful discussions and to Buzz Baum for critical comments.

## Authors contributions

E.J. and J.P.K.D. developed and applied the workflow for 2D lattice prediction. B.I. and T.A.M.B. performed experiments and cryo-EM structural biology. V.A. performed bioinformatic annotations and deep-homology analyses. J.P.K.D., T.A.M.B. and V.A. wrote the paper with support of all authors.

## Data and materials availability

Please send requests for materials to the corresponding author. All data related to this paper will be deposited in appropriate databases and released at the time of publication.

## References and Notes

1. M. Faini, R. Beck, F. T. Wieland, J. A. Briggs, Trends Cell Biol. 23, 279 (2013).

2. A. Klug, R. E. Franklin, S. P. F. Humphreys-Owen, Biochim. Biophys. Acta 32, 203 (1959).

3. T. A. M. Bharat, A. von Kügelgen, V. Alva, Trends Microbiol. 29, 405 (2021).

4. J. Jumper, et al., Nature 596, 583 (2021).

5. T. J. Beveridge, Curr. Opin. Struct. Biol. 4, 204 (1994).

6. U. B. Sleytr, P. Messner, D. Pum, M. Sara, eds., Crystalline bacterial cell surface proteins (Academic Press, 1996).

7. W. Baumeister, G. Lembcke, J. Bioenerg. Biomembr. 24, 567 (1992).

8. R. P. Fagan, N. F. Fairweather, Nat. Microbiol. 12, 211 (2014).

9. U. B. Sleytr, et al., FEMS Microbiol. Lett. 267, 131 (2007).

10. S.-V. Albers, B. H. Meyer, Nat. Rev. Microbiol. 9, 414 (2011).

11. A. von Kügelgen, V. Alva, T. A. M. Bharat, Cell Rep. 37, 110052 (2021).

12. L. Gambelli, et al., Front. Microbiol. 12, 766527 (2021).

13. T. A. M. Bharat, et al., Nat. Microbiol. 2, 17059 (2017).

14. A. von Kügelgen, et al., Proc. Natl. Acad. Sci. USA 120, e2215808120 (2023).

15. E. Baranova, et al., Nature 487, 119 (2012).

16. A. Fioravanti, et al., Nat. Microbiol. 4, 1805 (2019).

17. A. Sogues, et al., Nat. Commun. 14, 7051 (2023).

18. D. Pum, J. L. Toca-Herrera, U. B. Sleytr, Int. J. Mol. Sci. 14, 2484 (2013).

19. C. J. Comerci, et al., Nat. Commun. 10, 2731 (2019).

20. M. Herdman, et al., Nat. Commun. 15, 3355 (2024).

21. M. Abdul-Halim, et al., mBio 11, e00349 (2020).

22. P. Oatley, J. A. Kirk, S. Ma, S. Jones, R. P. Fagan, Sci. Rep. 10, 14089 (2020).

23. S. F. Koval, K. F. Jarrell, J. Bacteriol. 169, 1298 (1987).

24. M. Sára, U. B. Sleytr, J. Bacteriol. 182, 859 (2000).

25. E. Calabi, N. Fairweather, J. Bacteriol. 184, 3886 (2002).

26. A. Lupas, et al., J. Bacteriol. 176, 1224 (1994).

27. W. Baumeister, S. Volker, U. Santarius, System Appl. Microbiol. 14, 103 (1991).

28. U. B. Sleytr, M. Sára, Z. Küpcü, P. Messner, Arch. Microbiol. 146, 19 (1986).

29. M. A. Arbing, et al., Proc. Natl. Acad. Sci. USA 109, 11812 (2012).

30. A. von Kügelgen, et al., Cell 180, 348 (2020).

31. P. Lanzoni-Mangutchi, et al., Nat. Commun. 13, 970 (2022).

32. L. Gambelli, et al., eLife 13, e84617 (2024).

33. H. Ochner, T. A. M. Bharat, Structure 31, 1297 (2023).

34. R. Evans, et al., bioRxiv p. 2021.10.04.463034.

35. J. P. Roney, S. Ovchinnikov, Phys. Rev. Lett. 129, 238101 (2022).

36. S. Scheuring, et al., Mol. Microbiol. 44, 675 (2002).

37. M. Herdman, et al., Structure 30, 215 (2022).

38. P. Messner, K. Steiner, K. Zarschler, C. Schäffer, Carbohydr. Res. 343, 1934 (2008).

39. K. Meier-Stauffer, et al., Int. J. Syst. Bacteriol. 46, 532 (1996).

40. T. J. Beveridge, et al., FEMS Microbiol. Rev. 20, 99 (1997).

41. H. Imachi, et al., Nature 577, 519 (2020).

42. R. J. Blackler, et al., Nat. Commun. 9, 3120 (2018).

43. J. Witwinowski, et al., Nat. Microbiol. 7, 411 (2022).

44. G. A. Coleman, et al., Science 372 (2021).

45. D. Megrian, N. Taib, J. Witwinowski, C. Beloin, S. Gribaldo, Mol. Microbiol. 113, 659 (2020).

46. L. C. Antunes, et al., Elife 5 (2016).

47. N. Taib, et al., Nat. Ecol. Evol. 4, 1661 (2020).

