## Supplementary Information for "Punctuated and continuous structural diversity of S-layers across the prokaryotic tree of life"

Edward Johnston and Jonathan P. K. Doye
*Physical and Theoretical Chemistry Laboratory, Department of Chemistry, University of Oxford*

Buse Isbilir and Tanmay A. M. Bharat
*Structural Studies Division, MRC Laboratory of Molecular Biology, Cambridge CB2 0QH, United Kingdom*

Vikram Alva
*Department of Protein Evolution, Max Planck Institute for Biology Tübingen, Tübingen 72076, Germany*
(Dated: May 28, 2024)

### S1. METHODS: STRUCTURE PREDICTION

An overview of our approach to predict the structures of S-layers is outlined in Fig. S1. The individual steps are detailed below.

#### A. Alphafold

Central to our approach is the use of AlphaFold-multimer [1] to predict the structures of hypothetical complexes of the S-layer proteins (SLPs). We use the "ColabFold" implementation [2]. AlphaFold takes as its main inputs the sequence of the relevant protein and a multiple-sequence alignment (MSA). Following the default workflow of ColabFold we generally use MMseqs2 [3] to generate the MSA. AlphaFold leverages the structural information contained within this set of evolutionarily-related sequences to direct its search for the most favourable structure of the protein(s) [4]. Occasionally, we use the more computationally demanding JackHMMER algorithm [5] to generate an alternative MSA for certain cases. AlphaFold can also take "templates" from the Protein Data Bank (i.e., experimentally-determined structures for related sequences) as input. We did not use this feature, mainly because in the vast majority of cases there are not likely to be any relevant templates, and, in the few cases where the structure of the S-layers are known, we wanted to test if AlphaFold could locate the structures without this additional help.

For each hypothetical SLP complex, we run predictions for 5 models with up to four different seeds. For larger systems 20 predictions would often not be achieved for the default run time that we used. Based on the results, we would then decide whether to run further calculations to reach that number; specifically, if a very high-confidence prediction was already achieved or we deemed there was little hope of achieving a good prediction, we would not run further predictions. For each system we usually ran predictions for hypothetical SLP complexes with between two to seven copies of the SLP.

#### B. Analysis metrics

AlphaFold computes three principal confidence metrics for its structural predictions, namely predicted local-distance difference test (pLDDT), predicted aligned error (PAE), and predicted template modelling (pTM) score.

Firstly, pLDDT is defined for each amino acid and provides a measure of local accuracy for the protein backbone. It is measured on a scale of 0–100 with high values indicating greater confidence. pLDDT has been shown to align well with the true LDDT score during validation across protein structures that have been deposited in the PDB but not been seen by AlphaFold during training [6]. Low values indicate that that region of the protein is either intrinsically disordered or that AlphaFold is unable to make a confident prediction of the true local structure. The most likely reason for the latter is that the MSA is not of sufficient depth for AlphaFold to extract enough information to guide its search for the correct structure.

We use the value of the pLDDT averaged over all the amino acids in a protein as a metric for the order in AlphaFold's prediction of a monomer. Generally, SLPs are found to be well ordered with intrinsically disordered regions restricted to short sections at the N and C termini, or in flexible linkers between domains. Low average pLDDT for the protein monomer is generally a sign that we will be less likely able to generate a good model of the S-layer lattice from AlphaFold-multimer predictions.

Secondly, PAE provides a measure of confidence in the relative position of two amino acids in units of Angstroms (Å). The value $(x, y)$ represents AlphaFold's confidence in the location of residue $x$ if the predicted structure was aligned to the true structure at residue $y$. The two-dimensional plot of the PAE matrix for the *Haloferax volcanii* S-layer protein is illustrated in Fig. S2(b). The plots allow the identification of a protein's domains and any interactions between them. In this case there are six domains and each domain interacts with the domains adjacent in sequence. Although the inter-domain interactions give this protein a well-defined shape it is not expected to be completely rigid; e.g. the PAE between residues in domain 1 and domain 6 is high. In a similar way the PAE plots for complexes allow the identification

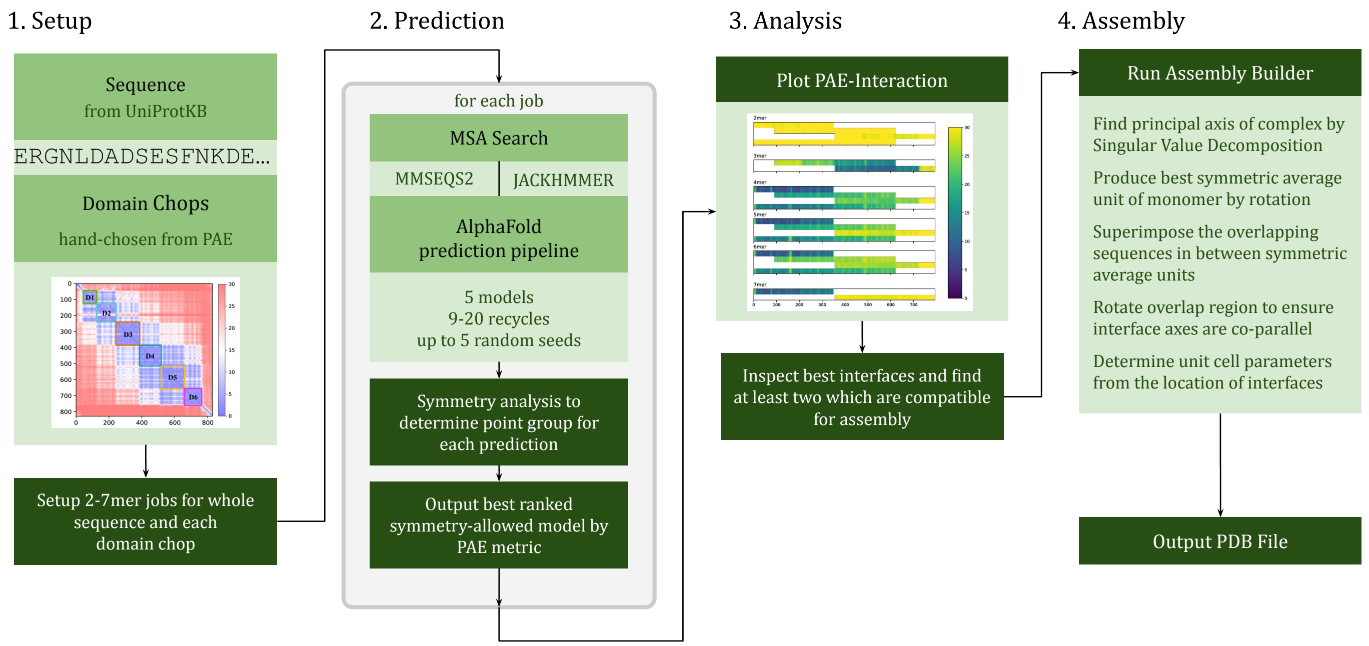


FIG. S1. Diagram summarising our pipeline for predicting the structure of an S-layer lattice.

of inter-protein interactions.

AlphaFold also generates a pTM score as a measure of global accuracy. For Alphafold-multimer predictions this is useful to assess the tendency to form (hypothetical) complexes, since strong inter-protein binding spatially localises the relative positions of the protein monomers in the complex. AlphaFold-multimer also has an interface pTM score (ipTM) that is a modified pTM to consider solely inter-chain interfaces [1]. For a given set of AlphaFold $n$-mer predictions, we generally only considered the complex with the highest pTM that had a symmetry compatible with an S-layer lattice (i.e. we excluded complexes that had dihedral symmetry). It has been shown that pTM provides a robust measure to rank the relative quality of multiple predictions [4].

To compare the relative propensity to form complexes of different sizes, we used the PAE metric to generate the plots introduced in Fig. 1A of the main text. These "PAE-interaction" plots show the average PAE value between a particular amino acid and its equivalent in the other proteins in the complex. These allow the identification of both particularly stable complexes and the parts of the protein involved in the inter-protein interactions. Another example PAE-interaction plot is shown in Fig. S3(a) for *Haloferax volcanii*. In these plots, blue represents low uncertainty in the relative positions of the amino acids and hence a confident multimer prediction.

#### C. Strategy for identifying crystal interfaces

Clearly, AlphaFold-multimer cannot directly make predictions of crystal structures as it can only consider finite complexes with open boundary conditions (i.e. where complexes are surrounded by empty space). To make direct predictions of crystal structures, periodic boundary conditions would be required; this however is not currently available within any of the state-of-the-art protein structure prediction algorithms.

To enable 2D crystal structure prediction, our aim was to use AlphaFold-multimer to generate models of the structure around the rotational symmetry axes in the crystals and then to use these to build up models of the crystal. The 2D space groups relevant to S-layers are $p6$, $p4$, $p3$, $p2$, and $p1$. Because S-layers and other membrane bound crystals have two sides (exterior-facing and interior-facing) that are different, this restricts the space groups to the one-sided plane groups (i.e. those groups without $C_2$ axes in the plane of the translational symmetry) [7]. Furthermore, as individual proteins are chiral, only space groups without mirror planes are allowed. Our basic hypothesis is that $n$-mer complexes have the potential to provide models of the structure around a $C_n$ axis of the crystal.

To identify complexes that can potentially provide models of the structure around a $C_n$ axis in the crystal, we use pTM values and the PAE-interaction plots to help identify the most stable complexes. As noted already, we typically run calculations for complexes with two to seven copies of the protein. However, one might ask why it is informative to make predictions for pentamers and heptamers of individual monomers (of SLPs or other lattice-forming proteins), as periodic crystals may never have $C_5$ or $C_7$ axes. Furthermore, if the plane group of an S-layer is experimentally known, we can also exclude other hypothetical complexes (i.e. complexes with certain rota-

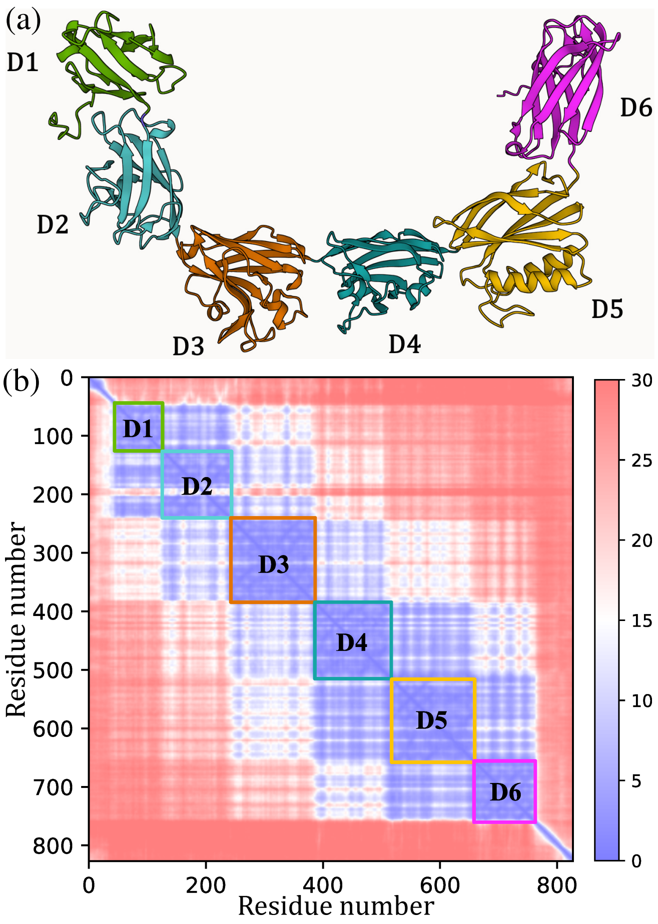


FIG. S2. (a) AlphaFold structure prediction (coloured by domain) and (b) PAE confidence plot for the SLP of *Haloferax volcanii*. Domains are labelled from 1-6 from N- to C- terminus.

tional symmetries) from providing models of the crystal; e.g. tetramers are not relevant to $p6$ crystals as they do not contain any $C_4$ axes. One reason to perform predictions for a complete range of sizes is to provide more confidence in our results. For example, it is expected that the tendency for a $p6$ SLP to form hexameric complexes should be readily apparent from the AlphaFold predictions. If it is not, one should be cautious in using any $C_6$ hexameric predictions in building models of the S-layer lattice or assuming that the SLP has been correctly identified.

Another reason for considering $n$-mers that cannot exhibit crystallographic symmetries (e.g. pentamers and heptamers with $C_5$ or $C_7$ symmetry, respectively) is that, even if they are not relevant to a perfect 2D lattice, they may be relevant to the S-layers that cover the cell. For example, an SLP that predominantly exhibits hexameric interactions, may also be able form pentameric and heptameric "defects" to enable the lattice to better follow the curvature of the surface of the cell [8]. This feature is also seen in capsid proteins of viruses including HIV-1 [9].

As well as considering the complete SLPs (without the proteolytically processed signal peptides), we consider sub-proteins in our analyses. This is for a number of reasons, as outlined below.

Firstly, SLPs (or indeed many lattice forming proteins) are often large and thus the memory requirements of AlphaFold may mean that it is not feasible to run predictions for the full protein for larger $n$-mers. For example, for the *Haloferax volcanii* SLP (called csg containing 827 amino acid residues) on a 48 GB GPU we were only able to run calculations up to $n$=5.

Secondly, and unsurprisingly, given that AlphaFold is essentially performing a search and optimization task, it becomes increasingly hard for AlphaFold to make confident predictions for larger systems. Thus, predictions are potentially easier for complexes of sub-proteins. Also, the MSA for the complete protein can sometimes be dominated by matches to a particular common domain, whereas relevant structural information may be more accessible in an MSA focused on a sub-set of the domains.

Thirdly, SLPs often contain multiple domains and only part of the polypeptide chain may be involved in interactions local to a symmetry site in the crystal. Probably the simplest example of this is when the N-terminal section of the protein is involved in interactions around one symmetry site and the C-terminal section with interactions about a second symmetry axis [10]. For example, this is the case for the *Corynebacterium glutamicum* S-layer. Allowing AlphaFold to focus its attention on the protein structure and interactions local to a symmetry site may thus provide a more efficient approach. Furthermore, the SLP may have domains not involved in the assembly of the 2D crystal. For example, this is often the case for anchoring domains attaching the S-layer to the underlying cell or domains responsible for external interactions that may protrude outwards from the lattice. Anchoring domains can often be identified by homology to known proteins; e.g. SLH (S-layer homology) domains that support binding to peptidoglycan. Another method to identify an anchoring domain is to detect bioinformatically sequences that could be post-translationally modified, e.g. a lipobox lipidation motif on the HPI S-layer protein of *Deinococcus radiodurans*. Experiments that have studied the assembly behaviour of truncated SLPs can also provide useful information on which domains are essential for assembly [11]. These clues are all important in choosing which sub-protein to use in predictions (see also Fig. 5).

Fourthly, it may be necessary to consider sub-proteins to obtain models of a given $C_n$ site. AlphaFold often produces many different structural predictions that optimise the same interface and therefore predictions for the complete protein may provide only a subset of the information required to describe the complete S-layer. For example, the predictions for the trimer of the full *Haloferax volcanii* csg protein are an incomplete hexameric spiral pyramid with $C_1$ symmetry. However, by considering multimers of the final three domains, we were able to ob-

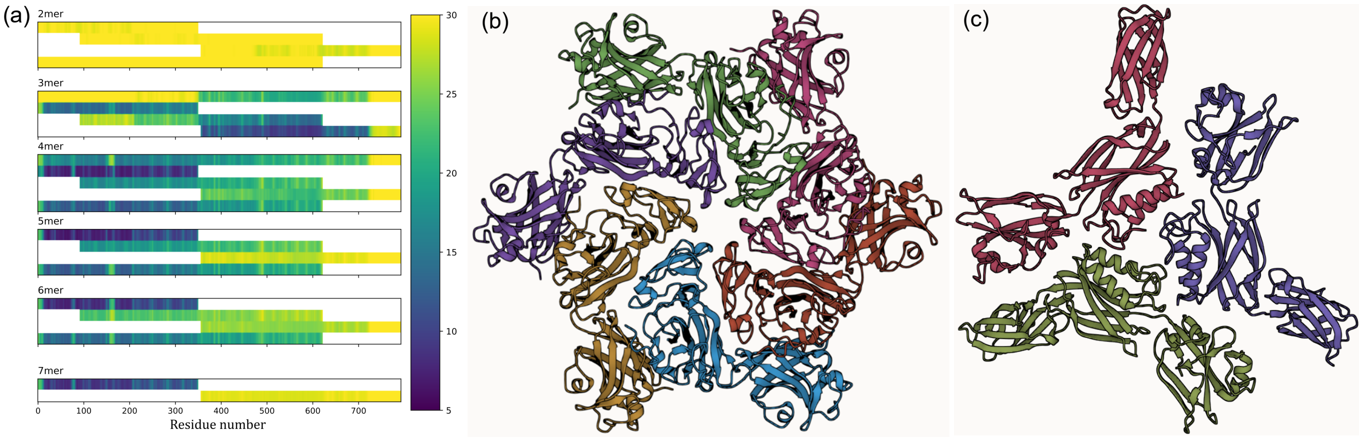


FIG. S3. (a) PAE-interaction plot for the *Haloferax volcanii* SLP. The PAE metric is in units of Å. (b) Highest pTM prediction for a hexamer of the first three domains. (c) Highest pTM prediction for a trimer of the last three domains.

tain a confident prediction of the structure around the $C_3$ site. Furthermore, in $p4$ there are two distinct $C_4$ axes, in $p3$ there are two distinct $C_3$ axes, and in $p2$ there are four distinct $C_2$ axes. In these cases predictions for the relevant $n$-mer might only give information about one of these symmetry sites. For example, the tetrameric predictions for vapA, the SLP for *Aeromonas salmonicida*, always produced a $C_4$ tetramer involving interactions at the N-terminus. To obtain a model of the structure at the other $C_4$ site in the lattice, predictions for tetramers of a sub-protein involving just the two domains nearest the C-terminus were required.

Fifthly, for flexible SLPs, the protein may distort itself to form multiple interactions. Considering sub-proteins allows these interactions to be considered individually. For example, for *Pseudoalteromonas tunicata* the AlphaFold tetramer prediction involves the monomer bending back on itself so that both ends of the SLP form tetrameric interactions.

However, there can be issues with this approach of considering sub-proteins of SLPs. For example, the new surface that is exposed by considering a sub-protein may provide a site for a strong interaction that is not relevant to the S-layer lattice. This is more likely to be a problem when the protein cannot be easily segmented into domains. For example, this can be the case for the SLPs where the assembly domains consists of a large $\beta$-helix. For example, for *Microcystis aeruginosa*, in the dimer predictions of the C-terminus half of the protein, the cut ends interact to re-form a continuous $\beta$-helix. This particular interaction is very strong, giving rise to a "false positive" in the PAE-interaction plot. False positives are also sometimes observed for proteins with low pLDDT, as the relatively disordered structure of the protein monomer can give it more conformational freedom to form strong inter-protein interactions. However, complexes relevant to S-layers typically have both high pLDDT and high pTM.

To illustrate how to interpret the PAE-interaction plots to identify the complexes relevant to the S-layer, we consider that for the SLP of *Haloferax volcanii* (Fig. S3(a)). It it is clear that the N-terminal section of the SLP is able to form confident complexes with 4 to 7 sub-proteins. The rough centering of this window at 6 indicates that this is the preferred hypothetical complex. Note also that the figure shows that the N-terminal domain plays a key role; for example, the confidence of the hexamer is significantly greater for a sub-protein consisting of the first five domains compared to one consisting of domains two to five. This feature is understandable from the structure of the hexameric complexes with the N-terminal domain forming the interactions closest to the $C_6$ axis (Fig. S3(b)). By contrast the C-terminal section shows a clear tendency to form confident trimers. The trimer formed by sub-proteins consisting of domains four to six is shown in Fig. S3(c).

#### D. Building 2D crystals

In those cases where the analysis metrics suggest that we have been able to identify the likely interactions around the symmetry sites in the S-layer lattice, we would then attempt to use these symmetric complexes to build a model of the crystal unit cell. In particular, we construct a model of the asymmetric unit from the most confident complexes with cyclic symmetry by overlapping shared regions of the protein. We then place the combined unit in a unit cell to build a concatenated image of the protein crystal. An overview of the approach is outlined in Fig. S4. For S-layers with $p6$, $p4$, and $p3$ symmetry, models of the structure around only two of the three unique symmetry axes is required, whereas for $p2$ symmetry models representing three of the four unique symmetry axes are required. In the section below, we will initially describe the procedure for the higher symmetry cases.

First, the best asymmetric unit at each interface is de-

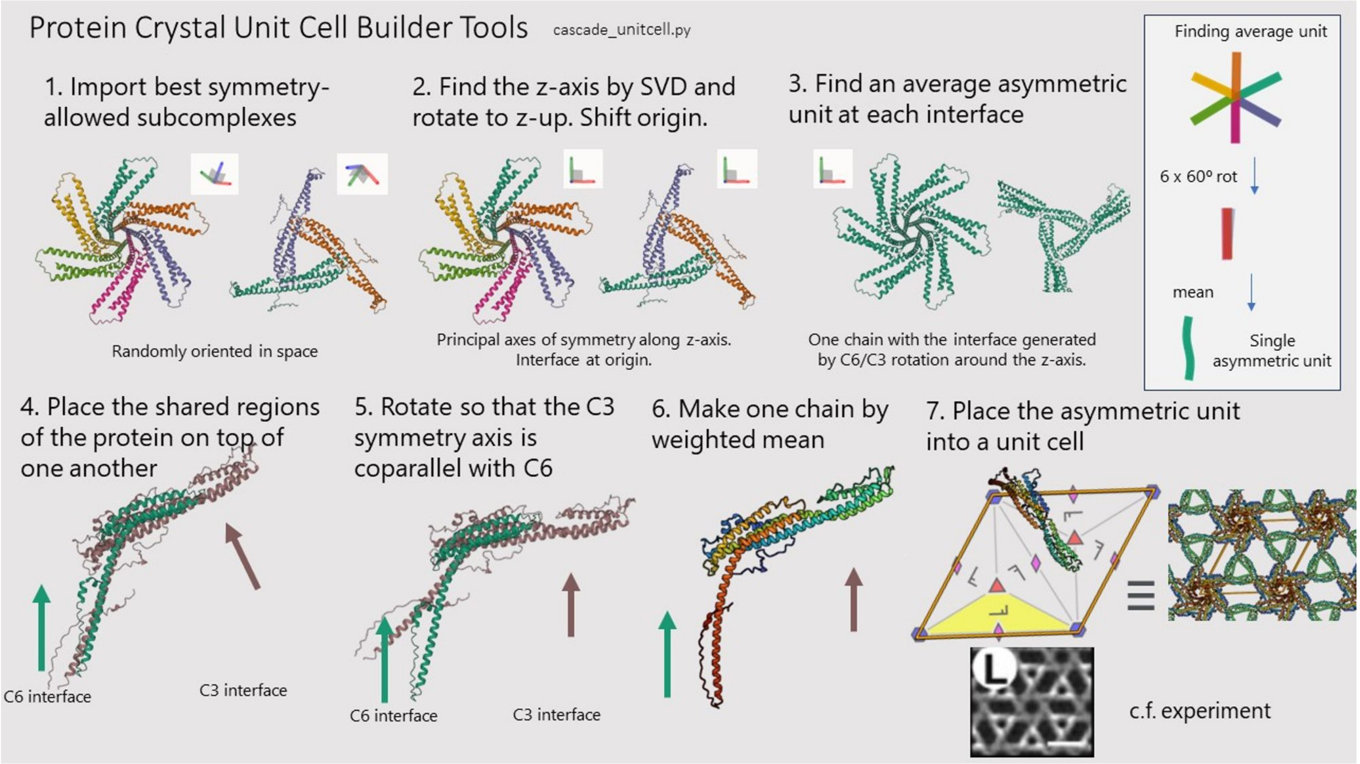


FIG. S4. Diagram summarising the algorithm for building a 2D crystal from AlphaFold-Multimer predictions, illustrated for the *Corynebacterium glutamicum* SLP.

termined. A singular value decomposition (SVD) on the three-dimensional coordinates of $\alpha$-carbon atoms determines the principal moment of inertia axes of the complex relative to its centroid. The symmetry axis is selected by finding the outlier moment of inertia in the case of a symmetric top, or by determining which axis overlays the monomer units on rotation otherwise. The symmetry axis of the complex is then aligned with the $z$-axis in coordinate space by a matrix rotation. Next, a series of rotations are applied in the $xy$-plane overlaying each of the monomer units on top of one another. The mean of these overlapped coordinates is taken to produce the average asymmetric unit about that axis. The root-mean-squared distance (RMSD) between monomers is calculated to ensure the symmetry operation is valid and any chains with RMSD $> 5$ Å are ignored.

In order to predict a 2D crystal structure, it is necessary to calculate the relative positions of the primary and secondary symmetry interfaces. This is achieved by joining two asymmetric units together by overlapping residues shared between predictions or by bridging the two sites with an AlphaFold prediction of the monomer structure between the units. The latter is typically only necessary for particularly large S-layer proteins (e.g. this approach was used for *Nitrosomarinus catalina*).

In the former case the sequences of the subcomplex predictions are used to find the overlapping residues between the two chains. The residues of the secondary interface undergo iterative superimposition on the chain with the primary interface by the Matchmaker method [12]. This moves the secondary interface away from the origin in coordinate space. The RMSD between chains is used as a measure of the quality of the overlap. For the S-layer structure prediction to be taken as a valid model of a 2D crystal, the symmetry axes of the two interfaces must be co-parallel. The chain of the secondary interface is then rotated about the centroid of the overlap region to make its principal axis co-parallel with the $z$-axis. The coordinates of each chain are coalesced by a weighted mean according to their distance from the termini of the overlapping region to ensure a continuous protein bend is observed. We record the angle required to make the two symmetry axes co-parallel as a measure of the distortion applied to the protein. A small angle is typically an indicator that the correct interactions have been identified and that the predicted crystal model is likely to be accurate.

When the bridging interaction approach is used, the same algorithm is used to overlap the primary chain in the subcomplex and the bridge, followed by an overlap with the bridge and the secondary chain. A rotation to make the secondary interface co-parallel occurs afterwards.

The combined asymmetric unit from either method discussed above can then be placed in a unit cell positioning the interfaces at their relevant locations. The unit cell dimensions (determined from the difference in $x$ and $y$ coordinates of the two symmetry axes) and space group are encoded in a PDB file along with remarks labelling the subcomplexes used for construction. The unit cell can then be interpreted by protein viewers to display an entire S-layer in any convenient visualisation software capable of reading PDB files along with the symmetry matrices. We have generally used Mol* for this visualization [13].

In many of the $p6$, $p4$, and $p3$ examples of S-layers, similar to the *Corynebacterium glutamicum* S-layer, there are only significant interactions around two of the three unique symmetry axes in the crystal. However, it is also not uncommon for there to be significant interactions at all three symmetry axes. Often building the crystal from two of the interfaces naturally places the relevant parts of the protein in the correct position to interact at the third symmetry site. However, we can also include information from this tertiary interface by overlapping the relevant sub-complex with the deduced asymmetric unit by the method described above. Furthermore, if the sub-protein in the tertiary complex are relatively unconstrained in their positions (as indicated by the PAE plots) with respect to the primary and secondary complexes, we can apply a rotation in the $xy$-plane to shift those residues to a position nearer to a tertiary interface (this was used, for example, for *Zestosphaera tikiterensis*). In neither case, though, is the lattice constant adjusted as this is set by the distance between the primary and secondary symmetry axes.

In the case of $p2$ S-layers, applying the above procedure to the primary and secondary $C_2$ complexes only defines the periodic repeat along one of the two lattice directions. A tertiary $C_2$ complex is also required to complete the definition of the crystal unit cell. This can be incorporated in the same manner as the above, i.e. by overlapping with the unit derived from the primary and secondary interfaces and then applying a rotation to make the third symmetry axis co-parallel with the first two.

It is important to note, though, that the above methodology does not provide a method for deriving a unit cell in the case of $p1$ symmetry, as there are no direct equivalents of the rotational symmetry axes of the complexes to define the orientation of any complexes with respect to the plane of the S-layer. However, $p1$ S-layers seem comparatively rare and only a few examples have been definitively identified (e.g. SbsB of *Geobacillus stearothermophilus*[14] and EA1 for *Bacillus anthracis* [15]). That being said AlphaFold-multimer predictions can still potentially provide significant insights into the interactions stabilizing these S-layers.

In many cases, this procedure is able to build excellent models of the S-layer lattices from the high-confidence complexes identified by AlphaFold. Good indicators of a successful build include that the rotation angle required to align the symmetry axes of the overlapped complexes is small and that the resulting structure is physically plausible, i.e. it does not result in significant overlaps. For example, where experimental information on the S-layer structure is available, good agreement with experimental data can provide further confidence. This might simply be on the level of the measured lattice constant, but can also be consistency with low-resolution reconstructions of the S-layer lattice.

When the unit-cell building process requires large angular distortions or generates a physically unreasonable structure, the reason may simply be that the identified symmetric complexes may not be those relevant to the S-layer. Indeed, the building process provides a good test of the relevance of the complexes when the indicators are marginal. However, when the complexes are of high confidence and there is a clear preference for oligomers of that size, the reasons for the issues with the crystal-building process are likely to have other origins.

One of the most common major issues is when the protein possesses a flexible inter-domain linkage that is unconstrained by either complex (this lack of constraint is usually evident from the PAE plots). For example, the *Aeromaonas salmonicida* class of SLPs consist of three domains where the N- and C-terminal domains interact at the two four-fold axes in the lattice. However, the link between the second and third domains is relatively weak and flexible. In these cases the success of the crystal-building is dependent on whether the angle between the second and third domains in the AlphaFold-monomer structure happens to be representative of the crystal. In the case of *Aeromaonas salmonicida* it is, and a good S-layer model is obtained that is consistent with experimental data. However, in many of the other examples that we considered in this class it is not, leading to crystal structures with significant overlaps and unrealistic lattice constants. Interestingly for *Aeromaonas salmonicida*, the S-layer has been observed to exhibit two forms associated with inter-domain rotations, giving rise to more open and closed forms of the lattice [16].

Building good crystal models for the largest S-layer proteins can also be difficult for similar reasons to the above, even when good models of the structure of the S-layer local to the symmetry axes are available. For example, if the part of the proteins that bridges between the primary and secondary interfaces is large, then even relatively small deviations of the structure of this section from that in the actual 2D crystal can lead to significant overlaps in the resulting crystal model.

For high-symmetry S-layers where there are interactions at the three unique symmetry axes, although, as noted above, the construction process often leads to a lattice prediction where the protein is in the correct position to interact at the tertiary interface, the positioning can also be off, leading either to some overlaps or the residues being somewhat too far apart to fully interact. In the actual S-layer the interactions at the primary and secondary interfaces must adjust somewhat from their locally preferred geometry to simultaneously optimize the structure of the tertiary interface, but of course such balancing is absent from the AlphaFold-multimer predictions.

The crystal-building process is more likely to lead to lower-quality crystal predictions for the lower-symmetry $p2$ examples. This is probably partly because the dimeric interactions that supply models for the structure of the S-layer lattice at the $C_2$ sites provide less constraints on the overall structure of the protein (than say hexameric or tetrameric interactions) and so local optimization of these interactions are likely to lead to greater deviations between the structure of the protein in the hypothetical complexes and in the S-layer. In addition, the proteins associated with the $p2$ S-layers in Bacillota (the phylum for which there are the most known $p2$ examples) tend to be relatively flexible.

Often SLPs contain exterior-facing domains or cell-anchoring domains that are not directly involved in the interactions stabilizing the S-layer lattice. In particular, the exterior-facing domains are often connected to the rest of the SLP by flexible links or elongated tethers. Including these domains in the S-layer predictions can sometimes lead to overlaps between these domain, as their positions are often not constrained in the complexes used to build the S-layer lattice models. For this reason, our lattice models usually exclude exterior domains unless they have a well-defined geometry with respect to the lattice-assembly domains. We also sometimes do not include the cell-anchoring domain if its inclusion impedes the generation of a good model of the lattice.

Of course, the ideal way to predict the structure of S-layers would be to have a protein structure prediction algorithm that directly incorporated periodic boundary conditions to predict the unit cell. In order to achieve this, the neural networks would also need to optimize the unit cell dimensions and shape, as well as the amino acid positions. Such an algorithm would be particularly beneficial for examples with oblique symmetry. It should also lead to improvements in the predictions where the current building process leads to overlaps or missing interactions. It should also allow increased automation (the current approach requires significant manual intervention; for example in the choice of sub-proteins and in the assessment of the AlphaFold results). However, we note that predictions for a $pn$ lattice would require calculations involving $n$ monomers (the number in the primitive unit cell) and so for higher-symmetry examples that have long S-layer proteins the memory requirements may be prohibitive even if such an algorithm was available.

We will again use the *Haloferax volcanii* SLP to provide an example of the crystal-building process. We used as the primary complex a hexamer of the first five domains and a trimer of the last 3 domains as the secondary complex. Thus, the overlap between the two complexes corresponds to domains four and five. After overlapping the two complexes a rotation of just $3.7^\circ$ was required to align the two symmetry axes. The resulting $p6$ lattice model had a lattice constant of 16.37 nm. This compares to an experimental value of 16.8 nm. The RMSD between our predicted asymmetric unit and that from experiment [8] is just 3.1 Å. This compares to an RMSD of 18.1 Å for the AlphaFold monomer. The protein adapts its shape to optimize the contacts in the lattice. The above quantitative comparisons confirm that our approach has generated an excellent model of the *Haloferax volcanii* S-layer lattice.

#### E. S-layer proteins

We started our survey of SLPs with examples where they had been experimentally identified or where identification was possible from existing experimental data (e.g. high abundance in proteomics or transcriptomics studies) coupled with homology. As the number of such proteins is limited, further putative proteins were identified based on sequence similarity indicative of homology to these known proteins (in particular targeting organisms for which S-layers have been characterized, but the SLP not identified). In these cases, the structural similarity of the predicted AlphaFold monomer was able to provide further supporting evidence of a putative proteins's S-layer character. An SLP must also possess the following two features: firstly, a signal peptide (or equivalent; e.g. serralysins [17, 18]) directing it to be exported to the cell surface; secondly, a means of anchoring to the underlying cell. A number of such anchoring mechanisms have been identified and are often common across a wide range of S-layers (Fig. 5). These mechanisms include a binding domain (e.g. the SLH domain is common amongst S-layers of the phylum Bacillota), a C-terminal alpha-helix that forms a coiled coil the end of which embeds itself into the cell membrane (these are common in the $p4$ S-layers in Thermoproteota), a co-expressed protein responsible for cell-anchoring (e.g. the SlaB protein that is common amongst hexagonal S-layers in Thermoproteota), and a C-terminal sequence that is excised and replaced by a lipid (e.g. the PGF-CTERM protein-sorting motif is the norm for S-layers in Halobacteriota) [19–22]. Of course, the above two features are necessary but not sufficient features required for a protein to be an SLP, as other secreted proteins might also possess these features. Furthermore, not all cell-anchoring mechanisms have been identified; indeed, one of the unforeseen outcomes of the current study has been the identification of a number of new cell-anchoring motifs in bacteria (see Section S4). We should also note that automatic assignments of S-layer character in databases are not always reliable.

A confident prediction of 2D lattice-forming ability thus provides an important new means to identify with a high degree of certainty, a protein as having S-layer character. Somewhat similarly, if for a putative SLP no indicators of S-layer character were identified in structural predictions, without there being an obvious methodological cause (e.g. due to a shallow MSA), then it is reasonable to view the assignment in existing databases with some caution. For example, for *Bacteroides thetaiotaomicron* a protein (Q8A6F7) had been experimentally suggested to be the SLP [23]; however, it evidenced no obvious S-layer character in our AlphaFold calculations and we were subsequently able to identify a suite of proteins in this organism that showed clear 2D lattice-forming ability.

The homologs that were confirmed to have S-layer character were then used as stepping stones to explore even more distant regions of taxonomic space. This approach was particularly successful for archaea due to the general evolutionarily relatedness of their SLPs and allowed us to comprehensively explore the structural space of archaeal S-layers. For example, this allowed us to conclusively identify SLPs for the first time in many archaeal phyla (e.g. Hydrothermarchaeota, Aenigmatarchaeota, Undinarchaeota, Iainarchaeota, and Altiarchaeaota) and classes (e.g. Syntropharchaeia, Methanocellia, Bathyarchaeia, and Methanomethylicia).

For organisms where S-layers had experimentally been observed to be present, but no SLPs could be identified through homology, we also scanned the genomes for putative SLPs. Typical features that we used as potential indicators (as mentioned above) included signal peptides, a mechanism for cell-anchoring, multiple domains in the same polypeptide chain, typical S-layer folds (e.g. Ig-like), plausible monomer structure (as predicted by AlphaFold), and no clear functional assignment in databases. This allowed us to identify further families of new SLPs. These included SLPs for the archaeal order Thermofilales and the bacterial orders Tepidiformales and Bacteroidales (e.g. *B. thetaiotaomicron* and *P. buccae*) and the phylum Patescibacteria.

It is also noteworthy that even if a protein is clearly identified as an SLP by bioinformatics and structural prediction, it does not necessarily mean that the protein will be the main component of the S-layer or that the organism usually has an S-layer. Organisms have been identified that have multiple SLPs. Sometimes they can form structurally distinct S-layers that are expressed under different situations; for example, this has been detailed for *Bacillus anthracis* [24] and *Geobacillus stearothermophilus* [25]. In the same vein *Campylobacter fetus* has a suite of SLPs that is thought to enable antigenic variation [26]. Multiple SLPs have also been shown to be advantageous for protection against viruses [27]. In other instances some of the genes encoding SLPs may not be transcribed in ordinary circumstances. Sometimes the multiple SLPs encoded in a genome are similar in sequence and structure, with one being the major component of the S-layer and the others perhaps being a minor component that can be incorporated into the S-layer in the same way as the major component but with perhaps additional domains that can provide added functionality.

A list of the SLPs that we have studied are listed in Table S1 for archaea and Table S2 for bacteria. The tables include links to their entry in UniProt or NCBI, and, where relevant, references to the experimental evidence for the identification of the S-layer gene and to their structural characterization. Where no experimental reference for the identification of the gene has been included, these have been identified by the methods outlined above. Only those for which the overall evidence for their S-layer character is convincing have been included (except for a couple of bacterial examples noted in the text).

#### F. Determinants of success

The success of the current approach in making confident lattice predictions for over 150 SLPs, as well as being a testament to the power of modern protein structure prediction methods [6], is also likely due to a number of features of SLPs that make them particularly well-suited for this approach. Firstly, S-layers are generally stabilized by strong inter-protein interactions, therefore AlphaFold can often readily identify these interactions, even given the relative long lengths of many SLPs. Secondly, as already noted, the residues stabilizing the interactions around the different symmetry axes in the S-layer are often well-separated along the protein chain, as illustrated in the test case of *C. glutamicum*. A frequently observed feature is that the two ends of the SLP form interactions near different symmetry axes [10]. This is why considering sub-proteins is often a successful strategy to identify those different interactions. Thirdly, the majority of S-layers have high-symmetry (i.e. $p6$ or $p4$ space groups) and we generally found that full crystal predictions are easier to obtain for the higher-symmetry S-layers. This is for a couple of reasons. $p6$, $p4$, and $p3$ crystals have three unique symmetry axes and only a model of the structure around two of these axes are required to build a crystal model. By contrast $p2$ crystals have four unique $C_2$ axes, which means that three different dimeric interactions are required to build a crystal model. Even when the dimeric interactions are identified, it is generally harder to build a good model of the crystal (e.g. without significant overlaps) because the dimeric interactions constrain the overall geometry of the protein less than, say, a hexameric interaction, therefore there is a possibility of larger differences between the structure of the protein in a predicted dimer and in the S-layer.

AlphaFold takes as input an MSA, which is then used by its "evoformer" network to generate an estimate of the matrix of separations between all pairs of amino acid residues. This information often sufficiently narrows the search of the massive configuration space associated with proteins to allow the second "structure module" to build confident predictions of the protein structure [4]. When our approach failed, the predominant reason for failure was simply that the MSAs lacked sufficient depth due to the lack of homologs to a protein in the sequence databases or because of the extreme divergence of the homologs. This lack of MSA depth was often visible in a lack of order in the AlphaFold-prediction of the

TABLE S1. Details of archaeal SLPs studied. The length given is the full-length of the protein and includes, for example, signal peptides that are cut off prior to incorporation in the S-layer prediction. For many species of the phylum Thermoproteota with a hexagonal S-layer, a second protein is responsible for anchoring the S-layer to the cell. The relevant protein is also listed for completeness; predictions were not run for these proteins, because, as has been shown previously, they are invariably predicted to form trimers with a C-terminal coiled coil [28]. The lines separate organisms in different phyla. The majority of the accession IDs are for UniProt, but when a protein is not in UniProt the ID corresponds to that in the NCBI. The space group gives the symmetry suggested based on previous experiments; hex: hexagonal (either $p3$ or $p6$). The prediction column lists the $C_n$ complexes that one can be confident provide a good representation of the S-layer around its $C_n$ site. A "+" is included where these complexes have then been used to build a high-confidence 2D crystal model of the S-layer. Predictions that are of lower confidence but still plausible are bracketed. Occasionally for $p6$ S-layers, a sub-section of the protein exhibits high-confidence predictions for both $C_3$ and $C_2$ complexes, but it is not unambiguous which of the two competing complexes should be used to build an S-layer model; this situation is indicated by '3/2'.

| organism | protein name | Accession ID | length | identification of gene | space group | structural characterization | prediction |
|---|---|---|---|---|---|---|---|
| **Halobacteriota** | | | | | | | |
| *Haloferax volcanii* | csg | P25062 | 827 | [29] | *p*6 | [8, 30] | 6+3 |
| *Haloarcula japonica* | csg | Q9C4B4 | 862 | [31] | *p*6 | [32] | 6+3 |
| *Halobacterium salinarum* | csg | B0R8E4 | 852 | [33] | *p*6 | [34] | 6+3 |
| *Haloquadratum walsbyi* | csg | Q18KV5 | 988 | [35] | *p*6 | [36] | 6+3 |
| *Halorutilus salinus* | | A0A9Q4C3H4 | 1020 | | | | 6+3,2 |
| *Halopenitus malekzadehii* | | A0A1H6J162 | 944 | | | | 6+3,2 |
| *Halohasta litchfieldiae* | | A0A1H6SJD9 | 984 | [37] | | | 6+3/2 |
| *Haloarcula hispanica* | slg2 | G0HV86 | 922 | [38] | | | 6+3/2 |
| *Halogeometricum borinquense* | | E4NUF2 | 939 | [39] | | | 6+3/2 |
| *Natronobacterium gregoryi* | | L0AHY9 | 981 | | | | 6+3/2 |
| *Natronococcus amylolyticus* | | L9X9C3 | 784 | | | | 6+3/2 |
| *Natrarchaeobaculum sulfurireducens* | | A0A346PN77 | 1153 | | | | 6+3/2 |
| *Halonotius aquaticus* | | A0A3A6QEP0 | 1002 | | | | 6+3 |
| *Halobacteriales archaeon* | | A0A7J4UPU5 | 804 | | | | 6+3/2 |
| *Methanoculleus marisnigri* | | A3CSY3 | 773 | [40] | *p*6 | [41] | 6+2 |
| *Methanospirillum hungatei* | | Q2FTS3 | 862 | [39] | *p*6 | [42] | 6+3 |
| *Methanoplanus limicola* | | H1YYH2 | 843 | | *p*6 | [43, 44] | 6+3 |
| *Methanocorpusculum parvum* | | WP_180738245.1 | 766 | | *p*6 | [45] | 6+3 |
| *Methanolacinia paynteri* | | WP_048152302.1 | 872 | | *p*6 | [46] | 6+2 |
| *Methanofollis tationis* | | A0A7K4HPA2 | 805 | | *p*6 | [47] | 6 |
| *Syntrophoarchaeum butanivorans* | | A0A7J2S018 | 789 | | | | 6+3 |
| *Methanolliviera hydrocarbonicum* | | A0A520KVP4 | 508 | | | | 3 |
| *Methanophagales archaeon ANME-1-THS* | | A0A520K040 | 743 | | | | 6+3 |
| *Methanosarcinales archaeon ANME-1 ERB7* | | A0A7G9Z804 | 695 | | | | 6+3 |
| *Archaeoglobus veneficus* | | F2KQ80 | 926 | | *p*6 | [48] | 6+3 |
| *Ferroglobus placidus* | | D3RXL2 | 1036 | | *p*4 | [49] | 6+3 |
| *Methanocella arvoryzae* | | Q0W2X2 | 652 | | hex | [50] | 3+3,3 |
| *Methanocella paludiocola* | | D1Z1E7 | 670 | | | | 3+3,3 |
| *Methanosarcina acetivorans* | | Q8TSG7 | 671 | [51] | | [52] | 3+3 |
| *Methanoperedens sp* | | A0A6A2FZZ2 | 689 | | | | 3+3 |
| *Methanonatronarchaeum thermophilum* | | A0A1Y3GH33 | 1462 | | | | 4+2 |
| *Methanohalarchaeum thermophilum* | | A0A1Q6DSL4 | 1738 | | | | 4+2 |
| **Methanobacterioata** | | | | | | | |
| *Methanothermus fervidus* | slgA | P27373 | 593 | [53] | *p*6 | [54] | (6+2) |
| *Methanothermus sociabilis* | slgA | P27374 | 593 | [53] | | | (6+2) |
| **Methanobacterioata A** | | | | | | | |
| *Methanococcus voltae* | sla | Q50833 | 576 | [55] | *p*6 | [56] | 6+2 |
| *Methanocaldococcus jannaschii* | sla | Q58232 | 558 | [57] | *p*6 | [58] | 6+2 |
| *Methanotorris igneus* | | F6BC80 | 524 | [54] | | | 6+2 |
| **Methanobacterioata B** | | | | | | | |
| *Pyrococcus abyssi* | | Q9V0N3 | 604 | | *p*6 | [59] | 6+2 |
| *Pyrococcus horikoshii* | | A0A832WGR9 | 604 | [60] | | | 6+2 |
| *Thermococcus piezophilus* | | A0A172WII3 | 602 | [61] | | | 6+2 |
| *Methanofastidiosum methylthiophilus* | | A0A150IV63 | 792 | | | | 6+3 |
| *Theionarchaea archaeon DG-70* | | A0A151EZ88 | 789 | | | | 6+3 |
| **Hydrothermarchaeota** | | | | | | | |
| *Hydrothermarchaeota archaeon* | | MDI6654781.1 | 517 | | | | 6+2 |
| *archaeon BMS3Bbin15* | | A0A2H6JUG7 | 533 | | | | 6+2 |
| *Hydrothermarchaeales archaeon* | | MEE8167763.1 | 527 | | | | 6(+)2 |

| | | | | | | | |
|---|---|---|---|---|---|---|---|
| **Thermoplasmatota** | | | | | | | |
| *Picrophilus torridus* | | Q6L2C5 | 1216 | [62] | | | |
| *Picrophilus oshimae* | | A0A8G2L779 | 1138 | | *p*4 | [63] | 4 |
| *Thermoplasmatales archaeon SG8-52-3* | | A0A151ES68 | 1234 | | | | 4+4 |
| *Thermoplasmata archaeon* | | A0A842ZRL3 | 1140 | | | | 4(+)2 |
| *Thermoplasmatales archaeon* | | A0A7C1IMS2 | 1183 | | | | 4+4 |
| *Thermoplasmata archaeon* | | A0A662L5X7 | 1234 | | | | 4(+)2 |
| **Thermoproteota** | | | | | | | |
| *Bathyarchaeota archaeon B24-2* | | A0A2H9I136 | 1055 | | | | 4+2 |
| *Bathyarchaeota archaeon* | | A0A3R7EJK5 | 829 | | | | 4+4 |
| *Bathyarchaeota archaeon* | | A0A419KN45 | 916 | | | | 4+4 |
| *Bathyarchaeota archaeon* | | A0A842SLC1 | 821 | | | | 4+4 |
| *Bathyarchaeota archaeon* | | A0A8T4ZI70 | 973 | | | | 4+4 |
| *Bathyarchaeota archaeon* | | A0A7C0UWI3 | 1397 | | | | 4+4 |
| *Bathyarchaeum tardum* | | WGM90076.1 | 791 | | | | 4,4 |
| *Hecatella orcuttiae* | | WP_309492968.1 | 1132 | | | | 6,3 |
| *Bathyarchaeota archaeon* | | A0A497MH16 | 1168 | | | | 6(+)3 |
| *Bathyarchaeota archaeon* | | A0A7C5ZYT4 | 2022 | | | | 3,6 |
| *Methanosuratincola petrocarbonis* | | A0A7J3UZ30 | 766 | | | | 4+4 |
| *Methanosuratincola subterraneus* | | A0A444L8N0 | 780 | | | | 4+4 |
| *Methanomethylicales archaeon* | | A0A842LTV7 | 1125 | | | | 3 |
| | | A0A842LSR3 | 298 | | | | |
| *Geothermarchaeota archaeon* | | A0A662TE17 | 663 | | | | 4+4,2 |
| *Nitrososphaerota archaeon* | | A0A6B2C2V5 | 754 | | | | 4(+2) |
| *Conexivisphaera calida* | | A0A4P2VKM0 | 699 | | | | 4 |
| *Nitrosomarinus catalina* | | A0A2Z2HI46 | 1525 | | | | 6+3 |
| | | A0A2Z2HJ13 | 312 | | | | |
| *Cenarchaeum symbiosum* | | A0RTY7 | 1680 | | | | 6,3 |
| | | A0RXH0 | 321 | | | | |
| *Nitrosopumilus maritimus* | | A9A4Y9 | 1734 | [64] | hex | [65] | 6,3 |
| | | A9A4Y8 | 334 | | | | |
| *Nitrosotalea devanaterra* | | A0A128A0P9 | 1800 | | | | 6,3 |
| | | A0A128A0Z8 | 291 | | | | |
| *Nitrososphaera viennensis* | | A0A060HS03 | 1202 | [66, 67] | hex | [68] | 6+3 |
| | | A0A060HR06 | 340 | | | | |
| *Nitrososphaera gargensis* | | K0IGS6 | 1151 | | | | 6,3 |
| | | K0IDX9 | 359 | | | | |
| *Nitrosocaldus cavascurensis* | | A0A2K5AP30 | 1242 | [67] | | | 6 |
| | | A0A2K5AP17 | 333 | | | | |
| *Ignisphaera aggregans* | | E0SPF1 | 1111 | | hex | [69] | 6+3 |
| | | E0SPF2 | 460 | | | | |
| | | A0A832EMJ0 | 1191 | | | | 4+2+4 |
| *Desulfurococcus mucosus* | | E8R795 | 904 | | *p*4 | [70] | 4+2 |
| *Fervidicoccus fontis* | | I0A0F7 | 893 | | | | 4+2 |
| *Zestosphaera tikiterensis* | | A0A2R7Y451 | 1065 | | | | 4+2+4 |
| *Staphylothermus marinus* | | Q54436 | 1524 | [71] | *p*4 | [72] | 4(+)2 |
| *Staphylothermus hellenicus* | | D7D997 | 863 | | | | 4+4 |
| *Aeropyrum pernix* | | Q9YEG7 | 1550 | [73] | *p*4 | [74] | |
| *Hyperthermus butylicus* | | A2BLH8 | 1368 | | hex | [75] | |
| | | A2BLH9 | 448 | | | | |
| *Pyrodictium occultum* | | A0A0V8RS45 | 1366 | | hex | [76] | |
| | | A0A0V8RRU2 | 421 | | | | |
| *Metallosphaera sedula* | slaA | A4YHQ8 | 1360 | [77, 78] | hex | [78, 79] | |
| | slaB | A4YHQ9 | 416 | | | | |
| *Acidianus ambivalens* | slaA | B1GT61 | 1016 | [77] | hex | [77] | |
| | slaB | B1GT62 | 511 | | | | |
| *Acidianus brierleyi* | | A0A2U9IFP9 | 1361 | | hex | [80] | |
| | | A0A2U9IFV1 | 446 | | | | |
| *Saccharolobus solfataricus* | slaA | Q980C7 | 1231 | [77] | hex | [21, 81] | |
| | slaB | Q980C6 | 397 | | | | |
| *Sulfolobus acidocaldarius* | slaA | Q4J6E5 | 1424 | | hex | [21, 28, 82–84] | |
| | slaB | Q4J6E6 | 475 | | | | |

| | | | | | | |
|---|---|---|---|---|---|---|
| *Thermofilum pendens* | A1RY64 | 1711 | | *p*6 | [74, 85] | 3+6 |
| *Infirmifilum uzonense* | A0A0F7FI64 | 1723 | | | | 3+6 |
| *Nitrososphaerota archaeon* | NHV98707.1 | 1989 | | | | 3+6 |
| *Caldiarchaeum subterraneum* | E6N5P7 | 2586 | | | | 3(+)6 |
| *Calditenuis fumarioli* | MDJ0274653.1 | 2470 | | | | 3+6 |
| *Nezhaarchaeota archaeon* | MCX8142016.1 | 1374 | | | | 6 |
| *Terraquivivens yellowstonensis* | MCL7394657.1 | 2327 | | | | 3,6 |
| *Wolframiiraptor allenii* | MCL7393743.1 | 2392 | | | | 3+6 |
| *Culexarchaeum yellowstonense* | MCR6623725.1 | 1482 | | | | 6 |
| *Brockarchaeota archaeon* | A0A8J7QJZ2 | 1365 | | | | 6 |
| *Korarchaeum cryptofilum* | B1L757 | 917 | | *p*6 | [86] | |
| **Micrarchaeota** | | | | | | |
| *Micrarchaeum harzensis* | A0A7U3GQJ2 | 978 | [87] | *p*6 | [87] | 6+2 |
| *Micrarchaeota archaeon* | A0A7J2ZEL0 | 873 | | | | 6+2 |
| *Fermentimicrarchaeum limneticum* | A0A7D5XBC7 | 876 | | | | 6+2 |
| *Anstonella stagnisolia* | A0A5E4HZI4 | 931 | | | | 6+2,3 |
| **Nanoarchaeota** | | | | | | |
| *Nanoarchaeum equitans* | Q74MU7 | 941 | [88] | *p*6 | [89] | 6+2 |
| *Nanopusillus sp* | A0A833E5G9 | 842 | | | | 6+2 |
| *Nanobsidianus stetteri* | A0A2T9WQU0 | 988 | | | | 2 |
| *Nanobdella aerobiophila* | A0A915SZM1 | 1000 | | | [78] | 2 |
| *Woesearchaeota archaeon* | A0A7J2NRS4 | 832 | | | | 6+2 |
| *Woesearchaeota archaeon* | A0A3A4UMN2 | 914 | | | | 6+2 |
| *Woesearchaeota archaeon* | A0A2J6GXT8 | 700 | | | | 6+2 |
| *Parvarchaeota archaeon* | A0A519BSJ4 | 887 | | | | 6+2,3 |
| *Pacearchaeota archaeon* | A0A8T4NB04 | 862 | | | | 6+3,2 |
| **Nanohaloarchaeota** | | | | | | |
| *Nanohalobium constans* | A0A5Q0UGZ7 | 1015 | [90] | | | 6+2 |
| *Nanohaloarchaea archaeon* | A0A842K9G1 | 839 | | | | 6+2 |
| *Asbonarchaeaceae archaeon* | MDY6761655.1 | 1069 | | | | 6+2 |
| **Aenigmatarchaeota** | | | | | | |
| *Aenigmarchaeota archaeon* | A0A842Q7F5 | 745 | | | | 6+2 |
| *Aenigmarchaeota archaeon* | A0A8T5PKF4 | 794 | | | | 6+2 |
| **Undinarchaeota** | | | | | | |
| *Undinarchaeum marinum* | A0A832UZT5 | 756 | | | | 6+2 |
| *Naiadarchaeum limnaeum* | A0A832V242 | 769 | | | | 2 |
| **Iainarchaeota** | | | | | | |
| *Diapherotrites archaeon* | A0A2G9N319 | 1039 | | | | 6+2 |
| **Altiarchaeaota** | | | | | | |
| *Altiarchaeales archaeon IMC4* | A0A1D2RG66 | 918 | | | | 6+2 |
| *Altiarchaeum hamiconexum MSI* | A0A098EAS5 | 896 | | | | 6+2 |

monomer. For example, this was a particular problem for the hexagonal S-layers in the archaeal order Sulfolobales, which were relatively common test specimens for early electron microscopy studies. By contrast, organisms that are uncultured and have mainly been studied by metagenomic approaches often had sufficiently good MSAs both for the SLP to be identified and confident predictions made for the S-layer structure, the Bathyarchaeia class being a good example of this.

Still, it is not always easy to identify the reasons for success or failure. For example, for a number of the cyanobacterial examples, e.g. *Microcystis aeruginosa*, the MSAs had large depths, and although it was possible to easily find the hexameric interface, identifying the secondary interface (trimer or dimeric) proved to be elusive. Another interesting example was provided by the *Methanosarcina acetivorans* SLP. Only in 1 of 25 runs was it possible to identify one of the strong trimeric interaction, even though this structure had particularly high confidence. Curiously, when the pseudo-dimeric monomer was cut into two units, it then became much easier for AlphaFold to locate the trimeric interactions.

In cases where the MSA did not quite have sufficient depth to obtain a full S-layer prediction, a productive strategy, in some cases, was to consider a fairly close homolog. For example, although it only proved possible to locate the hexameric interface of the *Thermus thermophilus* S-layer, it was possible to obtain a full prediction for *Deinococcus geothermalis* with the two S-layers likely to be very similar. Another strategy that we occasionally used was to generate the MSA by a different method. For example, we were unable to obtain an S-layer prediction for *Nanoarchaeum equitans* using the MSA generated by MMseqs2; however, using an

TABLE S2. Details of bacterial SLPs studied. The columns are as for Table S1. Space groups in inverted commas indicate that the experimental evidence for the suggested symmetry is not conclusive. obl=oblique (i.e. either $p1$ or $p2$).

| organism | protein name | UniProt ID | length | identification of gene | space group | structural characterization | prediction |
|---|---|---|---|---|---|---|---|
| **Pseudomonadota/Proteobacteria** | | | | | | | |
| *Aeromonas salmonicida* | vapA | P35823 | 502 | [91] | *p*4 | [92] | 4+4 |
| *Aeromonas hydrophila* | ahsA | Q44072 | 472 | [93] | *p*4 | [94, 95] | 4,4 |
| *Azotobacter vinelandii* | | C1DRT1 | 454 | [96] | *p*4 | [97] | 4,4 |
| *Pseudoalteromonas tunicata* | slr4 | A4C8H9 | 579 | [98] | *p*4 | [98] | 4,4 |
| *Vibrio aerogenes* | | A0A1M5ZCF8 | 646 | | | | 4+4 |
| *Thiothrix nivea* | | A0A656HGD9 | 1329 | [99] | *p*6 | [99] | 6+3 |
| *Polycyclovorans sp* | | A0A2E2JAJ0 | 1247 | | | | 6+3 |
| *Thiosulfativibrio zosterae* | | A0A6F8PQN4 | 1066 | | | | 6+2 |
| *Halomonas salicampi* | | A0A7Z0LJX4 | 884 | | | | 6+3 |
| *Ectothiorhodospira mobilis* | | A0A1I4SJH9 | 1002 | | *p*6 | [100] | 6+3 |
| *Marichromatium gracile* | | A0A4R4AJU3 | 777 | | *p*6 | [101] | 6 |
| *Marichromatium purpuratum* | | W0E3L0 | 1148 | | | | 6+3 |
| *Thiocystis violascens* | | I3YA85 | 1169 | | | | 6+3 |
| *Methylotuvimicrobium kenyense* | | A0A543VC43 | 1184 | | | | 6+3,2 |
| *Methylotuvimicrobium alcaliphilum* | | G4T3W6 | 2181 | [102] | *p*6 | [103, 104] | |
| *Thiothrix fructosivorans* | | A0A8B0SNA6 | 512 | | | | 2 |
| *Serratia marcescens* | slaA | Q54455 | 1004 | [105] | | | |
| *Simplicispira metamorpha* | | A0A4R2NFA2 | 1092 | | *p*6 | [106] | 6+3/2 |
| *Simplicispira psychrophila* | | WP_027996328.1 | 843 | | | | 6+3 |
| *Aquaspirillum serpens* | | WP_022654557.1 | 1286 | | *p*6 | [107] | 6 |
| *Thauera selenatis* | sef | G3DVZ6 | 961 | | | | 6+3 |
| *Cupriavidus sp* | | A0A5C8AP55 | 1246 | | | | 6+3 |
| *Nitrosomonas marina* | | A0A1I0CSN4 | 1418 | | | | 6+3 |
| *Noviherbaspirillum saxi* | | A0A3A3FZ63 | 1269 | | | | 6+3 |
| *Pseudoduganella albidiflava* | | A0A411WTW9 | 1239 | | | | 6+3,2 |
| *Giesbergeria anulus* | | A0A1H9L801 | 1272 | | *p*6 | [108] | 6+2 |
| *Aquaspirillum sp. LM1* | | A0A1U9JQH6 | 1311 | | | | 6+2 |
| *Niveibacterium umoris* | | A0A840BL31 | 476 | | | | 4+4 |
| *Pelomonas puraquae* | | A0A254NAE2 | 475 | | | | 4,4 |
| *Delftia acidovorans* | | A0A286J602 | 569 | [109] | *p*4 | [110] | 2 |
| *Paracidovorax citrulli* | npdA | A0A8E2VSX4 | 536 | [111] | | | 2 |
| *Simplicispira psychrophila* | | WP_027995450.1 | 505 | | | | 2 |
| *Lampropedia hyalina* | | A0A1M5A0B1 | 607 | | | | 2 |
| *Caulobacter vibrioides* | rsaA | P35828 | 1026 | [112] | *p*6 | [11] | 6+3 |
| *Rhizobium rhizoryzae* | | A0A7W6LK69 | 827 | | | | 6+2 |
| *Rhodospirillum rubrum* | | Q2RML2 | 830 | | *p*6 | [113] | 6+2 |
| *Labrenzia sp. PHM005* | | A0A4Y6RRS6 | 833 | | | | 6(+)2 |
| *Roseospira visakhapatnamensis* | | A0A7W6REI0 | 1004 | | | | 6+2 |
| *Elstera cyanobacteriorum* | | A0A255XLK5 | 1168 | | | | 6+3 |
| *Pelagibacterales bacterium* | | A0A2E4X8C6 | 1320 | | | | 6 |
| *Robiginitomaculum sp* | | A0A2G2LGS2 | 437 | | | | 4,4 |
| **Desulfobacterota** | | | | | | | |
| *Desulfotignum phosphitoxidans* | | S0G712 | 950 | | | | 6+3 |
| *Desulfocicer vacuolatum* | | A0A1W2BUS2 | 1473 | | | | 6+3 |
| *Desulfamplus magnetovallimortis* | | A0A1W1HD46 | 1403 | | | | 6 |
| *Desulfobacter postgatei* | | I5B1N9 | 1325 | | | | 6+2 |
| *Desulfobacula phenolica* | | A0A1H2DQP8 | 1460 | | | | 6+2 |
| *Desulfobacteraceae bacterium* | | A0A7X1L2I8 | 465 | | | | 4,4 |
| **Campylobacterota** | | | | | | | |
| *Campylobacter rectus* | crs | O30524 | 1361 | [114] | *p*6 | [115] | 6+3 |
| *Halarcobacter mediterraneus* | | A0A4Q1B1B3 | 981 | | | | 6,2 |
| **Bacteroidota** | | | | | | | |
| *Flavobacteriales bacterium* | | A0A2E4RDV3 | 1171 | | | | 6 |
| *Bacteroides thetaiotaomicron* | | Q8A6F7 | 928 | [23] | | [23] | |
| | | Q8A7L2 | 1206 | | | [23] | 6,3+2 |
| | | Q8A7L7 | 1056 | | | [23] | 6,3+2 |
| | | Q8A6Q1 | 1038 | | | [23] | 6+3 |
| *Prevotella buccae* | | E6K6E1 | 1296 | | *p*6 | [116] | 6+2,3 |
| *Odoribacter splanchnicus* | | F9ZC13 | 977 | | | | 6+2,3 |
| *Alistipes onderdonkii* | | A0A1Y3QUG0 | 1025 | | | | 6,3+2 |
| *Parabacteroides distasonis* | | A6LF17 | 1110 | [117] | obl | [117] | 2+2+2 |
| | | A6LF88 | 1151 | [117] | | | 2+2+2 |

| | | | | | | | |
|---|---|---|---|---|---|---|---|
| **Verrucomicrobiota** | | | | | | | |
| *Opitutus terrae* | | B1ZZQ1 | 1083 | | | | 6+3 |
| **Planctomycetota** | | | | | | | |
| *Kuenenia stuttgartiensis* | | Q1PYU4 | 1591 | [118] | *p*6 | [118] | |
| | | A0A6G7GLF5 | 1563 | [119] | | | |
| **Methylomirabilota** | | | | | | | |
| *Methylomirabilis lanthanidiphila* | | A0A564ZIL9 | 316 | [120] | *p*6 | [120] | 2 |
| **Bacillota** | | | | | | | |
| *Aneurinibacillus thermoaerophilus* | satA | Q6TL22 | 759 | [121] | *p*4 | [122] | 4+4 |
| *Paenibacillus polymyxa* | | E0RID9 | 788 | | *p*4 | [123, 124] | 4+4 |
| *Hydrogenibacillus schlegelii* | | A0A947CTK6 | 727 | | *p*4 | [125] | 4+4 |
| *Lysinibacillus sphaericus* | sbpA | Q9RER7 | 1268 | [126] | *p*4 | [127, 128] | 4+4,2 |
| | slpC | P38537 | 1176 | [129] | obl | [130] | 2,2,2 |
| | slp1 | M4N8T6 | 1104 | [131] | *p*4 | [131] | 4+4 |
| *Lysinibacillus sp. B2A1* | | A0A2S0JH17 | 1353 | | *p*4 | [132] | 4 |
| *Sporosarcina ureae* | sslA | Q0VJW4 | 1097 | [133] | *p*4 | [134] | 4+4+2 |
| *Sporosarcina psychrophila* | | A0A127W311 | 1224 | | *p*4 | [132] | 4,4 |
| *Alkalihalophilus marmarensis* | | U6SLL1 | 919 | | | | 4+4 |
| *Viridibacillus arvi* | slp1 | A0A0K2Z0V7 | 1016 | [135] | *p*4 | [135] | 4+4 |
| | slp2 | A0A0K2Z2U9 | 983 | [135] | | [135] | 2+2+2 |
| *Brevibacillus brevis* | MWP | P06546 | 1053 | [136] | *p*6 | [137, 138] | 6(+)3 |
| *Lactobacillus acidophilus* | slpA | P35829 | 444 | [139] | *p*2 | [140, 141] | 2+2+2 |
| *Geobacillus stearothermophilus* | sbsA | P35825 | 1228 | [142] | ‘*p*6’ | [143, 144] | |
| | sbsB | Q45664 | 920 | | *p*1 | [14] | |
| | sbsC | O68840 | 1099 | [145] | obl | [146, 147] | (4),(4) |
| | sgsE | Q8VTF1 | 903 | [148] | obl | [25] | 2+2+2 |
| *Bacillus anthracis* | Sap | A0A6L7H9D8 | 814 | [149] | *p*2 | [24, 150] | 2 |
| | EA1 | P94217 | 862 | [151] | *p*1 | [15, 24, 152] | |
| *Paenibacillus larvae* | splA | I0BWH5 | 1007 | [153] | obl | [153] | |
| **Bacillota B** | | | | | | | |
| *Desulfotomaculum nigrificans* | | F6B5K7 | 761 | | *p*4 | [143] | 4+4 |
| *Moorella glycerini* | | A0A6I5ZP87 | 740 | | *p*4 | [154] | 4+4 |
| *Desulfosporosinus metallidurans* | | A0A1Q8QZW0 | 1094 | | | | 4+4 |
| *Dehalobacter restrictus* | | A0A857DL58 | 1072 | | *p*6 | [155] | 4(+)4 |
| *Desulfosporosinus lacus* | | A0A1M5RX17 | 1082 | | | | 6(+)3 |
| **Bacillota A** | | | | | | | |
| *Clostridium aceticum* | | A0A0D8IAA2 | 786 | | *p*4 | [156] | 4+4 |
| *Thermoanaerobacterium thermosaccharolyticum* | | D9TM53 | 767 | | *p*4 | [157–159] | 4+4 |
| *Eubacterium yurii* | | SKC38145.1 | 841 | | *p*4 | [160] | 4+4 |
| *Thermoanaerobacter thermohydrosulfuricus* | sttA | C0L3L9 | 731 | [161] | *p*6 | [157, 162] | 6+3 |
| *Caldanaerobacter subterraneus* | | U5CTJ9 | 726 | | *p*6 | | 6 |
| *Thermobrachium celere* | | R7RMI1 | 1309 | | | | 4,4 |
| *Thermoanaerobacter kivui* | | P22258 | 762 | [163] | *p*6 | [164] | 6+2,3 |
| *Thermoanaerobacterium saccharolyticum* | | I3VS96 | 845 | [165] | | | 6(+)2 |
| *Acetivibrio thermocellus* | slpA | O86999 | 1036 | [166] | obl | [166] | 2,2,2 |
| *Gracilibacter sp. BRH_c7a* | | A0A101VNW3 | 1070 | | | | 2,2,2 |
| **Actinomycetota** | | | | | | | |
| *Corynebacterium glutamicum* | PS2 | Q6QUT4 | 490 | [167] | *p*6 | [168] | 6+3 |
| | PS2 | Q6QUU5 | 498 | [169] | *p*6 | [169] | 6+3 |
| *Corynebacterium aurimucosum* | | A0A553FT37 | 644 | | | | 6+3 |
| *Nocardioides marinus* | | A0A7Y9YF59 | 1067 | | | | 6+3 |
| **Deinococcota** | | | | | | | |
| *Deinococcus radiodurans* | hpi | P56867 | 948 | [170] | *p*6 | [22] | 6 |
| *Deinobacterium chartae* | | A0A841HX00 | 991 | | | | 6(+)2 |
| *Deinococcus geothermalis* | | Q1IY84 | 1050 | | | | 6+3 |
| *Thermus thermophilus* | | A0A7R7YJ88 | 1131 | | *p*6 | [171] | 6 |
| *Oceanithermus profundus* | | E4U6V1 | 954 | | | | 6,3 |
| **Chloroflexota** | | | | | | | |
| *Dehalococcoides mccartyi* | | Q3Z6N3 | 1001 | [172] | *p*6 | [173] | 6+3 |
| *Dehalogenimonas etheniformans* | | A0A2P5P6T6 | 1024 | | *p*6 | [173] | 6+3 |
| *Tepidiforma bonchosmolovskayae* | | A0A5J6SYT4 | 1419 | | *p*4 | [174] | 4+4+4 |
| *Tepidiforma thermophila* | | A0A2A9HDU3 | 1296 | | *p*4 | [174] | 4(+)4+4 |
| *Dehalococcoidia bacterium* | | A0A7Y5SD43 | 1230 | | | | 4+4+4 |

| | | | | | | | |
|---|---|---|---|---|---|---|---|
| **Cyanobacteriota** | | | | | | | |
| *Microcystis aeruginosa* | | A8YC53 | 697 | [175] | p6 | [175] | 6 |
| *Parasynechococcus marenigrum* | swmA | Q7UA16 | 835 | [176] | | [176] | |
| *Synechocystis* sp. Strain PCC 6803 | | P73817 | 1741 | [177] | *p*6 | [177] | 6 |
| *Prochlorococcus marinus XMU1408* | | A0A318R0N2 | 1213 | | | | 6 |
| *Microcystis wesenbergii* | | A0A3E0M907 | 813 | | | | 6 |
| *Microcystis aeruginosa NIES-2549* | | A0A0F6RJK9 | 813 | | | | 6 |
| *Dolichospermum circinale* | | WP_271804994.1 | 1121 | | | | 6 |
| *Anabaena sp. MDT14b* | | A0A1B7WTE9 | 1326 | | | | 6 |
| *Cyanobium sp. NAT70* | | A0A2E0AMV9 | 518 | | | | 4+4 |
| **Patescibacteria** | | | | | | | |
| *Nealsonbacteria bacterium* | | A0A1G2EN61 | 1131 | | | | 6+3 |
| *Doudnabacteria bacterium* | | A0A1F5NRV7 | 1264 | | | | 6+2,3 |
| *Kaiserbacteria bacterium* | | A0A0G1WWP5 | 1173 | | | | 6+2,3 |
| *Kazan bacterium* | | A0A1F4NSF7 | 1274 | | | | 6+3,2 |
| *Parcubacteria bacterium* | | GMX59286.1 | 1185 | [178] | | | 6+3 |
| *Nealsonbacteria bacterium DGGOD1a* | | UMX47672.1 | 727 | | | [179] | 6+2 |
| *Yanofskybacteria bacterium* | | GMX57981.1 | 692 | [178] | | | 6+2 |
| *Vogelbacteria bacterium* | | A0A1G2QP48 | 643 | | | | 6+2 |

MSA generated by the more computationally expensive JackHMMER approach allowed a successful S-layer prediction.

As noted already, most SLPs are multi-domain proteins with different domains interacting at the different symmetry sites in the lattice. However, in a few examples the SLP is a single-domain protein. In these cases we found it much harder to identify the second interface that stabilized the lattice, as it was not possible in these cases to divide the SLP into a sub-protein that could only form the secondary interface. Two such examples are the SLP of *Delftia acidovorans* (and homologs) and *Methylomirabilis lanthanidiphila*. Both form strong dimers, but it was not possible to locate the interactions by which these dimers further assemble into the observed $p4$ and $p6$ lattices, respectively. One might think that by studying an octamer of the *Delftia acidovorans* protein it might form a tetramer of dimers that mimics the S-layer structure around the $C_4$ lattice site. However, in such cases, AlphaFold is generally more likely to predict a three-dimensional assembly rather than recapitulate the two-dimensional growth typical of an S-layer. The reason for this behaviour is not fully clear but it may be because it allows the formation of more interactions, albeit perhaps individually weaker than the secondary interactions in the S-layer. Occasionally we do however see two-dimensional growth. For example, a hexamer of the *Lactobacillus acidophilus* SLP allowed the identification of the identification of the S-layer structure around a second $C_2$ site in its $p2$ lattice.

#### G. Methods: Bioinformatics

To obtain taxonomic information on the prokaryotic species encountered in this study, we utilized the Genome Taxonomy Database (GTDB) [180]. We downloaded the archaeal and bacterial trees (release R214) from GTDB and visualized them using iTOL [181]. To enable this, we used the convert_to_itol module from GTDB-tk v2.1.1 to convert GTDB trees into a format compatible with iTOL.

To investigate the sequence diversity of archaeal and bacterial SLPs, we employed cluster analysis and constructed separate cluster maps for the two groups. Sequences were gathered by querying the UniProt database [182] for homologs of representative archaeal and bacterial SLPs using BLAST [183]. The search parameters were set with an E-value cutoff of 1e-06 and a maximum of 1000 target sequences. We pooled the full-length sequences of bacterial and archaeal SLPs and excluded those labeled as ‘Fragment’. Both bacterial and archaeal datasets were then filtered in MMseqs2 [3] to maintain a maximum pairwise identity of 70% with a length coverage of at least 70%. The filtered sets were subsequently clustered separately in CLANS [184, 185], utilizing the strength of all-against-all BLAST P-values. Clustering was carried out until equilibrium was achieved in a 2D space, with a P-value cutoff of 1e-10 and using the default settings of CLANS (Figs. S5 and S6).

To analyze the domain architecture of SLPs, we employed InterPro [186] as well as HHpred [187] searches with default settings over the PDB70 and ECOD70 databases. These databases are curated versions of the PDB [188] and ECOD [189], respectively, filtered for a maximum pairwise identity of 70%. We also extended our HHpred analysis to the Pfam database [190] and profile HMM databases representative of archaeal and bacterial proteomes [185]. To identify homologs of individual lattice and anchoring domains, we supplemented HHpred searches with BLAST searches over the UniProt database. Additionally, signal peptides were predicted using SignalP 6.0 [191]. These domain architectures are reported in Figs. S7 and S8.

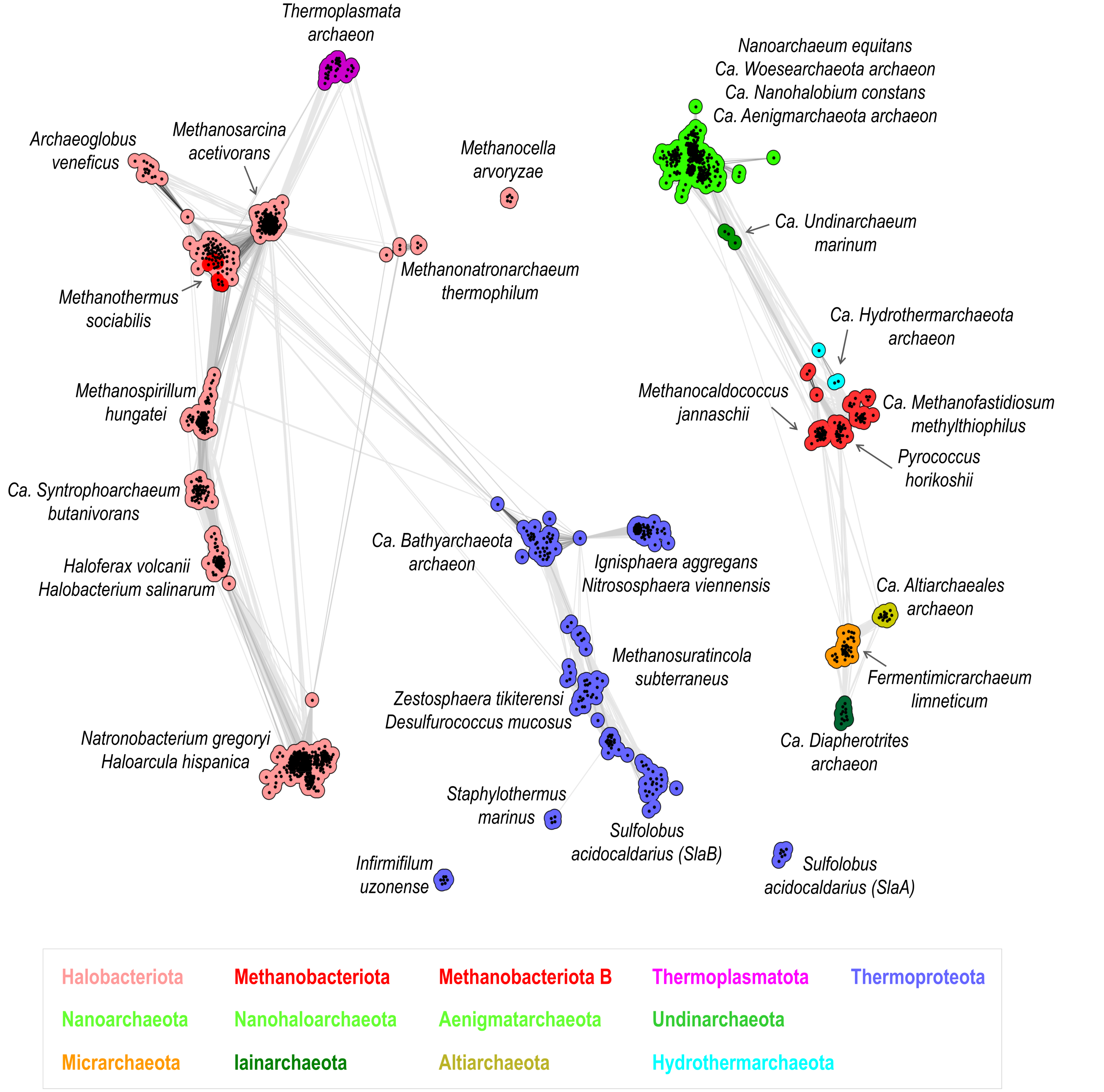


FIG. S5. Cluster map of archaeal SLPs. Homologs of representative archaeal SLPs were collected from the UniProt database using BLAST and were subsequently clustered in CLANS based on the strength of their all-against-all BLAST P-values. Each protein sequence in the map is depicted as a dot, with sequences belonging to the same phylum denoted by the same color. The intensity of the line color reflects the significance of sequence similarities; darker lines indicate higher significance.

### S2. EXPERIMENTAL MATERIALS AND METHODS

#### A. Purification of PS2 S-layer

*Corynebacterium glutamicum* 541 ATCC13058 cells were grown in beef extract-peptone medium at 30°C. Cells were harvested in late-log phase by centrifugation at 4000 rcf (relative centrifugal force) and stored at −80°C until further use. A one litre culture cell pellet was resuspended with 80 mL lysis buffer (50 mM HEPES/NaOH pH=7.5, 150 mM NaCl, 1 mM MgCl2, 50 µg/mL DNaseI, 0.2 mM TCEP (tris(2-carboxyethyl)phosphine), 2 cOmplete Protease Inhibitor tablets (Roche), 2% SDS (sodium dodecyl sulphate) and incubated at room temperature (21°C) for 25 minutes on a rotating wheel. Cell debris was pelleted at 3000 rcf for 15 minutes at 20 °C and the supernatant was spun down at 20,000 rcf for

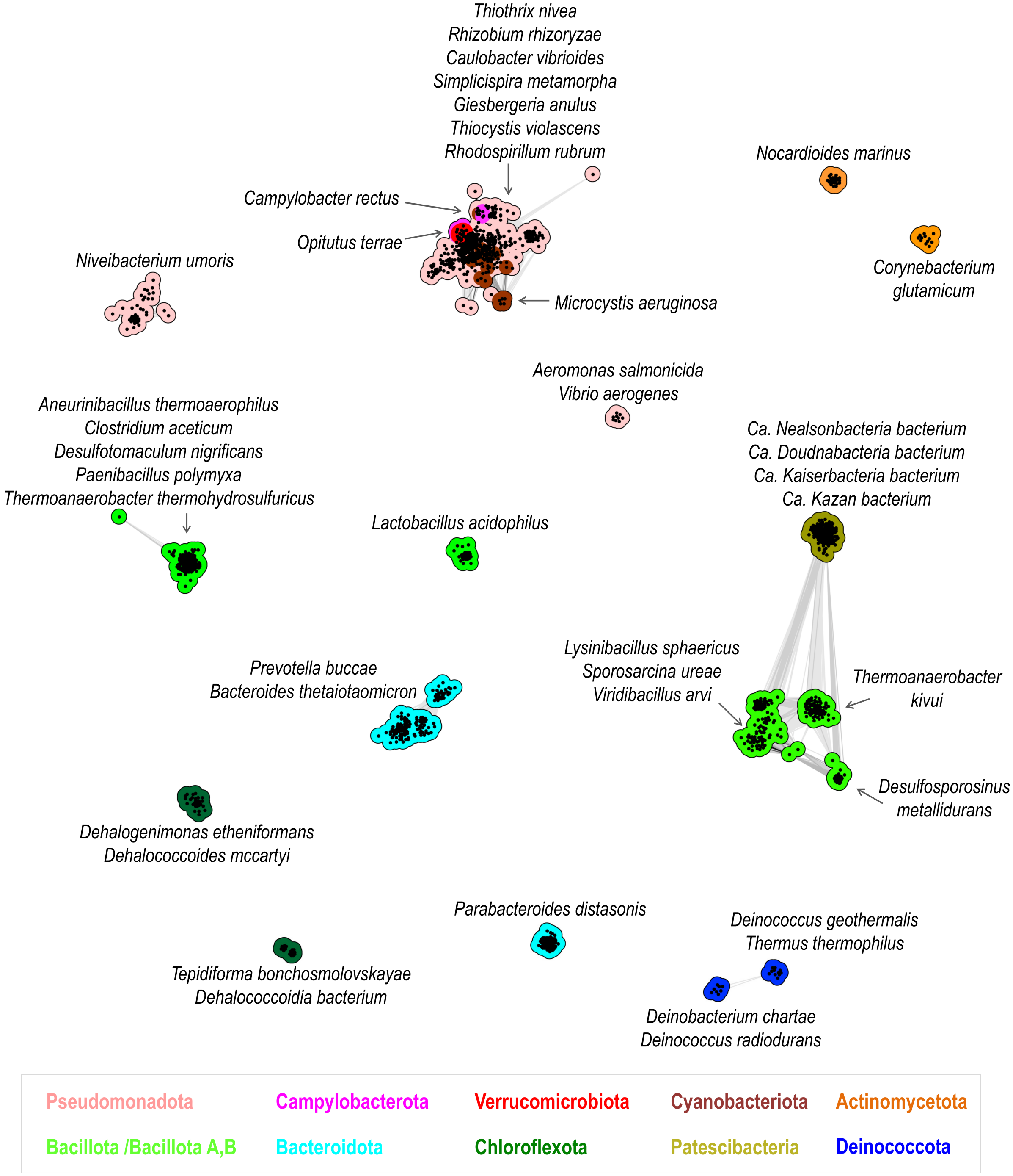


FIG. S6. Cluster map of bacterial SLPs. Compared to archaeal SLPs, bacterial SLPs exhibit substantially greater sequence diversity.

30 minutes at 20 °C. Pellet was washed with 1 mL wash buffer (50 mM HEPES/NaOH pH=7.5, 150 mM NaCl, 1 mM $MgCl_2$, 1 x cOmplete Protease Inhibitor) and spun down at 20 000 rcf for 15 minutes at 4 °C. The final pellet was resuspended in 120 µL of resuspension buffer (50 mM HEPES/NaOH pH=7.5, 150 mM NaCl, 1 mM MgCl2) for cryo-EM grid preparation.

#### B. Cryo-EM grid preparation

Cryo-EM grids were prepared by adapting a previously established workflow for S-layers [22]. Briefly, 2.5 µl of resuspended sample was applied to a freshly glow discharged Quantifoil R3.5/1 Cu/Rh 200 mesh grids and plunge-frozen using Vitrobot Mark IV with −3 blot force and 2.5 seconds blot time. For cryo-ET, the sample was

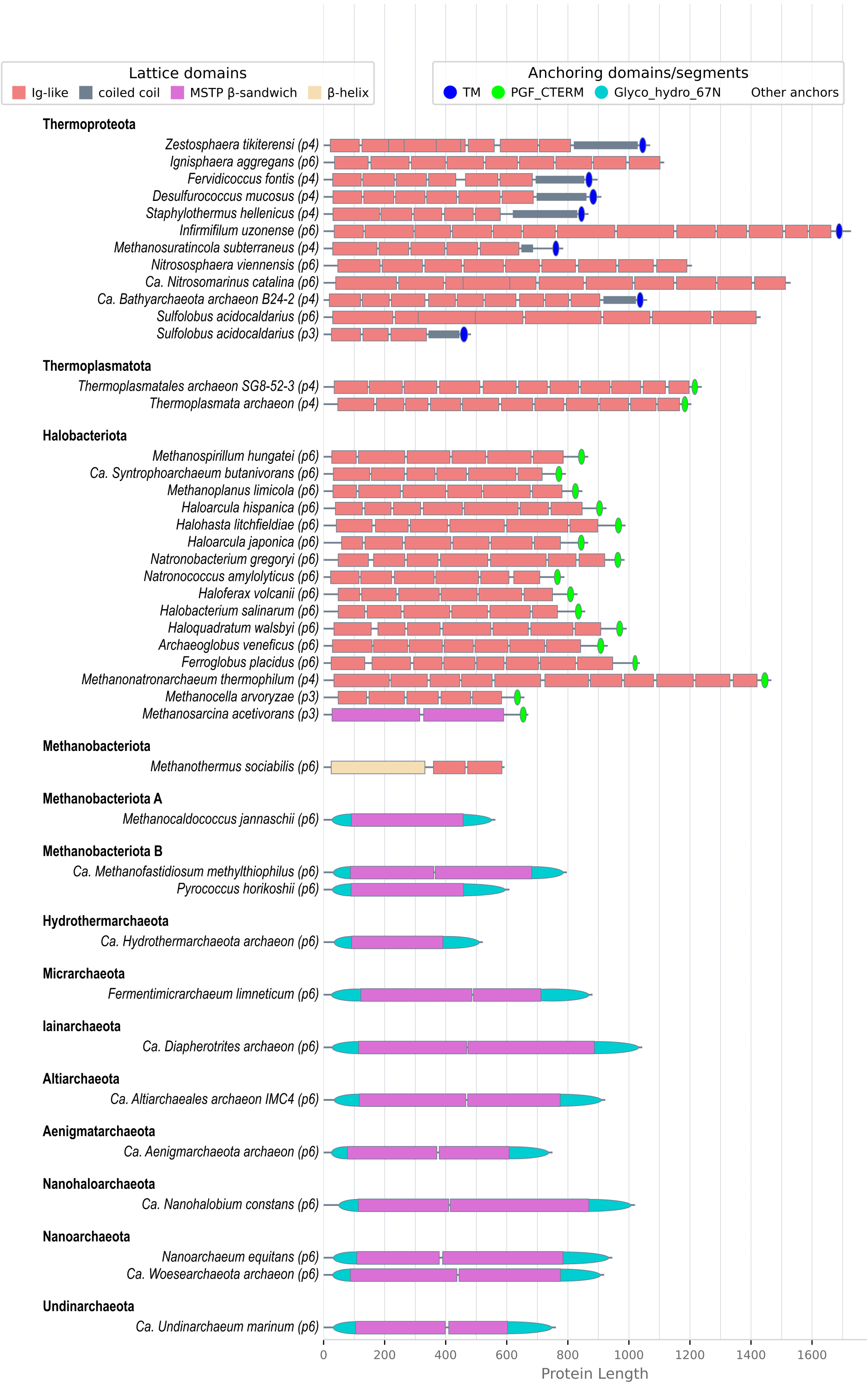


FIG. S7. Domain organization of archaeal SLPs. Three distinct folds characterize the lattice-forming domains in archaea: the Ig-like fold observed in the phyla Thermoproteota, Thermoplasmatota, and Halobacteriota; the $\beta$-helix fold found in some species of the Methanobacteriota phylum, including *Methanothermus sociabilis*; and the Methanosarcinales S-layer Tile Protein (MSTP)-like $\beta$-sandwich fold prevalent in the remaining phyla. Regarding anchoring mechanisms, archaeal SLPs employ transmembrane $\alpha$-helices (e.g., Thermoproteota), lipidation (Halobacteriota; PGF_CTERM; InterPro - IPR026371), a putative C-terminal Ig-like pseudomurein-binding domain in *Methanothermus sociabilis*, a putative C-terminal Ig-like membrane-binding domain in ammonia-oxidizing archaea (*Nitrososphaera viennensis*), a membrane-anchoring accessory protein (e.g., SlaB of *Sulfolobus acidocaldarius*), or a Glyco_hydro_67N-like domain exhibiting a Zincin-like fold in others. Additionally, numerous SLPs within the Thermoproteota phylum feature a coiled-coil stalk.

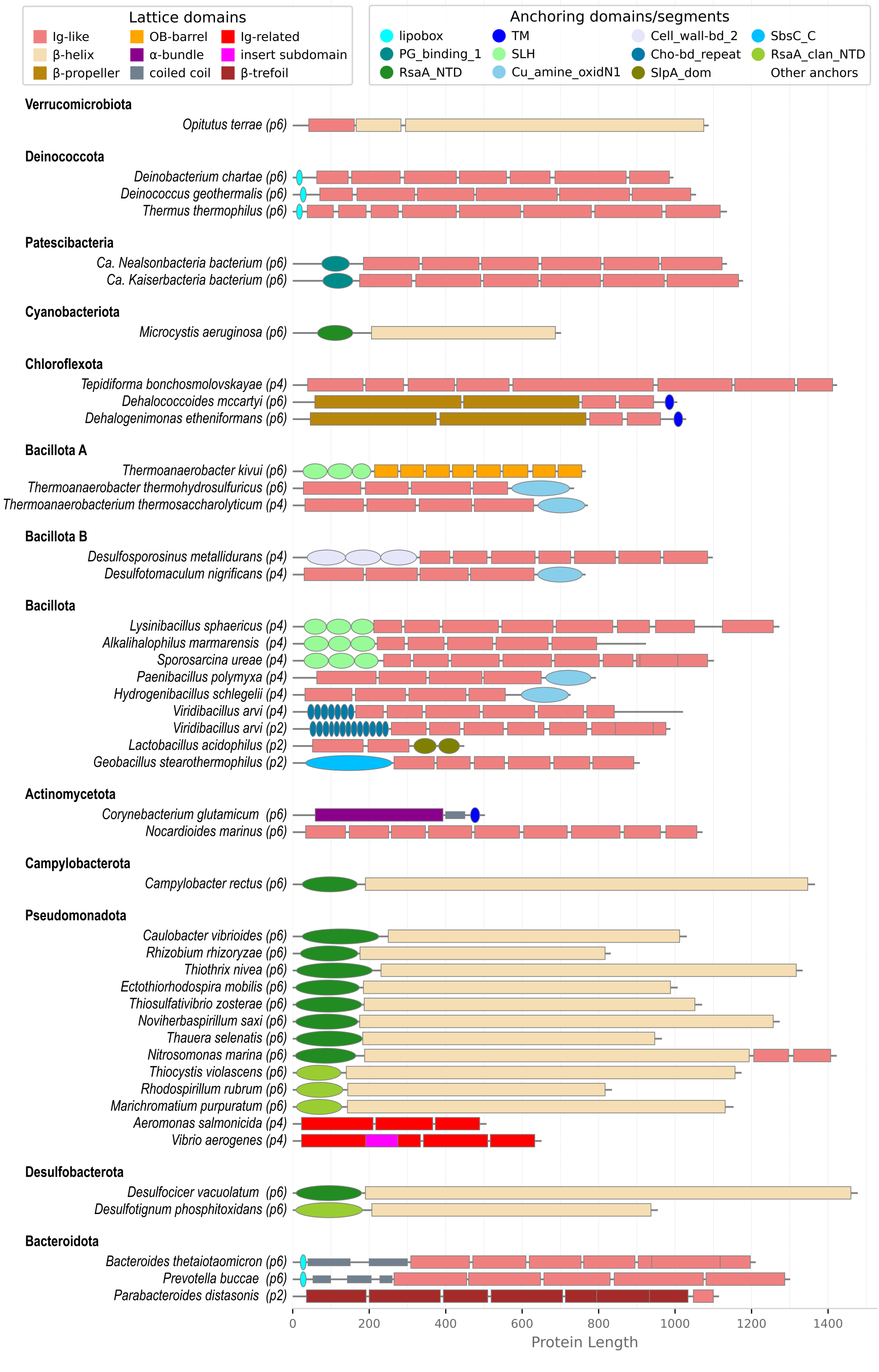


FIG. S8. Domain organization of bacterial SLPs. Compared to archaea, bacterial SLPs exhibit significantly more diversity in both their lattice and anchoring domains. Lattice domains display a variety of folds, including the Ig-like fold (*Thermus thermophilus*), $\beta$-helix fold (*Campylobacter rectus*), $\beta$-propeller fold (*Dehalococcoides mccartyi*), OB-barrel fold (*Thermanaerobacter kivui*), $\alpha$-bundle (*Corynebacterium glutamicum*), Ig-related fold (*Aeromonas salmonicida*), and $\beta$-trefoil fold (*Parabacteroides distasonis*). As for anchoring mechanisms, bacterial SLPs utilize a range of strategies, such as lipidation (*Thermus thermophilus*), transmembrane $\alpha$-helices (*Corynebacterium glutamicum*), PG_binding_1 (*Ca. Nealsonbacteria bacterium*; InterPro - IPR002477), RsaA_NTD (*Campylobacter rectus*; Pfam - PF19198), RsaA_clan_NTD (*Thiocystis violascens*), SLH domain (*Thermoanaerobacter kivui*; InterPro - IPR001119), Cu_amine_oxidN1 (*Desulfotomaculum nigrificans*; InterPro - IPR012854), Cell_wall-bd_2 (*Desulfosporosinus metallidurans*; InterPro - IPR007253), Cho-bd_repeat (*Viridibacillus arvi*; InterPro - IPR018337), SlpA_dom (*Lactobacillus acidophilus*; InterPro - IPR024968), SbsC_C (*Geobacillus stearothermophilus*; InterPro - IPR041378), and some putative Ig-like and Ig-related domains (*Opitutus terrae*, *Tepidiforma bonchosmolovskayae*, *Nocardioides marinus*, *Aeromonas salmonicida*, and *Parabacteroides distasonis*; InterPro - IPR045963).

supplemented with 10 nm gold conjugated with protein-A (CMC Utrecht).

#### C. Cryo-EM Data Collection and Processing

Cryo-EM data was collected from the prepared grids with a Titan Krios transmission electron microscope equipped with Falcon 4i direct electron detector (ThermoFisher) at 96 000 nominal magnification with a calibrated pixel size of 0.84 Å camera running in super resolution counting mode. EPU software was used to record movies in EER mode with a total electron dose of 50 $e^-/\text{Å}^2$ and defocus between $-1$ and $-2\,\mu$m. Data processing of cryo-EM data was performed using CryoSPARC v4.2.1 [192]. Movies were processed using patch-motion correction and patch CTF estimation. Particles were picked manually to produce initial 2D classes, which were then used as references for reference-based particle picking leading to a dataset containing 3763 particles. Particles were extracted with box size of 1200 pixel$^2$ and Fourier cropped to 150 pixel$^2$. Particles were 2D classified and bad looking classes were discarded to obtain the final set with 615 particles. Homogeneous refinement was performed with the final set of particles by applying $C_6$ symmetry, which produced 3D volume at 16.87 Å resolution. Predicted PS2 lattice (lattice constant 172 Å) was fitted into the cryo-EM volume using UCSF ChimeraX 1.6.1 [193]. Additionally, predicted PS2 hexamers were fitted separately into cryo-EM volume, which created a lattice with a lattice constant of 170 Å.

#### D. Cryo-ET data collection and processing

Cryo-ET data was collected as previously described [194]. Briefly, a Titan Krios transmission electron microscope equipped with K3 direct electron detector (Gatan) and Quantum energy filter (slit width 20 keV) was used for data collection. The SerialEM software [195] was used to acquire tilt series bidirectionally at 42 000 magnification (calibrated pixel size of 2.13 Å) in counting mode with a nominal defocus of $-5.5\,\mu$m, $\pm 60^\circ$ oscillation and $1^\circ$ tilt increment with a total dose of 116 $e^-/\text{Å}^2$. Tilt series were aligned by gold fiducial tracking, followed by tomogram generation in IMOD [196].

### S3. FURTHER RESULTS: STRUCTURE PREDICTION

#### A. Archaeal S-layers

Figures S10-S13 show our confident archaeal S-layer crystal predictions. We can divide our results into four broad types of S-layer architecture.

The first type is what we term the hexagonal spiral pyramids. All have a $p6$ assembly layer with the N-terminal domain next to the $C_6$ axis at the exterior-facing tip of the pyramid and the protein adopting a left-handed spiral geometry. The archetypal example is that of *Haloferax volcanii* and consists of six assembly domains. This type is common in the Halobacteria class of the Halobacteriota phylum, but also extends into the classes Methanomicrobia (e.g. *Methanospirillum hungatei*) and Syntropharchaeia (e.g. *Syntrophoarchaeum butanivorans*). The SLPs for these three taxonomic classes form three closely-linked clusters in Fig. S5, indicating that they are homologous. Generally, AlphaFold is relatively easily able both to locate the hexam-

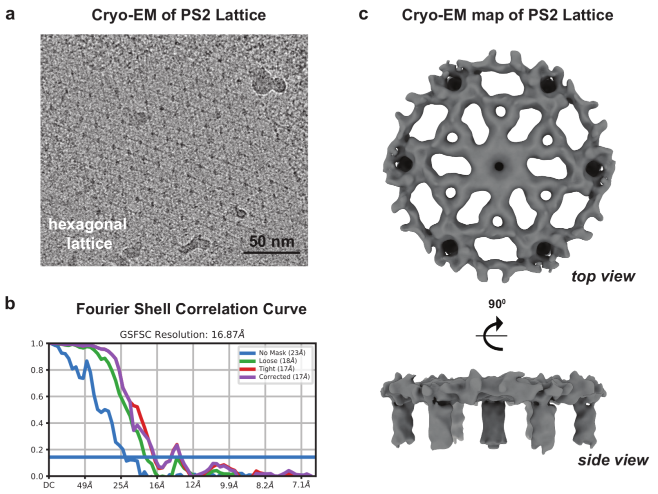


FIG. S9. Cryo-EM structure of *C. glutamicum* S-layer. (a) Cryo-EM image of purified *C. glutamicum* S-layer showing a lattice with hexagonal arrangement. (b) Fourier shell correlation (FSC) resolution estimation for the cryo-EM map of the S-layer. (c) Cryo-EM reconstruction of S-layer top and side views, as labelled.

TABLE S3. Cryo-EM data collection statistics

| | |
|---|---|
| Microscope | Titan Krios G2 |
| Magnification | 96 000 |
| Voltage(kV) | 300 |
| Electron Exposure ($e^-/\text{Å}^2$) | 50 |
| Detector | Falcon 4i (ThermoFisher) |
| Defocus range ($\mu$m) | $-1$ to $-2$ |
| Acquisition Mode | Super resolution |
| Pixel size (Å) | 0.84 |
| AFIS mode | No |
| Micrographs used (no) | 31 |
| Software reconstruction | CryoSPARC v4.2.1 |
| Software picking | CryoSPARC v4.2.1 |
| Initial particle images | 3763 |
| Final particles | 615 |
| Final box size (pix) | 150 |
| Pixel size final reconstruction (Å) | 3.296 |
| Symmetry imposed | $C_6$ |
| Map resolution (Å) | 16.87 |
| FSC threshold | 0.143 |

TABLE S4. Cryo-ET data collection statistics

| | |
|---|---|
| Microscope | Titan Krios G3 |
| Magnification | 42 000 |
| Voltage(kV) | 300 |
| Electron Exposure ($e^-/\text{Å}^2$) | 116 |
| Detector | K3 (Gatan) |
| Slit width (eV) | 20 |
| Defocus range ($\mu$m) | $-5.5$ |
| Acquisition Mode | Counting |
| Pixel size (Å) | 2.13 |
| Tilt-series increment | $\pm 1°$ |
| Tilt-series scheme | dose-symmetric |
| Tilt-series range | $\pm 60°$ |
| Software tilt-series alignment | IMOD (Kremer et al, 1996) |

eric and trimeric interactions relevant to the S-layer and to combine them into an accurate crystal model. Sometimes, predictions for hexamers of the C-terminal section generate a $C_2$ dimer of trimers that can also provide a direct model of the $C_2$ site. Even where available, this is generally not used to generate the crystal model as the combination of the hexamer and trimeric complexes naturally also give a good model of the $C_2$ site, albeit with occasional minor overlaps of the sixth domain whose geometry is not always fully constrained by the trimer.

The examples have been chosen mainly either because they have been studied experimentally or provide insight into the taxonomic extent or the structural and functional variations. For example, *Halorutilus salinus* is a representative of the recently discovered order Halorutilales in the class Halobacteria [197]. The strain of *Haloquadratum walsbyi* considered is unusual in that it has a double S-layer [36]. Whether the SLP considered is responsible for the inner or outer S-layer, or both is not clear. One interesting feature of this SLP is that it has an exterior domain at its N-terminus that is a cohesin domain. Cohesins generally bind to dockerin domains, and in *H. walsbyi*, the halomucin hmu3, as well as the third most highly expressed protein (UniProt ID: Q18KV7) [35], possess dockerin domains, presumably allowing them to bind to the exterior of this S-layer.

Interestingly, *Methanolliviera hydrocarbonicum*, a representative of the class Methanoliparia, only possesses a version of this S-layer gene with equivalents of the fourth to sixth assembly domains, suggesting that it might exhibit a $p6$ S-layer but with the top of the pyramid missing. However, as the MSA for this example was very shallow, we were not able to fully confirm this, but did find a similar trimeric complex (Fig. S17).

There is a second variant of the hexagonal pyramid that is common in the Halobacteria (and seems specific to this class). This set of SLPs was first identified for *Haloarcula hispanica* [38] and forms a well-separated cluster in Fig. S5. Some halobacteria possess genes for both variants, e.g. *Haloarcula japonica* [38], where the SLP consists of five assembly domains. In many examples, there is an additional N-terminal domain that although not involved in inter-protein contacts, is closely associated with the first assembly domain. The first two assembly domains form a confident hexamer that is the basis for the apex of the pyramid. However, the AlphaFold predictions for the three C-terminal assembly domains often exhibit confident dimer and trimers that involve similar interfaces, i.e. they are mutually exclusive. Our best estimate is that the trimer is more likely to be representative of the S-layer and so all our crystal models have been generated using the trimer predictions.

Another difficulty with this set of hexagonal spiral pyramids is that some examples have a relatively flexible linkage between the second and third assembly domains (as indicated, for example, by the PAE plots for the monomer). As this is relatively unconstrained by either the hexameric or trimeric complexes, in these cases there is an additional uncertainty with regards to this inter-domain angle in the crystal predictions and we sometimes see overlaps that could probably be avoided by adjustment of this angle. This feature also leads to an S-layer that is more open than *Haloferax volcanii* and has larger pores (e.g. this is evident in Fig. S10 for *Halogeometricum borinquense*). However, in other examples (e.g. *Natronobacterium gregoryi*), this flexibility is less evident and these tend to from a denser S-layer that looks more similar to *Haloferax volcanii*.

A third variant in the Halobacteriota phylum is specific to the class Archaeoglobi and is illustrated for *Archaeoglobus veneficus* in Fig. S10. The SLP in this case has seven assembly domains. As an example of this variant, we also considered a putative SLP from *Ferroglobus placidus*. Although this shows a very similar lattice, in this case it is likely to represent a silent/cryptic gene that is not the usual SLP, because freeze-fracture experiments have shown that the S-layer of *F. placidus* has $p4$ symmetry [49]. The identity of the SLP responsible for this $p4$ S-layer, however, has not been identified.

Hexagonal spiral pyramids are also one of the major architectural types for $p6$ Thermoproteota S-layers. The evolutionary relatedness of the SLPs to those in Halobacteriota is not fully clear. Despite being homologous due to their Ig-like domains, the low sequence identity between them ($< 15\%$) makes it difficult to determine whether they arose from a common ancestral SLP or if they arose independently in the two lineages, possibly through the amplification of Ig-like domains. One variant is exemplified by *Nitrososphaera viennensis*. The SLP has eight assembly domains with a ninth domain at the C-terminus lying below the S-layer that is likely involved in S-layer anchoring. As well as the class Nitrososphaeria, examples of this variant have also been found in the classes Thermoprotei A, Methanomethylicia, and Bathyarchaeia (order Hecatellales), albeit we were not always able to obtain a complete crystal prediction because of a combination of their relatively large size and sometimes shallow MSAs.

A subvariant of this type, but with much larger domains near to the C-terminus, are exemplified by *Ni-*

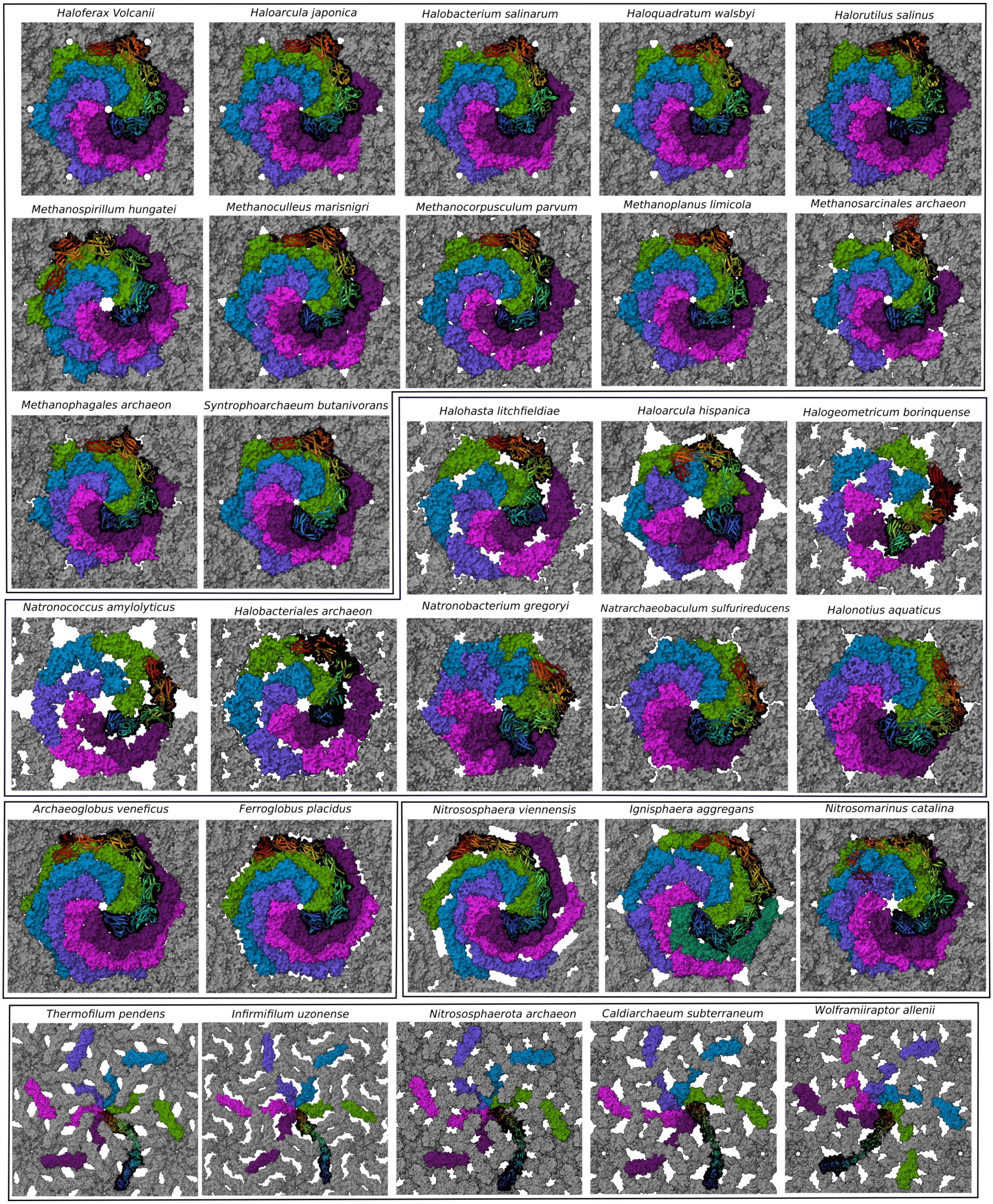


FIG. S10. 2D crystal structures of the archaeal $p6$ S-layer predictions for proteins composed of Ig-like domains. The boxes group the S-layers into those with significant structural and sequence similarity.

trosopumilus maritimus* and *Nitrosotalea devanaterra*. Full crystal predictions are particularly difficult for these examples, as memory limitations only allow us to consider hexamers of the first two domains and only the seventh and eighth domain interact at the trimeric interface. How the intermediate domains adapt their structure compared to the monomer to gain interactions and avoid overlaps in the hexagonal pyramid is required for a good crystal prediction.

Hexagonal spiral pyramids are also common in the order Sulfolobales of Thermoprotei A, as evidenced by many electron microscopy studies [75, 76, 80–83]. We considered most examples for which there have been experimental structural studies; however, the shallowness of the MSAs generally make crystal predictions unfeasible and most examples have poor AlphaFold monomer predictions (i.e. with low pLDDT and a significant amount of disorder).

We discovered a second architectural type of $p6$ S-layer that appears specific to Thermoproteota. This is exemplified by *Thermofilum pendens*. In this example, the SLP is the largest protein in the genome and consists of 13 domains. The six domains nearest to the C-terminus form a hexagonal cylinder/cone that extends down away from the S-layer and at the end of which are transmembrane $\alpha$-helices that facilitate cell anchoring. The N-terminal sub-protein forms an interwoven triangular motif that lies in the plane of the S-layer, and in this case mainly involve interactions between the third to fifth domain. On building the crystal we noticed that the N-terminus is positioned close to the domains at the "mouth" of the hexameric cone. AlphaFold predictions on a heterodimer of the first two domains with the sixth and seventh domains suggested an interaction between the two N-terminal domains with the seventh domain. The interwoven character of the S-layer and the extensive inter-protein interactions that it involves may help to explain the extreme stability of the *T. pendens* S-layer [198].

As well as in the class Thermoproteia A, more distant homologs were found in the class Nitrososphaeria A and also seemed to be the predominant type of S-layer in this class. Further examples were also found in the classes Methanomethylicia (*Culexarchaeum yellowstonense* and *Nezhaarchaeota archaeon*), Bathyarchaeia, and EX4484-205 (*Brockarchaeota archaeon*). It was not always possible to get a good crystal prediction, even when hexameric and trimeric complexes were identified, because it requires the predicted structure of the monomer in the relatively large intermediate section between the hexamer and trimer to be representative of the S-layer. Furthermore, the size of the angular distortion required to make the two symmetry axes co-parallel generally increased with protein size.

There is a third very extensive architectural type of $p6$ archaeal S-layer (Fig. S11). The SLPs in this class have a very distinctive topology and involve an assembly domain consisting of one or two $\beta$-sandwich-like domains and a cell-anchoring domain that is at the end of a $\beta$-sheet "stalk" and is formed from "partial-domains" at the N and C termini. This distinctive topology (and its resulting distinctive PAE diagram) allows relative easy identification of this type of SLP, even in the absence of confident predictions of complexes by AlphaFold (e.g. *Korarchaeum cryptofilum*).

The proteins of this third structural type typically form strong dimers mediated by interactions between the anchoring domain, the $\beta$-sheet "stalk", and the assembly domain. They also all form a hexameric interaction involving one end of the assembly domain. The six monomers around the $C_6$ axis form a star-like geometry, which in the S-layer often interlock to form a dense layer with only relatively small pores. The exceptions to the latter are those associated with the phylum Hydrothermarchaota and the order Thermococcales in the phylum Methanobacteriota B (e.g. *Pyrococcus horikoshi*). In the latter, the dimeric interactions between the assembly domain are mediated by $\beta$-sheet sidearms. The other main variations are the size of the assembly domain (typically biggest for those associated with the DPANN phyla) and the angle by which the assembly domain is tilted upwards with respect to the plane of the S-layer on moving away from the $C_6$ axis (e.g. for *Methanocaldococcus jannaschii* this angle is approximately 45°, whereas for some of the DPANN examples, the assembly domain can be close to planar).

This type of S-layer appears to be the predominant S-layer in the phyla Methanobacterioata A, Methanobacterioata B, Hydrothermarchaeota, the DPANN phyla, and the class Korarchaeia of Thermoproteota. These examples form a set of inter-connected clusters in Fig. S5 with the clusters aligning relatively closely with the respective phyla, thus confirming their evolutionary interrelatedness.

Generally, the success rate of complete crystal predictions is high for this type of S-layer, the main problem being the shallow MSAs for some of the DPANN examples (e.g. we were only able to obtain a prediction for *Nanoarchaeum equitans* when we supplemented the MSA produced by MMseqs2 with one produced by the JackHMMER algorithm) and for all the Korarchaeia examples that we considered. The shallowness of the MSA is typically evident from the AlphaFold-predicted monomer (e.g. low pLDDT and structural disorder).

The *Altiarchaeum hamiconexum* SLP is interesting in a number of ways. Firstly, in microscopy studies of organisms in this phylum, S-layers have not been previously identified. Secondly, this particular protein had been identified [199] as being the main component of the "hamus", a one-dimensional fibrillar assembly that protrudes from the cell and mediates intercellular aggregation [200]. Our results clearly show that this is a misidentification (there is only a very clear propensity to form 2-dimensional assemblies). Presumably, its experimental identification as a fibril was because of its presence at the cell surface in significant quantity; however, our

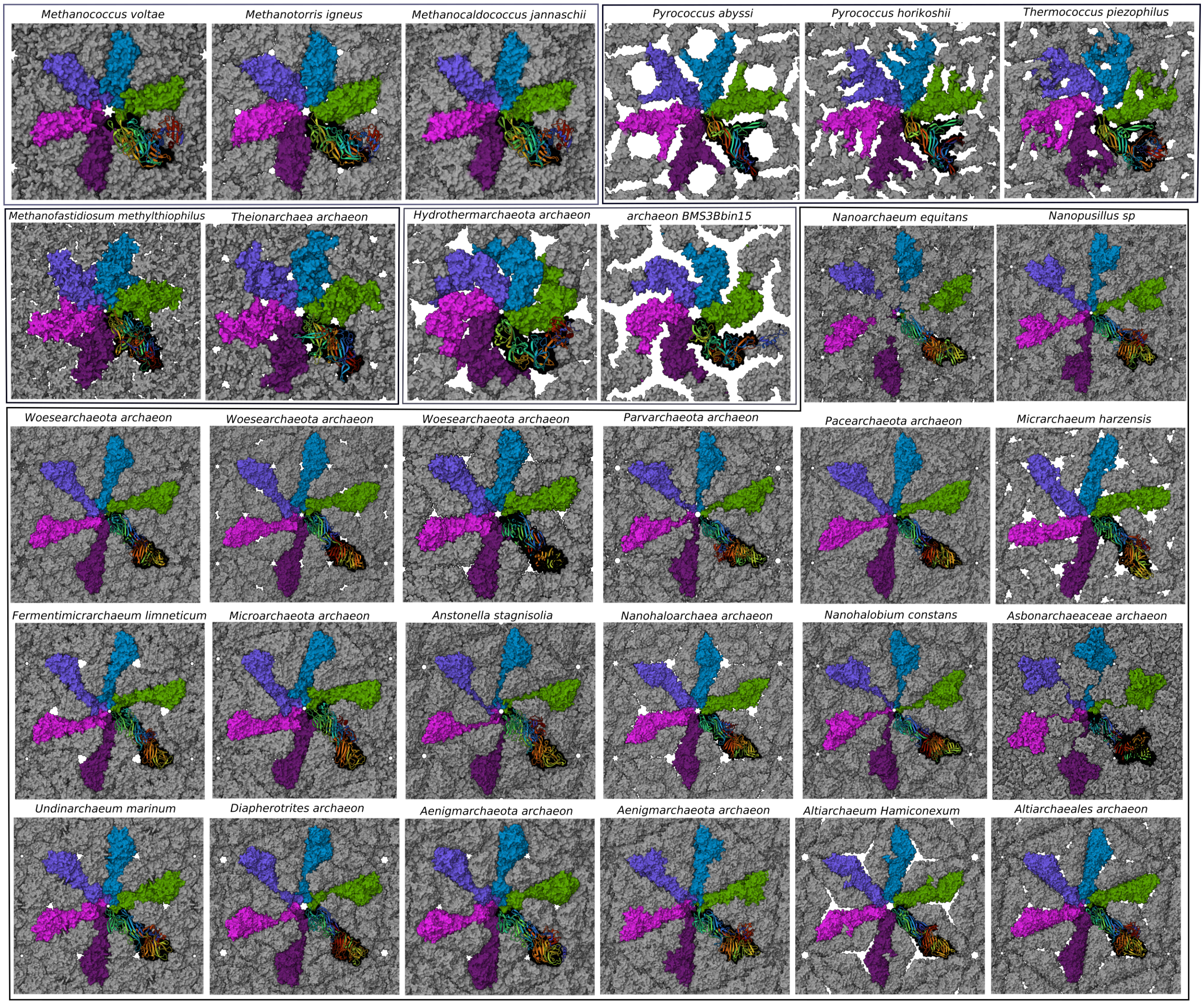


FIG. S11. 2D crystal structures for the hexagonal $\beta$-sandwich archaeal S-layer crystal predictions. The boxes group the S-layers into those with significant structural and sequence similarity.

results suggest that this protein could form S-layers to some degree in these organisms.

We have located two types of S-layers with a $p3$ assembly layer (n.b. the $p6$ character of the assembly layer in some of the S-layers in the order Sulfolobales appears to be reduced to $p3$ by the binding of trimers of the SlaB cell-anchoring protein [21, 28]). Why S-layers with $p3$ symmetry are relatively uncommon is not clear (note we have identified no $p3$ S-layers for bacteria).

The first type of a $p3$ S-layer is exemplified by *Methanosarcina acetivorans*. The SLP, also referred to as the Methanosarcinales S-layer Tile Protein (MSTP), is made up of $\beta$-sandwich domains, homologous to those in the SLPs of the DPANN phyla, but has two distinct features compared to the other $\beta$-sandwich S-layers. First, it does not possess an anchoring domain but instead has a PGF-CTERM signal peptide that gets replaced by a lipid by an archaeosortase (this anchoring mechanism seems to be universal in the Halobacteriota phylum (Fig. S7)). Secondly, the monomer has two $\beta$-sandwich domains in a pseudo-dimeric arrangement. These $\beta$-sandwich domains pack quite similarly to many of the other $\beta$-sandwich S-layers, but because the two domains are not equivalent, the symmetry of the S-layer is reduced from $p6$ to $p3$. Interestingly, this category of SLPs is much closer to the other Halobacteriota SLPs in the cluster map (Fig. S5) than to the $p6$ $\beta$-sandwich S-layers. The $\beta$-sandwich domains have highly divergent sequences, making their sequence similarity undetectable with BLAST. However, it can be detected using the more sensitive sequence comparison method, HHpred. In the cluster map, which is based on BLAST, the SLPs of this category cluster to-

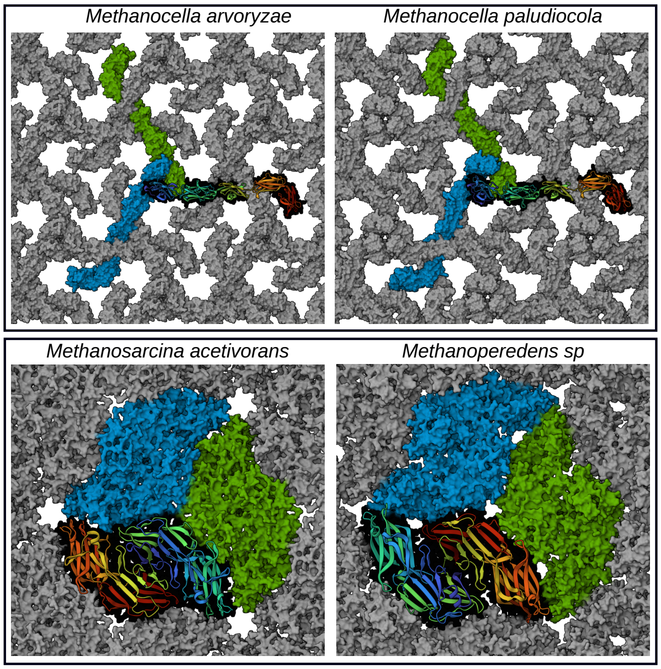


FIG. S12. 2D crystal structures of the archaeal $p3$ S-layer predictions.

gether with the SLPs of Halobacteriota due to the sequence similarity between their PGF-CTERM motifs.

A model of the *Methanosarcina acetivorans* was previously obtained from 3-dimensional crystals of the C-terminal $\beta$-sandwich domain; the crystal consisted of stacking of $p6$ layers [52]. By considering multimers of this individual domain we can also obtain a prediction for this $p6$ 2-dimensional crystal (Fig. S14). It is essentially the same as the structure observed experimentally [52] and both are equivalent to the S-layer of the full protein, except for the loss of symmetry due to the inequivalence of the N- and C-terminal domains. Note, if in the actual S-layer the orientation of the monomers was random, it would on average have $p6$ symmetry. However, there is no way of obtaining an orientationally-ordered $p6$ S-layer.

One methodological curiosity of this type of S-layer proteins is that AlphaFold finds it much easier to locate the $C_3$ hexamers of the individual $\beta$-sandwich domains than the equivalent $C_3$ trimers of the complete protein. Why this is the case, is not at all clear.

The other class of $p3$ archaeal S-layers appears specific to the Methanocella class of Halobacteriota. The SLPs in this class typically consists of five Ig-like domains with trimeric interactions involving the N-terminal domain, the middle domains, and the C-terminal domain occurring at the three unique $C_3$ sites in the $p3$ crystal.

All the $p4$ archaeal S-layers (Fig. S13) that we were able to characterize have some basic architectural similarities. They involve a relatively planar array of Ig-like domains that form a relatively open mesh in the plane of the S-layer. The C-terminal domains always lie at a $C_4$ axis and are also associated with anchoring. However, it is not clear whether those associated with the phyla Thermoplasmata and Halobacteriota are evolutionarily related to those in the phylum Thermoproteota (Fig. S5).

Interestingly, we were able to identify $p4$ S-layers in the phylum Halobacteriota associated with the class Methanonatronarchaeia. The only previously observed $p4$ S-layer in this phylum is for *Ferroglobus placidus*; however the identity of the SLP is unknown in this organism, as mentioned above. Note, in both examples from Methanonatronarchaeia the MSA associated with the N-terminal half of the protein was too shallow for predictions to be made about this portion of the protein, so whether these parts are also involved in additional interactions that promote assembly is unclear.

Shallow MSAs also affected our consideration of putative Thermoplasmata SLPs. For example, for *Picrophilus oshimae* we found a tetrameric complex (Fig. S17) that matched well the experimental electron-microscopy reconstruction, but we were unable to locate any further confident interactions that would enable the building of a crystal model. For *Aciduliprofundum boonei*, another example from Thermoplasmata where a $p4$ S-layer has been experimentally observed [201], although we were reasonably confident of the SLP's identity (UniProt ID: D3TBL7) AlphaFold was unable to even predict a reliable structure of the monomer due to the shallow MSA.

By contrast, for the class Bathyarchaeia of Thermoproteota, we were able to reveal a rich array of $p4$ S-layer structures due to the wealth of sequence data obtained from metagenomics studies. Only very recently has the presence of an S-layer been experimentally observed for this class [202]. We were also able to identify $p4$ SLPs in the classes Methanomethylicia (the presence of an S-layer for a *Methanosuratincola* species has recently been confirmed [203]) and Nitrososphaeria for the first time.

A number of $p4$ S-layers associated with the order *Sulfobales* have been experimentally characterized by electron microscopy. Despite the generally shallow MSAs we were able to obtain a number of good crystal predictions. Our predicted structure for *Desulfurococcus mobilis* is in excellent agreement with the experimental reconstruction [70]. The predicted S-layer for *Zestosphaera tikiterensis* is very similar, except that the protein has an additional sidearm that associates about the second $C_4$ axis in the crystal. Interestingly, the structure looks very similar to that reported for an Aeropyrum strain [74], however, it shares relatively little sequence similarity with the SLP of *Aeropyrum pernix* (which has an extremely shallow MSA and hence no predictions).

*Staphylothermus marinus* provides another interesting example. Experimentally the S-layer has been shown to assemble into very stable "tetrabrachions", which can further assemble into mesh-like assemblies (note although the S-layer is often denoted as being $p4$, the available images only show relatively short-range order) [72]. Another interesting feature is that the SLP has been shown to be cleaved into two sub-proteins [71]. Despite a shallow MSA we were able to obtain a good prediction for the

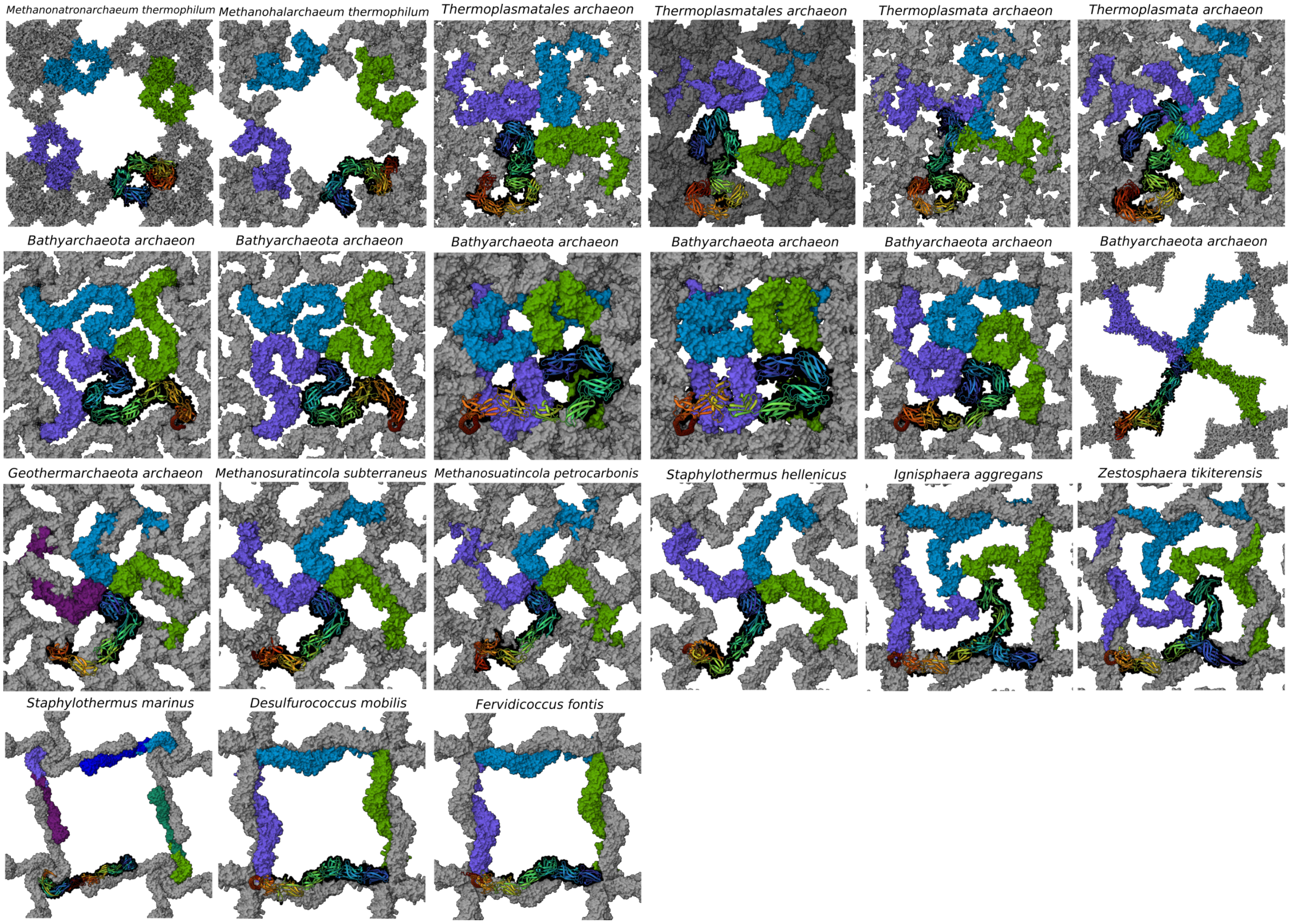


FIG. S13. 2D crystal structures of the $p4$ archaeal S-layer crystal predictions.

tetrabrachion (Fig. S15). The reason for the cleavage is not evident from the assembled structure as the cleaved ends of the two proteins are adjacent in the structure. We found that the arms of the tetrabrachion were able to dimerize but at an angle that would give rise to significant curvature of any assembly not involving significant deformation of the individual tetrabrachions. Perhaps consistent with this, the isolated tetrabrachions were observed to assemble into micellar-like structures *in vitro* [72]. Although we have generated a $p4$ crystal model, it should be borne in mind that this involved significant deformation of the tetrabrachion arms.

A $p4$ S-layer has also been experimentally observed for *Pyrolobus fumarii* [204]. Although we have likely identified the SLP (UniProt ID: G0ECC9), due to the shallow MSA, AlphaFold was unable to provide significant further insight, beyond confirming the presence of a C-terminal $\alpha$-helix that is typical of the Sulfobales $p4$ SLPs.

A final interesting interesting archaeal example is the S-layer of *Methanothermus fervidus*/*Methanothermus sociabilis* (note their SLPs are almost identical, having over 99% sequence identity). The SLPs consists of an N-terminal $\beta$-helix and two Ig-like domains and so do not fit into any of the main classes of archaeal S-layers that we have already discussed. These organisms have been reported to have a $p6$ S-layer with lattice constant of 20 nm [54]. A structure matching those criteria can be built from our multimer predictions for these proteins, however, the confidence metrics do not clearly identify the hexameric cluster used to build the crystal as being particularly stable compared to other sizes. So, the overall prediction should be seen as plausible but tentative. In this crystal the $\beta$-helices forms a hexagonal cylinder that are connected to each other by dimeric interactions between the C-terminal Ig-like domains. These domains are also likely to mediate binding to the pseudomurein that covers the cell; they are found in many *Methanothermus* cell-surface proteins and their sequence, particularly at the C-terminus, is highly conserved.

### B. Bacterial S-layers

Figures S18-S20 show our confident bacterial S-layer crystal predictions. The three figures show those with

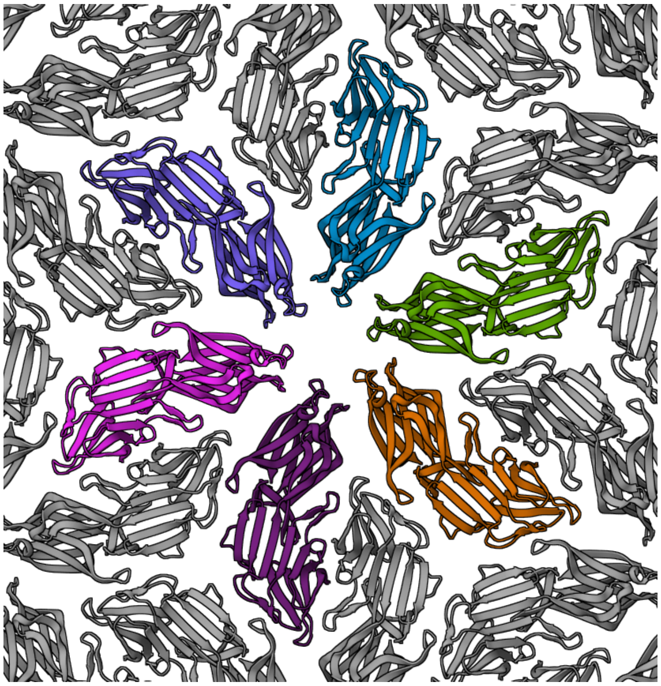

FIG. S14. $p6$ crystal of the C-terminal $\beta$-sandwich domain of *Methanosarcina acetivorans*.

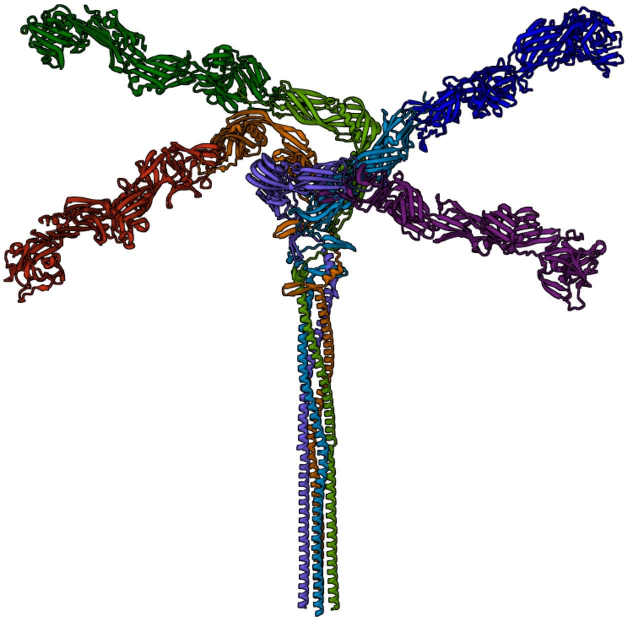

FIG. S15. The top of the *Staphylothermus marinus* tetrabrachion, consisting of 4 copies of the S-layer protein, cleaved into the "light" and "heavy" chain. The bottom of the coiled coil is not represented.

$p6$, $p4$, and $p2$ symmetry, respectively.

For bacteria, as noted in the main text, there are far fewer clear relationships between the S-layers in different bacterial phyla and the different types of S-layers generally form well-separated "islands" in the cluster map of Fig. S6. The majority of the SLPs we have considered come either from Pseudomonadota and nearby phyla (Desulfobacterota and Campylobacterota) or from the Bacillota phyla, as bacterial S-layers from these phyla have been the most studied experimentally.

Our successful predictions for the S-layers of Pseudomonadota and nearby phyla can be divided into two broad types, namely $p6$ S-layers where the assembly domain is a $\beta$-helix and $p4$ S-layers where the protein consists of three Ig-related domains. Concerning the first group, many are like the experimentally-determined *Caulobacter vibrioides* (or *Caulobacter crescentus*), where the $\beta$-helix has an L-shape. Generally, the assembly layers are relatively planar and the main variations are associated with whether the predominant secondary interaction is dimeric or trimeric and the positions on the $\beta$-helices at which these interactions occur (by contrast the hexameric interaction is always at the N-terminal end of the $\beta$-helix) and the lengths of SLP. Some, such as *Cupriavidus*, also have side-domains off the main $\beta$-helix.

Of these examples, *Thauera selenatis*, a selenate respiring bacterium, is interesting because a protein called SefA (Se factor A) has previously been implicated in the formation and export of intracellular selenium nanoparticles, it being suggested that it binds to Se, regulating the

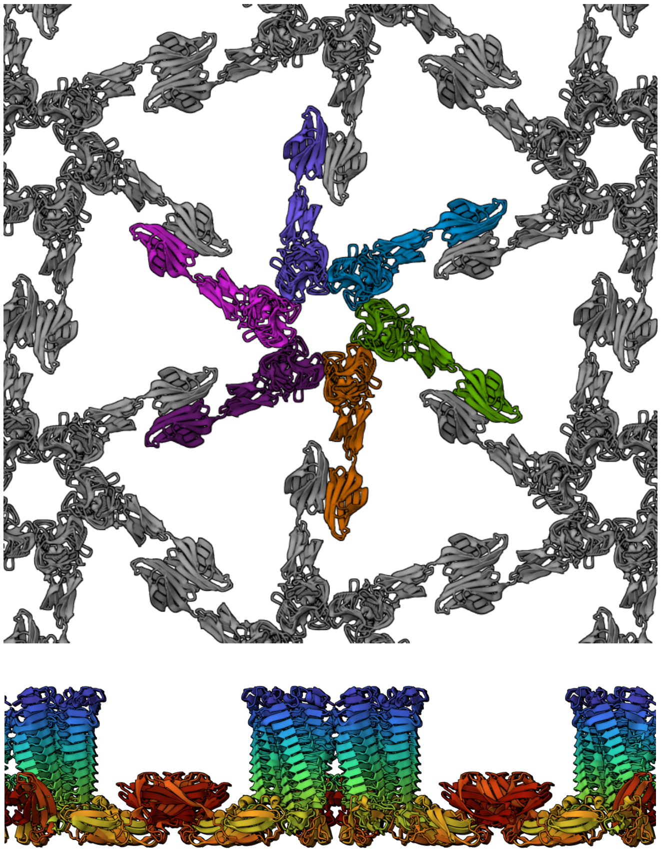

FIG. S16. Tentative prediction for the $p6$ S-layer of *Methanothermus sociabilis*.

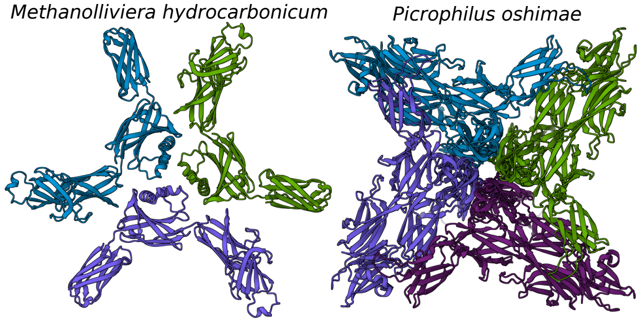


FIG. S17. Oligomer predictions for archaeal S-layer proteins.

nanoparticle formation and facilitating its export [205]. Our results show that SefA in its structural and assembly behaviour is a relatively typical Pseudomonadota SLP, which could be potentially interesting for the implications on SefA's cellular function.

The remaining $p6$ S-layers tend to be based on more bowl-shaped hexameric complexes. The example that has been experimentally most well-characterised is that for *Campylobacter rectus* and our predicted structure is in good agreement with the electron microscopy reconstruction [115]. There is significant variation in this broad group. Some are stabilized by dimeric interactions between a side-domain that lies below the main assembly layer (e.g. *Methylotuvimicrobium kenyense*). In the case of *Giesbergia anulus* and related examples, this was the only secondary interaction that we were able to identify that mediated interactions between the hexamers.

Interestingly, the above types of $\beta$-helix $p6$ S-layers extend beyond Pseudomonadota and into the nearby phyla Campylobacterota and Desulfobacterota. The latter is particularly noteworthy as S-layers had not been identified in this phylum previously. Also, there is evidence of horizontal transfer of these S-layer genes both into Verrucomicrobiota (e.g. *Opitutus terrae*) and Cyanobacteriota (e.g. *Microcystis aeruginosa*). The $\beta$-helix $p6$ SLPs form one large cluster in Fig. S6.

Although the Pseudomonadota $p4$ S-layers all have the same basic architecture, there is significant sequence diversity and in fact form two separated groups in Fig. S6. As noted in Section S1 D, although AlphaFold is generally able to locate the two tetrameric interactions that stabilize these S-layers, obtaining a reliable crystal prediction is challenging because the angle between the second and third domains is unconstrained by either tetrameric complex. We were only able to get sensible crystal predictions for three of the examples that we considered.

The *Delftia acidovorans* S-layer has been shown to consist of a tetragonal packing of rectangular units [110]. For this SLP, AlphaFold predicts an extremely confident dimer that has a large interface (Fig. S21). This dimer clearly corresponds to the rectangular unit observed experimentally. However, it was not possible to identify further interactions present in the crystal. This SLP is unusual in that it is a single-domain protein, thus making the approach of sub-dividing the protein to isolate other interactions harder. It did not prove possible to deduce the pattern of higher-order assembly from multiple dimers.

As already noted in Ref. [111], there are many homologs of the *Delftia acidovorans* SLP in *Pseudomonadota*, particularly amongst the *Betaproteobacteria*. A few examples are shown in Fig. S21. It is noticeable that the biggest shape difference is associated with the *Gammaproteobacteria* example, namely *Thiothrix fructosivorans*. Interestingly, *Simplicispira psychrophila* has been observed to possess a double S-layer, the inner one of which has features typical of this class of SLPs [111]. We were also able to identify a $\beta$-helix SLP in this organism (Fig. S18) and it is plausible that this is responsible for the outer S-layer. Similarly, the homolog in *Lampropedia hyalina*, which has a particularly complex S-layer, may be responsible for the inner part of that layer. This inner layer has been observed to consist of units located at the $C_2$ sites in the 2D crystal; these units have a shape that is consistent with the dimer in Fig. S21, however, they assemble into a hexagonal rather than a square lattice [206].

A number of bacteria in Pseudomonadota have been shown to have S-layers with particular large-scale features [101, 104]. One for which the SLP has been identified is *Methylotuvimicrobium alcaliphilum* [102]. The S-layer consists of large "cups" organised in an approximately hexagonal lattice with the cups being approximately 36 nm in diameter and 33 nm in height [104]. The SLP is particularly large and consists of multiple $\beta$-helix domains. From the AlphaFold calculations that we were able to perform, the tendency to form into cup-like structures is clear (Fig. S22) but with the number of monomers in each cup significantly larger than six—eighteen is probably the best estimate consistent with six-fold symmetry.

*Serratia marcescens* provides a final interesting example from Pseudomonodoata, as the putative SLP [105], although incorporating an L-shaped $\beta$-helix, is distinctly different from the other $\beta$-helix Pseudomonadota SLPs. For example, it has three $\alpha$-helical binding domains at its N-terminus (these sterically prevent the N-terminal end of the $\beta$-helix associating in a hexameric interaction similar to the other $\beta$-helix SLPs) and a distinctive feature at the "elbow" of the $\beta$-helix. We were unable to obtain any clear predictions suggestive of an ability to assemble in two dimensions, albeit our search was often hindered by sub-proteins interacting through the cut end or in a way that would involve steric overlap if the excised part of the protein was present. Given the lack of experimental observations of paracrystalline assembly it might be that this protein is just a highly-expressed surface protein rather than a 2D-crystal-forming SLP.

We were able to identify two types of S-layers in Bacteroidota, namely the $p6$ S-layers of *Bacteroides thetaiotaomicron* and *Prevotella buccae* and the $p2$ S-layer of *Parabacteroides distasonis*. After we observed no

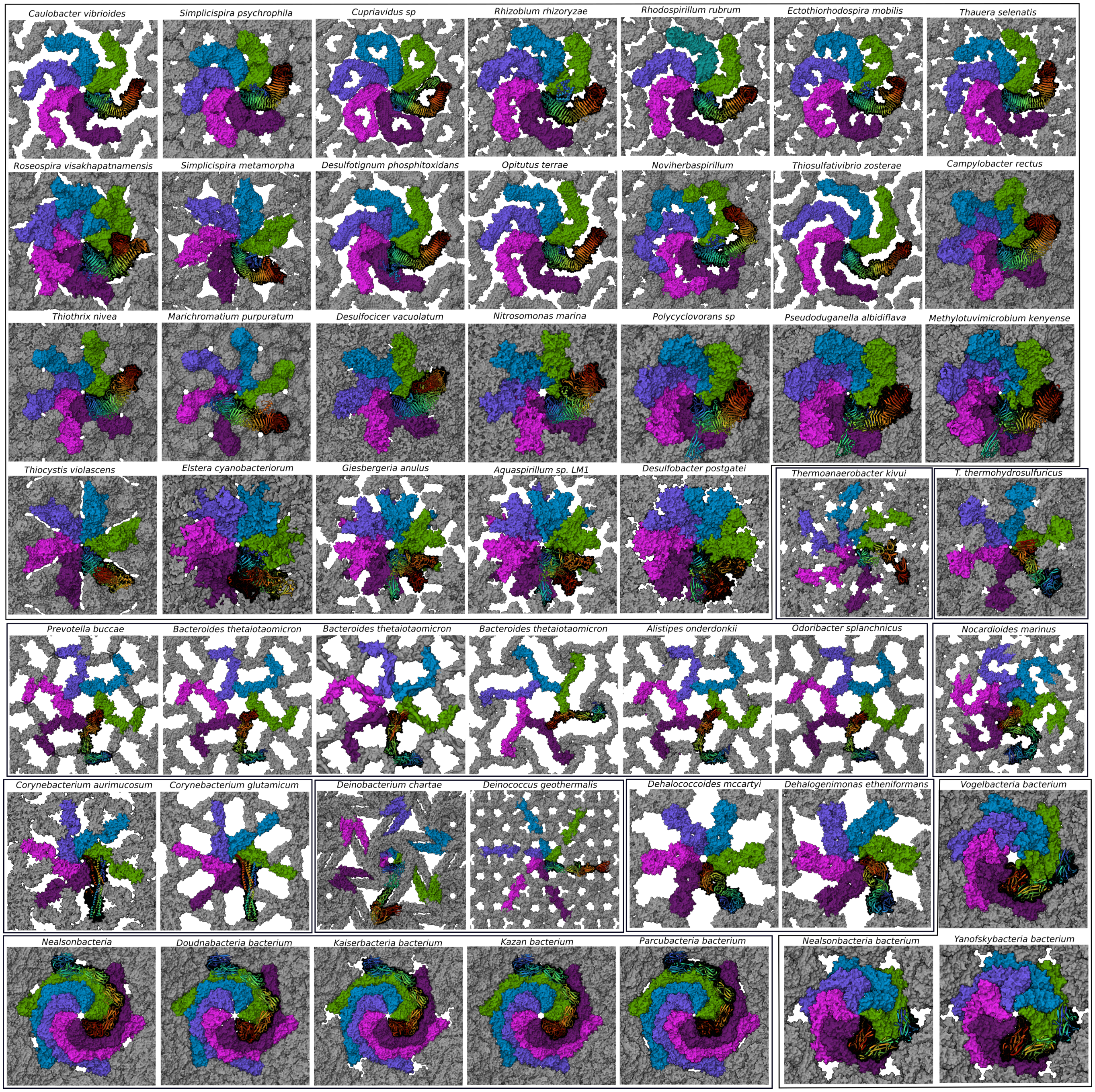


FIG. S18. 2D crystal structure predictions of the $p6$ bacterial S-layers.

propensity towards 2D assembly for the experimentally-suggested SLP for *Bacteroides thetaiotaomicron* [23], we scanned its genome for other putative SLPs and found a set with plausible AlphaFold-predicted monomer structures. These have an N-terminal $\alpha$ helix that forms a trimeric coiled coil that connects the assembly layer to the cell surface and then a series of Ig-like domains that form the assembly layer. Their discovery then allowed the identification of the equivalent protein for *Prevotella buccae* and its predicted structure agrees very well with the electron microscopy reconstruction of the S-layer for this organism. For *Bacteroides thetaiotaomicron*, the set of S-layer genes could be divided into two sub-types depending on whether the assembly layer involved an interaction at the $C_2$ site. This is also the origin of the two sub-clusters in Fig. S6.

The Bacillota phyla exhibit a wide variety of S-layers that span $p6$, $p4$, $p2$, and $p1$ symmetry. These have been comparatively well studied with the SLPs having been identified for a significant number of examples. Our ap-

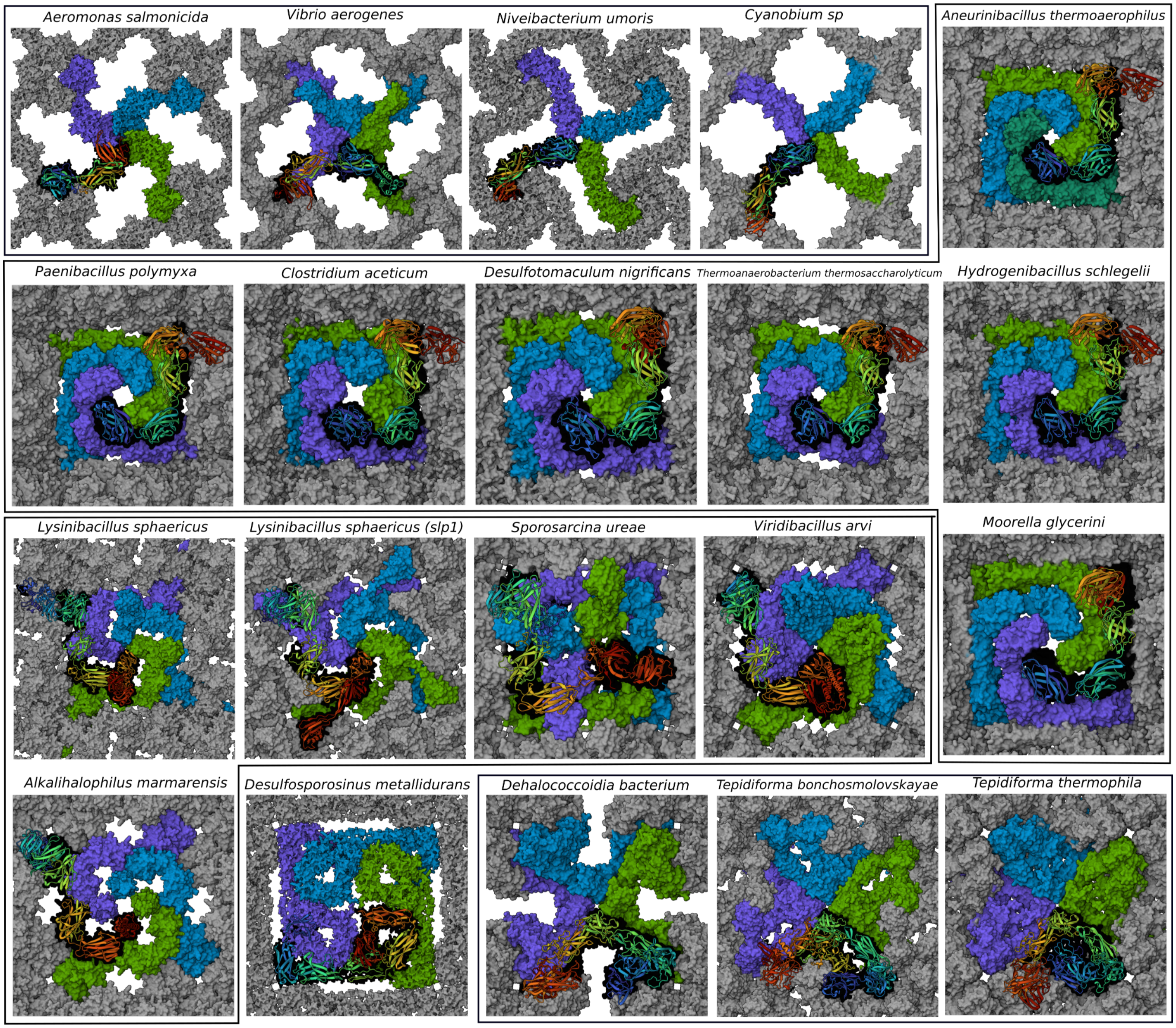


FIG. S19. 2D crystal structure predictions of the $p4$ bacterial S-layers.

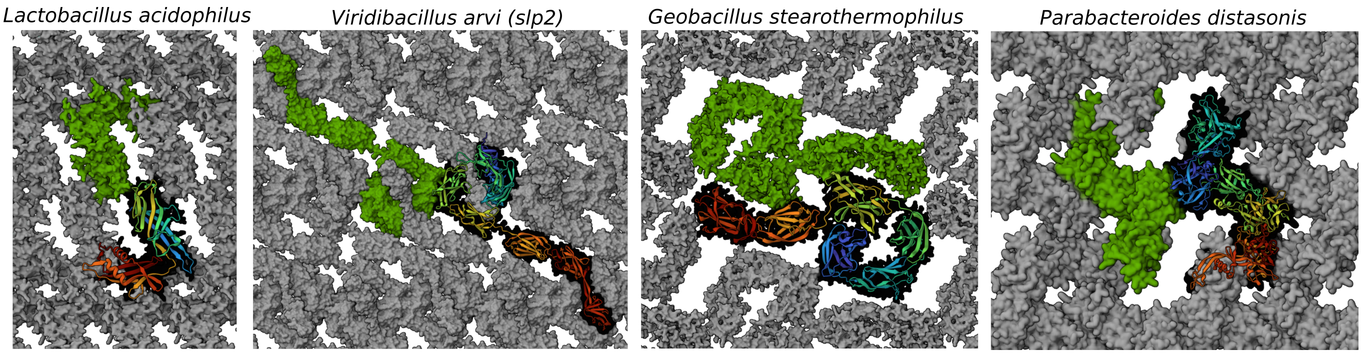


FIG. S20. 2D crystal structure predictions of the $p2$ bacterial S-layers.

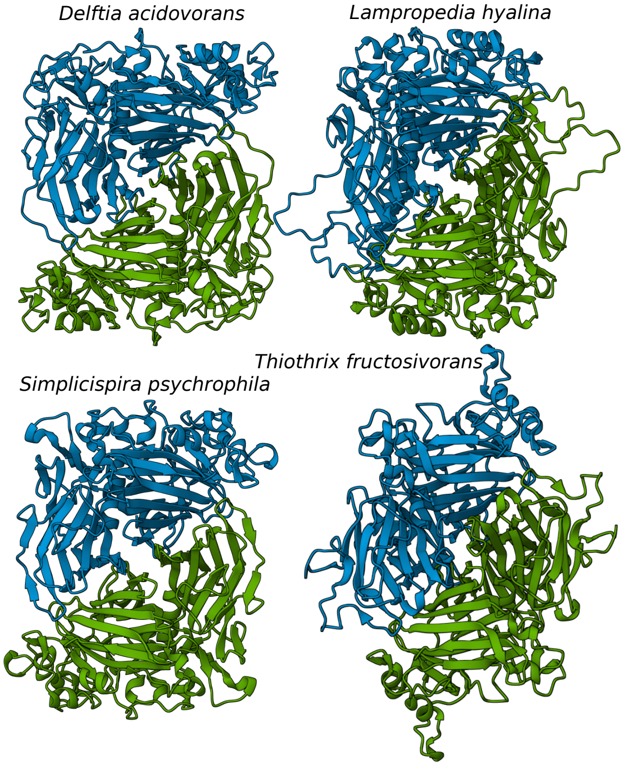


FIG. S21. Dimers of the *Delftia acidovorans* SLP and homologs.

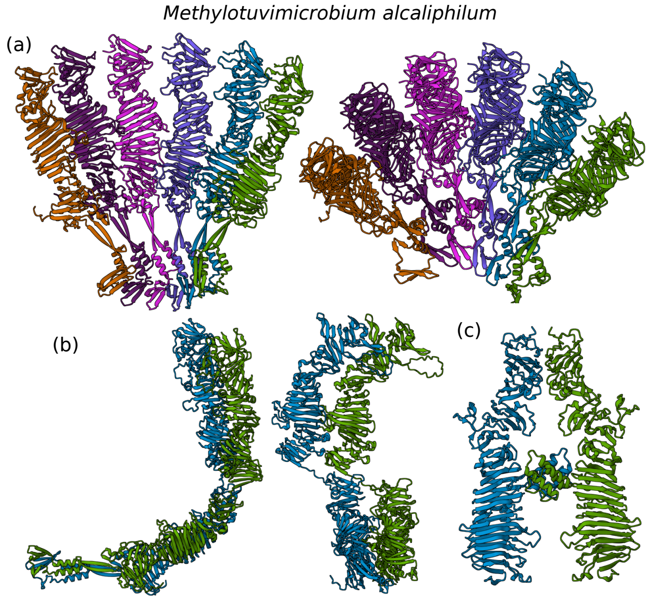


FIG. S22. Assemblies of portions of the *Methylotuvimicrobium alcaliphilum* SLP. (a) Hexamers of first $\beta$-helix domain, (b) dimers of the first two $\beta$-helix domains and (c) a dimer of the end $\beta$-helix domains.

proach has had most success for the $p4$ Bacillota S-layers. These successes can be divided into three architectural classes.

Firstly, we were able to show that the identified SLP of *Aneurinibacillus thermoaerophilus* forms a square pyramidal S-layer, where the protein spirals out in a left-handed fashion from the apex of the pyramid. The protein consists of four assembly domains and an anchoring domain at the C-terminus (Cu_amine_oxidN1). This protein has many homologs in the Bacillota phyla, and as mentioned in the main text, the discovery of the S-layer structure for this example facilitated the definitive identification of the SLP for a series of examples for which the S-layers had been extensively studied by electron microscopy (namely, *Paenibacillus polymyxa*, *Desulfotomaculum nigrificans*, *Clostridium aceticum*, *Thermoanaerobacterium thermosaccharolyticum*, and *Eubacterium yurii*), but for which the SLP was unknown. The predicted S-layers are in excellent agreement with the experimental data both in terms of the lattice constant and comparisons to the EM reconstructions. There is relatively little structural variation between the different examples in this architectural class.

The exemplar for the second $p4$ Bacillota architectural class is SbpA of *Lysinibacillus sphaericus*. SbpA is a large 10-domain protein that traces a complex path through the assembled S-layer. The first four domains are involved in a tetrameric complex with an SLH anchoring domain at its base. The protein then moves parallel to the edge of the unit cell with the fifth domain involved in a homodimeric interaction at the $C_2$ site, before again turning towards the centre of the unit cell with the seventh domain involved in a tetrameric interaction about the second $C_4$ axis. The final three domains are not involved in the assembly but point outwards from the S-layer. Despite SbpA's large size, AlphaFold is able to easily locate the interactions that enable the building of a crystal model. This model is in excellent agreement with an EM map [127] and AFM images [128].

We were also to generate crystal models for homologous proteins for another *Lysinibacillus sphaericus* strain isolated from a uranium mine, *Sporosarcina ureae*, *Alkalihalophilus marmarensis*, and *Viridibacillus arvi*. All have the same basic architecture with the main variations being in the nature of the exterior-pointing domains (e.g. for *Alkalihalophilus marmarensis* they are $\alpha$-helical in character). *Viridibacillus arvi* also has a different anchoring domain (Fig. S8). The *Sporosarcina ureae* crystal prediction is also in excellent agreement with the EM reconstruction of this S-layer [134].

The third type of $p4$ S-layer was found whilst searching for the SLP of *Dehalobacter restrictus*. Curiously, the PER-K23 strain of *Dehalobacter restrictus* has been shown to have a $p6$ S-layer [155], but one of the putative SLPs that we identified, albeit for a different strain, had a clear tendency for tetrameric assembly. Although we were not able to generate a crystal model without significant steric overlap, we were able to generate a better crystal model for the homologous protein in *Desulfosporosinus metallidurans.* The protein consists of seven Ig-like domains and an anchoring domain at the N-terminus. In the monomer, the first and second, and the fifth and sixth Ig-like domains are oriented at approximately 90° to each other. In both cases a simple packing of these L-shaped units gives rise to a clear tendency to tetramer formation. The second tetramer lies somewhat higher in the lattice helping to avoid overlaps between the different monomers. In the case of *Dehalobacter restrictus*, the domains in between the tetramers need to adjust their geometry to achieve this effect.

Intriguingly, SbsC of *Geobacillus stearothermophilus* shows a somewhat similar tendency to tetrameric assembly. This is curious as the relevant strain is able to express two SLPs, SbsC and SbsD, but both were thought to generate oblique lattices [207]. However, for a related strain both $p2$ and $p4$ S-layers have been observed, where SgsE, which has a very high sequence similarity to SbsD, is responsible for the $p2$ S-layer, but the protein responsible for the $p4$ S-layer is unknown [25].

We have generated crystal models for two architectural classes of Bacillota $p6$ S-layers. The first we consider is that for *Thermoanaerobacter thermohydrosulfuricus.* The SLP consists of 4 Ig-like domains plus an anchoring domain at the C-terminus that is of the "Cu_amine_oxidN1" type (i.e. the same as seen for the $p4$ *Aneurinibacillus thermoaerophilus* SLP and homologs). The two C-terminal domains form a hexamer that lies at the base of a hexagonal pyramidal hollow and the two N-terminal domains form a triangular trimer at the exterior face of the S-layer. The AlphaFold predictions are surprisingly good given a relatively shallow MSA and compare very well to the EM reconstruction [159]. Consistent with the shallow MSA, there are relatively few homologs of this SLP, and for the example we tried (*Caldanaerobacter subterraneus*) we were unable to get a complete crystal prediction.

The second type is exemplified by *Thermoanaerobacter kivui.* This SLP is unusual in that consists of OB-barrel domains (Fig. S8), as well as an SLH anchoring domain at the N-terminus. The N-terminal domains form a hexameric barrel. Clear dimeric and trimeric interactions are also located, and although the inter-domain flexibility hinders the generation of a high-quality crystal prediction, there is a clearly a good match between the predictions and the EM results for this S-layer [164]. This SLP has many homologs. The other OB-barrel examples that we considered (*Thermoanaerobacterium saccharolyticum*, *Brevibacillus brevis*, and *Desulfosporosinus lacus* in order of decreasing sequence similarity to *Thermoanaerobacter kivui*) all formed confident hexamers, but it was again difficult to generate good crystal models (e.g. without overlaps) because of the inter-domain flexibility.

As explained in Section S1 D, our approach generally finds it harder to get good crystal predictions for oblique 2D crystals, and these are relatively numerous in the Bacillota phyla. However, we were able to generate three complete crystal predictions. The first was for SgsE of *Geobacillus stearothermophilus.* This protein has six Ig-like domains and an $\alpha$-helical cell-anchoring domain that was first identified for SbsC in a different strain of this bacterium. The N-terminal domains form a 4-domain ring-like structure that is common to many of the $p2$ Bacillota S-layers, followed by a two-domain "arm". The SLP Slp2 of *Viridibacillus arvi* also possesses this 4-domain ring, but has a longer "arm" that protrudes further from the plane of the assembly layer, as well as having a different anchoring domain. We note that SgsE has a 94% similarity to SbsD (also of *Geobacillus stearothermophilus* [207]) and so the latter would be expected to have an almost identical S-layer. We also note that SlpC of *Lysinibacillus sphaericus*, *Acetivibrio thermocellus*, and *Gracilibacter* have SLPs with broadly similar architectures to these two examples.

The third $p2$ prediction was for the SLP of *Lactobacillus acidophilus.* The protein has just three domains with the C-terminal domain responsible for cell anchoring. We were fortunate that for this example predictions for the tetramers and hexamers of the two assembly domains exhibited two-dimensional assemblies with $C_2$ symmetry. We were able to generate a crystal model by combining information from the hexamer with a dimeric interaction involving domain 1. The generated crystal model agrees reasonably with a low-resolution map of the S-layer obtained by electron microscopy [140]; for example, the predicted lattice has $a$=11.2 nm, $b$=3.75 nm and $\gamma$=97.8° compared to $a$=11.8 nm, $b$=5.3 nm and $\gamma$=102°. This suggests that our predictions are somewhat compressed in the $b$-direction. Very recently an all-atom model of the S-layer was generated from a combination of 3D crystal structures, AlphaFold-multimer calculations and fitting to the above EM map [141]. In particular, the inter-domain angles were adjusted to match the above unit cell dimensions and the features in the EM map. The proposed structure involves essentially the same main inter-protein interactions as in our crystal model.

*Bacillus anthracis* Sap represents another type of $p2$ S-layer. We were able to identify a strong homodimeric interaction involving the second assembly domain. An added complexity for this example that is a consequence of the protein's flexibility is that the monomer structure is stabilized by an interaction between the third and sixth assembly domain [150], however in some of the complexes we studied, this same interaction formed intermolecularly rather than intramolecularly. A similar competition between intermolecular and intramolecular interactions (in this case between the fourth and seventh assembly domain) also hindered analysis of the assembly behaviour of SlpA for *Paenibacillus larvae.*

The S-layer of *Corynebacterium glutamicum* was one of the examples highlighted in the main text. We also considered a slightly longer homolog from *Corynebacterium aurimucosum.* It forms a very similar S-layer with almost the same lattice constant, the main difference being the presence of a number of additional disordered loops. This

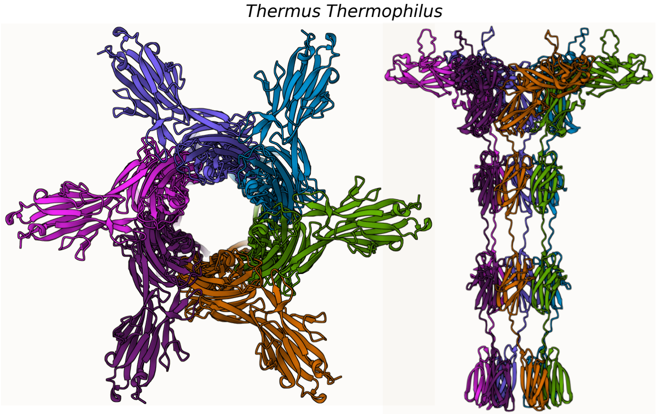


FIG. S23. Hexameric predictions for the first five domains of the *Thermus thermophilus* SLP.

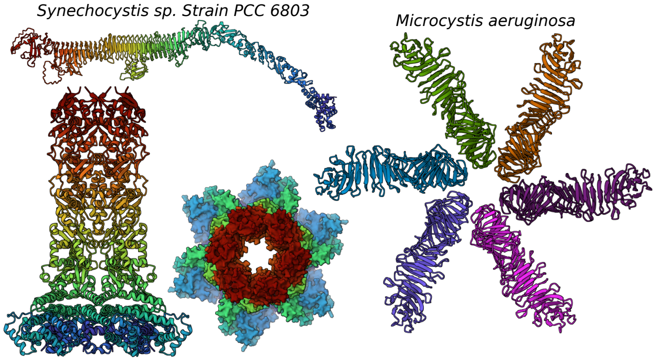


FIG. S24. Confident hexameric predictions for two cyanobacterial SLPs. The hexamer for the *Synechocystis* example is that for the N-terminal half of the SLP and the complete monomer is shown above the hexamer.

type of S-layer forms an isolated cluster in Fig. S6 and seems restricted to a relatively small part of the Actinomycetota phylum.

We also found a putative SLP for *Nocardioides marinus*. It consists of 9 Ig-like domains with the ninth being a likely anchoring domain and the sixth to eighth domains forming a cylindrical hexameric complex.

The structure of the S-layer of *Deinococcus radiodurans* has recently been determined by electron cryomicroscopy [22]. However, the MSA is relatively shallow and, although we were able to obtain a hexameric prediction for the N-terminal domains that matches the experimental structure well, we were not able to locate the dimeric interactions that bridge between the hexameric sites, nor the binding of the C-terminal domains at the top of the hexamer. We had more success for a homolog from *Deinobacterium chartae*, for which we were able to identify both hexameric and dimeric interactions, to build a crystal model.

The S-layer of *Thermus thermophilus* has also been characterized experimentally [171], albeit not at high resolution. It has a noticeably different structure from the S-layer of *Deinococcus radiodurans*, forming a more interwoven hexagonal mesh. Although we were not able to generate a full crystal model for *Thermus thermophilus* (due to the shallowness of the MSA) we were able to generate a crystal model for the homologous protein in *Deinococcus geothermalis* that matches well the experimental images for *Thermus thermophilus*. This consistency makes it very likely that we have identified the correct SLP for *Thermus thermophilus*; note that this differs from previous assignments [208, 209]. The hexameric complex that we were able to identify for this protein is shown in Fig. S23. These two sets of Deinococcota SLPs form nearby interlinked clusters in Fig. S6.

We were able to predict the structures of two types of S-layers in Chloroflexota. Firstly, the SLP has been experimentally identified for *Dehalococcoides mccartyi* [172]. It is an unusual SLP in that it has two $\beta$-propeller domains at its N-terminus; the end $\beta$-propeller domains interact side-on at the $C_3$ site. Our predicted lattice constant (22.1 nm) matches well that measured experimentally (~22 nm) [173].

Secondly, the presence of a $p4$ S-layer has been recently identified for two *Tepidiforma* species [174]. We were able to locate the SLPs by scanning the genome for suitable candidates. The S-layer has a particularly striking 3D porous structure with the protein involved in tetrameric interactions at three different $C_4$ sites at different heights in the S-layer.

An S-layer has also been experimentally observed for *Viridilinea mediisalina* from the class Chloroflexia [210], however we have not as yet been able to identify the SLP for this organism.

Although S-layers of cyanobacteria were one of the better-studied cases by electron microscopy [211, 212], unfortunately, the number of SLPs that have been experimentally identified is relatively few and for none of these examples were we able to generate a full crystal prediction. For *Microcystis aeruginosa* the monomer has an N-terminal cell-anchoring domain followed by a slightly kinked $\beta$-helix; this organisation has significant similarities to *Pseudomonadota* $\beta$-helix SLPs. A very clear tendency to form hexamers driven by interactions at the N-terminal end of the $\beta$-helix was observed (Fig. S24), however we were unable to locate the dimeric or trimeric interactions that would be needed to generate a model of the $p6$ crystal. The identified protein has many homologs and the hexamer is consistent with an EM reconstruction performed for a different *Microcystis aeruginosa* strain [213]. We considered two further homologs in *Microcystis* but were again only able to locate the tendency for hexamer formation.

The identified SLP of *Synechocystis* sp. Strain PCC 6803 is large and has an interesting monomer structure. The N-terminal half formed a hexameric cylindrical tube (Fig. S24) that resembled features observed in an EM reconstruction of a different *Synechocystis* strain [213]. At its N-terminus is a Bluetail putative surface-anchoring domain that likely facilitates anchoring to the cell. However, for the C-terminal half of the protein, which consists of a $\beta$-helix, we were unable to locate any plausible interactions, although this was unsurprising given both the shallowness of the MSA for this protein and its large size.

The S-layers of some unidentified Patescibacteria have been characterized by electron cryomicroscopy and shown to have a $p6$ S-layer consisting of hexagonal pyramids with a lattice constant of 13 nm [214]. By scanning a *Nealsonbacteria* genome, we were able to identify a putative SLP that has a striking resemblance to the SLP of *Haloferax volcanii* in its basic architecture. Both consist of a series of six Ig-like assembly domains with an overall left-hand spiral geometry. The protein showed a very clear tendency to form hexagonal spiral pyramid S-layers, but in this case with the C-terminal domain at the tip of pyramid. Homologs of this protein are particularly numerous across many of the Patescibacterial classes.

A very recent metatranscriptomics study identified the five most highly expressed genes with signal peptides in three closely-related Patescibacterial species [178]. In two of these species, homologs of our identified SLP were among these five proteins, further confirming its S-layer character (we also generated a crystal model for one of these species). In the third species (*Yanofskybacteria bacterium*) the largest protein amongst the five most expressed was a similar protein but with only three Ig-like domains. Furthermore, this species possessed no homologs of the six domain SLP in its genome. We also found that the 3-domain protein could assembly into a hexagonal pyramid S-layer, albeit with a smaller unit cell (13.5 nm rather than 18.5 nm). Given the lattice constant match, it could well be that the experimentally characterised S-layer was for a homolog of this shorter SLP.

#### C. Other lattice-forming proteins

The methods used to predict the 2D lattices of S-layers can also be potentially applied to other 2D lattice-forming proteins. Here we provide some illustrative examples from a variety of classes of proteins. A list of the proteins that we considered is given in Table S5. In their biological context these proteins often form lattices with significant curvature, e.g. virus capsids, but often they are observed to be also able to form 2D sheets *in vitro* under certain conditions [215–217]. For example, the structure of the capsid protein of HIV-1 was first determined by electron crystallography of 2D sheets [215].

In many of these examples, the proteins are significantly smaller than a typical SLP and often consist of a single domain. Like with the SLPs of *Delftia acidovorans* and *Methylomirabilis lanthanidiphila*, it is thus often more difficult to generate sub-proteins that allow the separate identification of the interactions at the different symmetry sites in the lattice. Furthermore, the lattices often assemble hierarchically with strong interactions stabilizing a given oligomer that then further assembles into the lattice through weaker interactions. In such cases, AlphaFold is usually able to identify the dominant complex easily, but it is much harder to identify the secondary interactions. Indeed, aside from MSAs that lack sufficient depth, this was the main reason for the failure to generate a lattice prediction. This was particular the case, where the dominant cluster was a hexamer (or pseudo-hexamer). In these cases, the predictions for dimers or trimers would typically generate a third or half of the hexameric complex, respectively, and the predictions for 12-mers would often generate predictions where two hexamers were directly on top of each other or stacked above each other, rather than a side-by-side configuration. The only examples of success where the hexamer was the dominant oligomer were where the part of the protein responsible for hexamer formation could be excised without disrupting the rest of the monomer structure (e.g. HIV-1 capsid and *Pseudomonas* bacteriophage E217). By contrast, for examples where the dominant cluster was a dimer or a trimer, it was much more likely that the hexamer prediction would be a trimer of dimers or a dimer of trimers, respectively, thus allowing the secondary interactions to be identified.

The successful lattice predictions are depicted in Fig. S25. Most correspond to examples where the structure of the lattices or closed capsules made up of the proteins (either as the sole or dominant component) have already been determined. Agreement is excellent in these cases. Some of the predictions where only the dominant oligomer was identified are illustrated in Fig. S26.

The exosporium proteins represent a particular interesting set of examples because their lattice structures have at best only been determined at low resolution. One family is represented by *Clostridium sporogenes* and *Clostridium pasteurianum.* The predicted lattices are in good agreement with the low-resolution lattice reconstructions obtained by electron microscopy [223]. The lattice formed by the CdeC protein of *Clostridioides difficile* represents a related family and has not been previously characterized. The ExsF and CotY proteins of *Bacillus thuringiensis* represent another class of exosporium proteins. In this case we were only able to identify the dominant hexameric complex. Again this matches well the hexamers that are apparent in the EM reconstructions of the ExsF lattice [222]. A trimer of ExsY has been found to bind to the lattice at the $C_3$ site. Whether this protein is necessary for lattice formation is not clear. Finally, fibres formed by the BclA protein have been observed to extend from these sites; the top of the BclA trimers are shown in Fig. S26.

The sheath proteins of archaea in the classes Methanomicrobia and Methanosarcina represent another interesting example. The proteins for the examples from *Methanospirillum hungatei* and *Methanothrix thermoacetophila* have been experimentally identified. The sheath protein of *Methanothrix soehngenii* that we considered is one of a suite of similar proteins in this organism and is reported for the first time here. They only have extremely weak homology with the former proteins, and so

TABLE S5. Details of other 2D lattice-forming proteins studied. The columns are as for Tables S1 and S2.

| organism | protein name | UniProt ID | length of gene | space group | Refs. | prediction |
|---|---|---|---|---|---|---|
| **Bacterial microcompartments** | | | | | | |
| *Salmonella typhimurium* | pduA | P0A1C7 | 94 | *p*6 | [218, 219] | 6 |
| *Haliangium ochraceum* | BMC-H | D0LID5 | 99 | *p*6 | [218, 219] | 6 |
| *Synechocystis sp. PCC 6803* | ccmK1 | P72760 | 111 | *p*6 | [218, 219] | 6 |
| **Encapsulin nanocompartments** | | | | | | |
| *Mycobacterium tuberculosis* | enc | I6WZG6 | 265 | *p*6 | [220] | 3+2 |
| *Synechococcus elongatus* | enc | Q55032 | 306 | *p*6 | [220] | 6 |
| *Bacillus thermotolerans* | enc | A0A0F5HPP7 | 282 | *p*6 | [220] | 6 |
| *Pyrococcus furiosus* | enc | Q8U1L4 | 345 | *p*6 | [220] | 6 |
| **Exosporia** | | | | | | |
| *Bacillus thuringiensis* | exsY | A0A6L5LKZ9 | 152 | *p*6 | [221, 222] | 6 |
| *Bacillus thuringiensis* | cotY | A0A6L5LL77 | 155 | *p*6 | | 6 |
| *Clostridium sporogenes* | csxA | J7T3V5 | 308 | *p*6 | [223] | 3+2 |
| *Clostridium pasteurianum* | | A0A0H3J2S4 | 262 | *p*6 | [223] | 3+2 |
| *Clostridioides difficile* | CdeC | Q18AS2 | 405 | | | 3+2 |
| **Virus capsids** | | | | | | |
| *HIV-1* | gag | P12493 | 500 | *p*6 | [215, 224, 225] | 6+2 |
| *PBCV-1* | | P30328 | 437 | *p*6 | [226, 227] | 3 |
| *Vaccinia virus* | | P68440 | 551 | *p*6 | [228] | 3+2 |
| *Hepatitis B* | | Q67855 | 185 | *p*6 | [229] | 3+2 |
| *Klebsiella* jumbo myophage $\phi$Kp24 | | A0A7U0GBA8 | 762 | *p*6 | [230] | 6 |
| *Pseudomonas* bacteriophage E217 | | A0A2K8HL59 | 382 | *p*6 | [231] | 6+3 |
| *Q$\beta$* | | Q8LTE1 | 133 | *p*6 | [232] | 3+2 |
| *Bacteriophage MS2* | | P03612 | 130 | *p*6 | [233] | 3+2 |
| *Canine parvovirustype 2* | VP2/VP1 | P17455 | 584/727 | *p*6 | [234] | 3+2 |
| **Virus nuclear shell** | | | | | | |
| *Pseudomonas phage 201$\phi$2-1* | chimallin | B3FIW8 | 631 | *p*2/*p*4 | [235] | 4 |
| **Archaeal/Bacterial sheath proteins** | | | | | | |
| *Methanospirillum hungatei* | | Q2FRN9 | 377 | obl | [236] | |
| *Methanothrix thermoacetophila* | | A0B829 | 575 | obl | [237] | |
| *Methanothrix soehngenii GP-6* | | F4BZP3 | 258 | obl | [238] | |

represents a new class of sheath proteins.

As the sheaths have *p*1 symmetry, we cannot use our approaches (i.e. by aligning the symmetry axes of complexes) to generate a crystal unit cell. However, these examples provide one of the few examples where AlphaFold-multimer calculations directly generate extended assemblies. All the sheath proteins have a $\beta$-sheet character and assemble in one of the two-dimensions through intermolecular $\beta$-sheet formation. This is expected, as these sheath proteins have been identified as an example of functional amyloids [237].

The clearest results are for *Methanospirillum hungatei*. The multimer predictions are first one-dimensional assemblies that propagate the $\beta$-sheet (e.g. the 4-mer in Fig. S27), but for larger multimers two-dimensional assemblies form (e.g. the 12-mer in Fig. S27). The interactions in this second dimension are stabilised by disulphide bridges. The repeats in both directions match well that observed experimentally [238]. Interestingly, the assemblies exhibit a curvature in this second dimension that breaks perfect translational periodicity in this dimension and may explain the longer length-scale striations observed in this direction in experiments and the dissolution of the sheath into loops [239]. In this picture the weakened lateral bonding would be associated with a regular resetting of the orientations of the amyloid fibres between uniformly curved sections. Very recently, cryoEM (supplemented by fitting the AlphaFold monomer into the EM density) has revealed the structure of the sheaths in detail, confirming the basic lattice revealed by AlphaFold-multimer and the above suggestion for the origin of the striations [240]

For the *Methanothrix soehngenii* example, we also find 2-dimensional assemblies. However, the MSA for the *Methanothrix thermoacetophila* sheath protein is too shallow to observe significant assembly. Its propensity for amyloid formation is clear from the monomer structure (Fig. S27).

### S4. FURTHER RESULTS: BINDING MECHANISMS

Bioinformatic analysis of SLP domain composition (described in the Methods section above), allowed us to detect significant differences in the level of diversity present

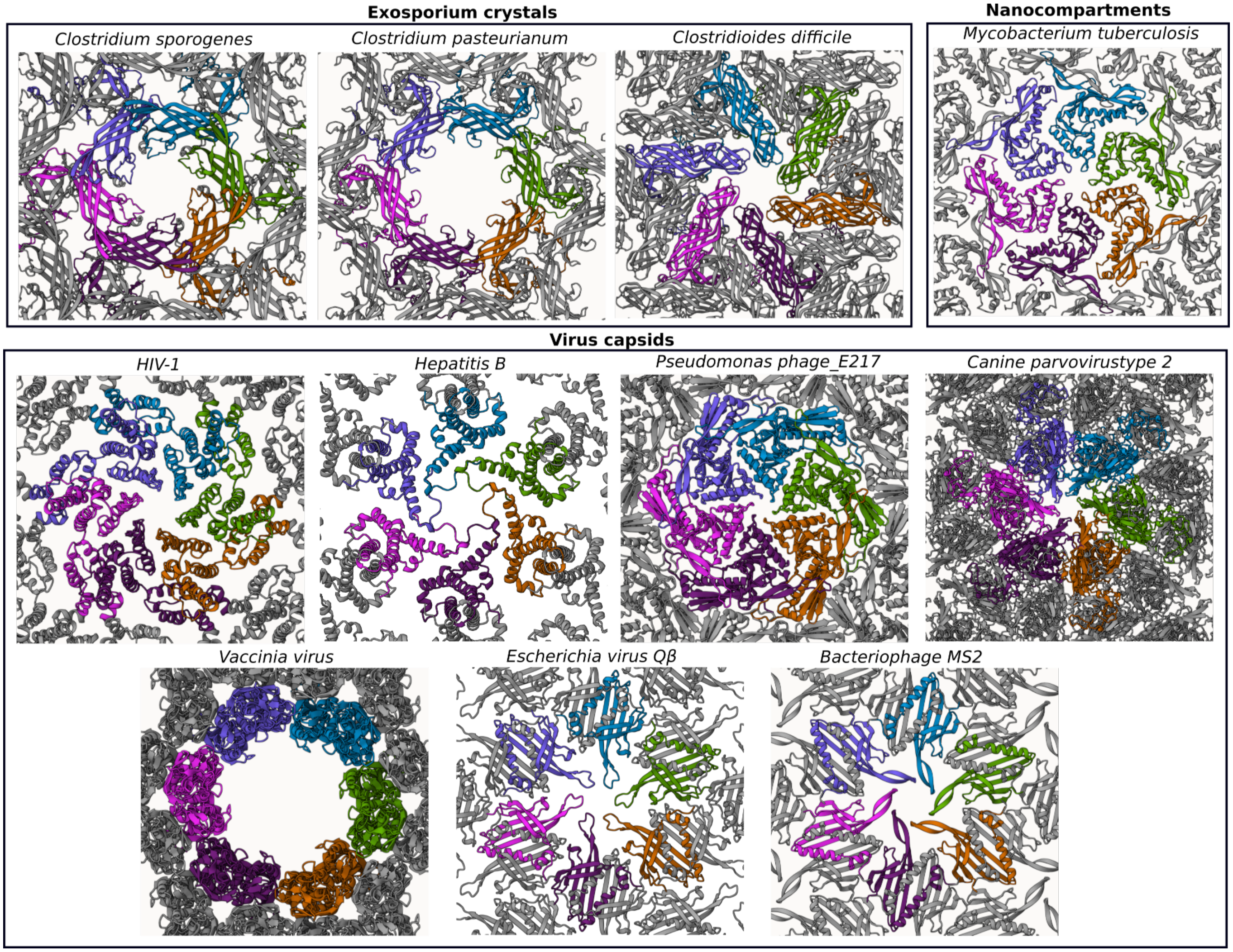


FIG. S25. Predicted 2D crystal structures of lattice-forming proteins not belonging to the S-layer class.

in bacterial and archaeal S-layers. Summarised in Fig. 5 of the main text, our analysis shows that, compared to archaeal S-layers, bacterial S-layers exhibit remarkable diversity in their anchoring mechanisms. There are far fewer anchoring mechanisms detected in archaea (Fig. S7), compared with bacteria (Fig. S8). In most archaea, S-layers are directly anchored to the cell membrane through lipidation, transmembrane helices, membrane-binding domains, or interactions with co-expressed accessory proteins. In contrast, bacterial S-layers can be anchored to various components of the cell envelope, including the cell membrane, peptidoglycan or lipopolysaccharide. Specifically, bacterial SLPs employ a variety of anchoring mechanisms, including lipidation [22], PG-binding or secondary cell wall polymer (SCWP)-binding domains [194, 241], and transmembrane $\alpha$-helices [21]. This diversity in anchoring mechanisms is reflected in the structural diversity of protein domains mediating anchoring, with only a few types observed in archaea, compared with myriad of anchoring domains in bacteria (Fig. S28). Please see the main text for a detailed discussion on the implications of these differences between bacteria and archaea.

### S5. LIST OF SUPPLEMENTARY MOVIES

Movie S1. Cryo-ET of the *C. glutamicum* S-layer. Sequential Z-slices of a tomogram of the *C. glutamicum* S-layer showing the hexagonally arranged lattice (scale bar= 100 nm).

[1] R. Evans, M. O'Neill, A. Pritzel, N. Antropova, A. Senior, T. Green, A. Žídek, R. Bates, S. Blackwell, J. Yim, O. Ronneberger, S. Bodenstein, M. Zielinski,

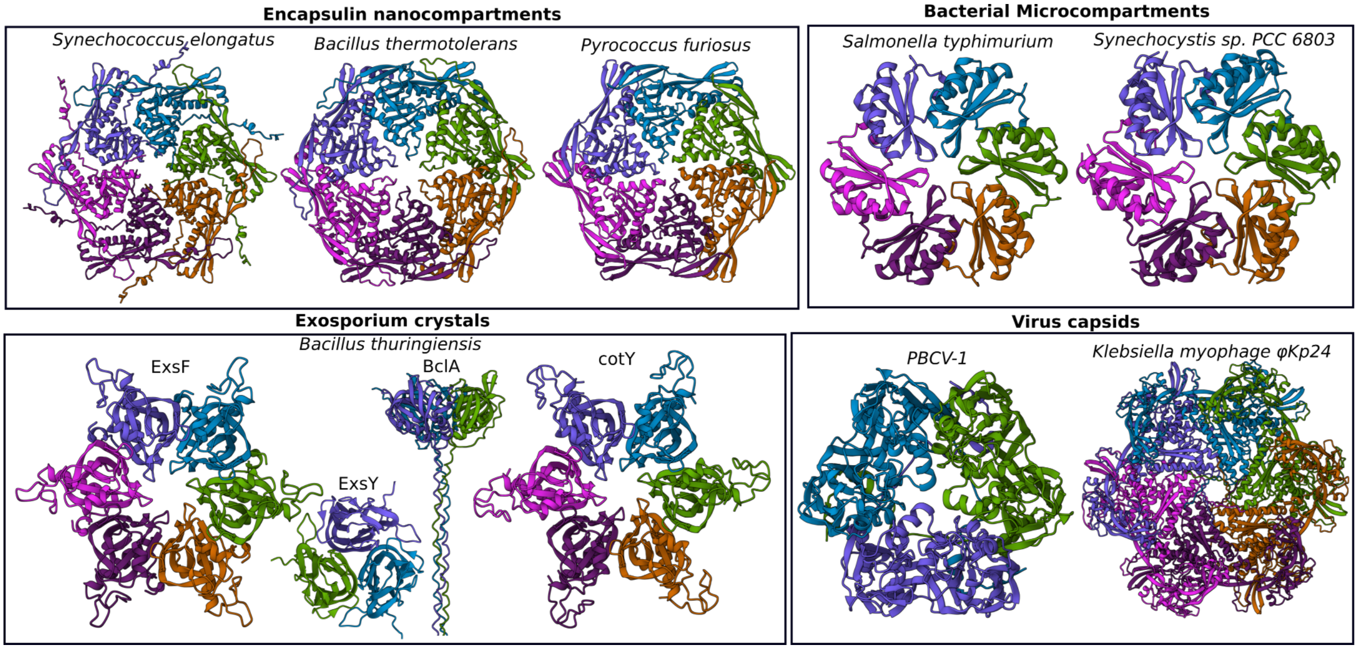


FIG. S26. Oligomer predictions for lattice-forming proteins not belonging to the S-layer class.

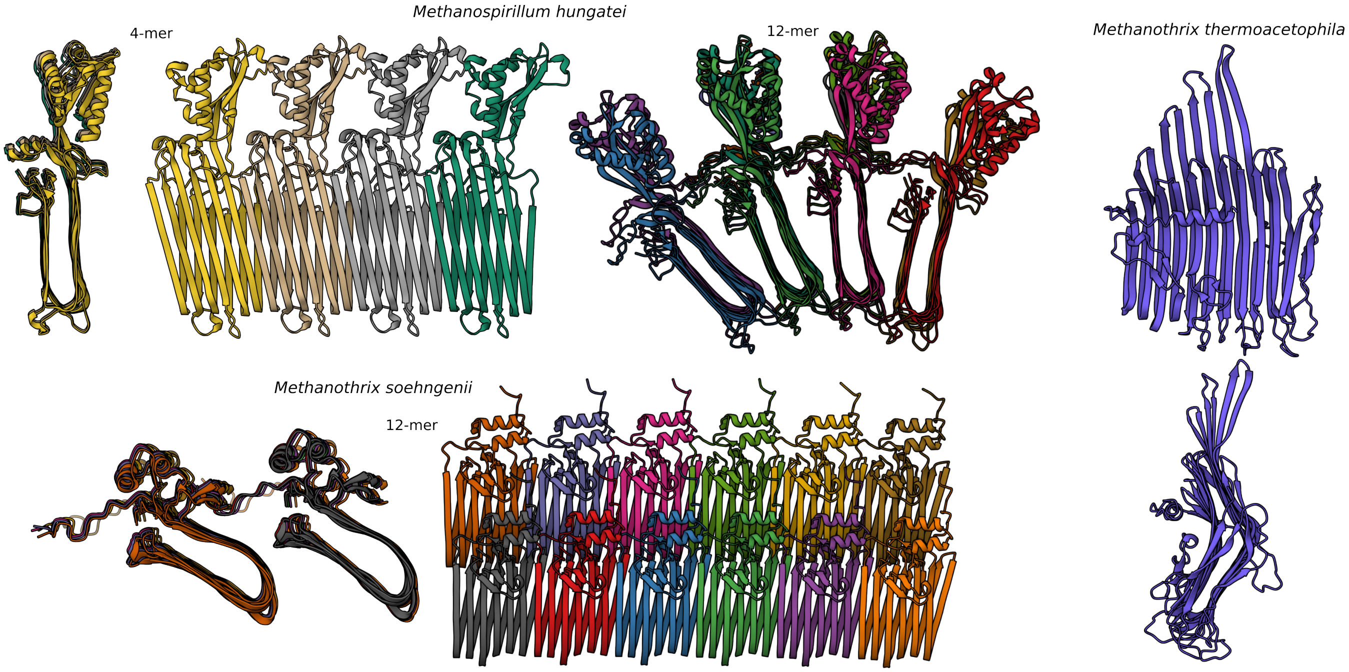


FIG. S27. Sheath proteins and their assemblies

A. Bridgland, A. Potapenko, K. Cowie, A. Tunyasuvunakool, R. Jain, E. Clancy, P. Kohli, J. Jumper, and D. Hassabis, Protein complex prediction with alphafold-multimer, bioRxiv , 2021.10.04.463034.

[2] M. Mirdita, K. Schütze, Y. Moriwaki, L. Heo, S. Ovchinnikov, and M. Steinegger, Colabfold: making protein folding accessible to all, Nature Methods **19**, 679 (2022).

[3] M. Steinegger and J. Söding, MMseqs2 enables sensitive protein sequence searching for the analysis of massive data sets, Nat. Biotechnol. **35**, 1026 (2017).

[4] J. P. Roney and S. Ovchinnikov, State-of-the-art estimation of protein model accuracy using AlphaFold, Phys. Rev. Lett. **129**, 238101 (2022).

[5] L. S. Johnson, S. R. Eddy, and E. Portugaly, Hidden Markov model speed heuristic and iterative HMM

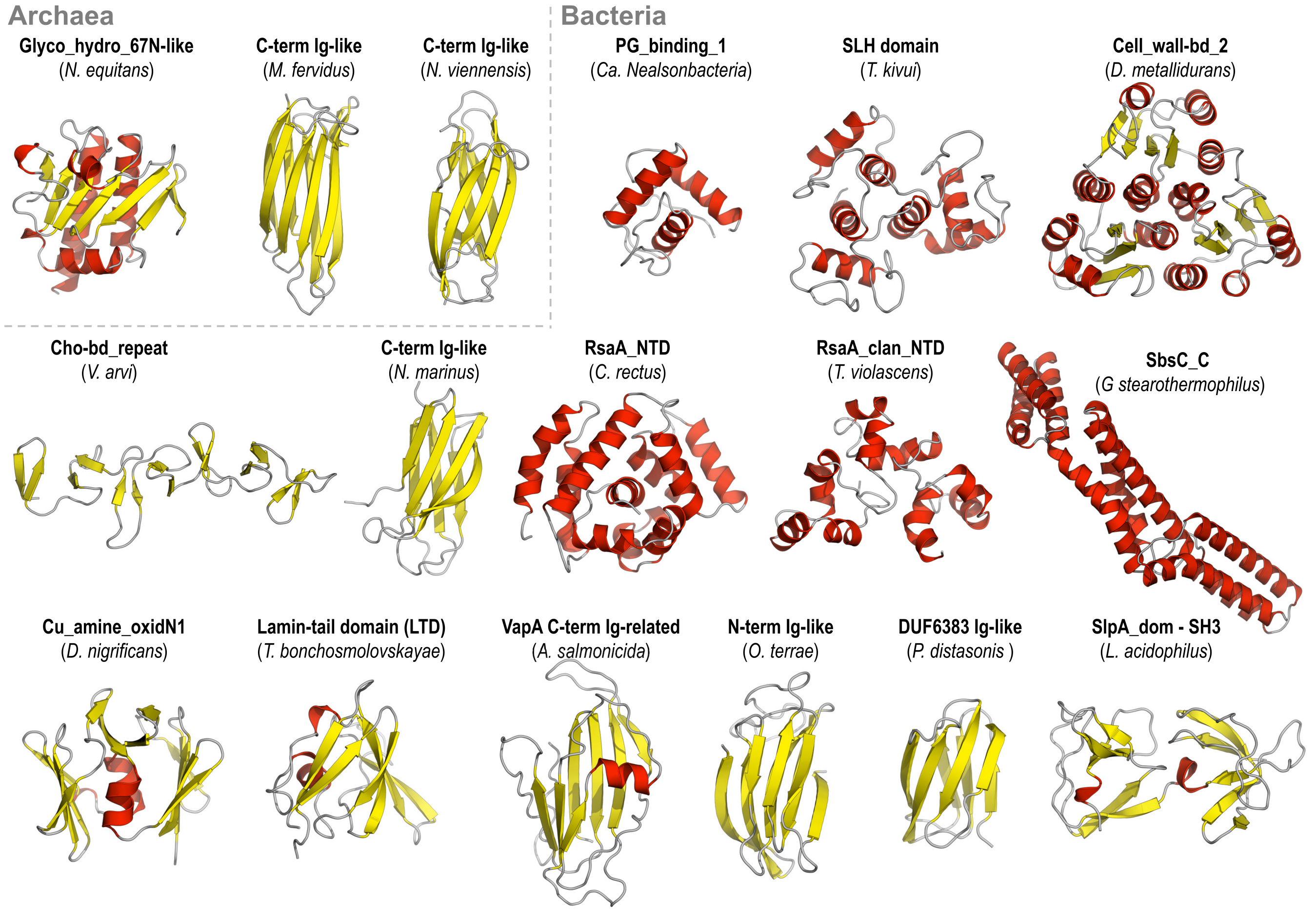


FIG. S28. Structural gallery of anchoring domains found in archaeal and bacterial SLPs. Representative structures of anchoring domains characterized in this study, including those previously known, are shown.


search procedure, BMC Bionformatics **11**, 431 (2010).

[6] J. Jumper, R. Evans, A. Pritzel, T. Green, M. Figurnov, O. Ronneberger, K. Tunyasuvunakool, R. Bates, A. Žídek, A. Potapenko, A. Bridgland, C. Meyer, S. A. A. Kohl, A. J. Ballard, A. Cowie, B. Romera-Paredes, S. Nikolov, R. Jain, J. Adler, T. Back, S. Petersen, D. Reiman, E. Clancy, M. Zielinski, M. Steinegger, M. Pacholska, T. Berghammer, S. Bodenstein, D. Silver, O. Vinyals, K. Kavukcuoglu, P. Kohli, and D. Hassabis, Highly accurate protein structure prediction with alphafold, Nature **596**, 583 (2021).

[7] D. Pum, J. L. Toca-Herrera, and U. B. Sleytr, Patterns in nature—S-layer lattices of bacterial and archaeal cells, Crystals **11**, 869 (2021).

[8] A. von Kügelgen, V. Alva, and T. A. M. Bharat, Complete atomic structure of a native archaeal cell surface, Cell Rep. **37**, 110052 (2021).

[9] O. Pornillos, B. K. Ganser-Pornillos, and M. Yeager, Atomic-level modelling of the HIV capsid, Nature **469**, 424 (2011).

[10] W. O. Saxton and W. Baumeister, Principles of organization in S layers, J. Mol. Biol **187**, 251 (1986).

[11] T. A. M. Bharat, D. Kureisaite-Ciziene, G. G. Hardy, E. W. Yu, J. M. Devant, W. J. H. Hagen, Y. V. Brun, J. A. G. Briggs, and J. Löwe, Structure of the hexagonal surface layer on *Caulobacter crescentus* cells, Nat. Microbiol. **2**, 17059 (2017).

[12] E. C. Meng, E. F. Pettersen, G. S. Couch, C. C. Huang, and T. E. Ferrin, Tools for integrated sequence-structure analysis with UCSF Chimera, BMC Bioinformatics **7**, 339 (2006).

[13] D. Sehnal, S. Bittrich, M. Deshpande, R. Svobodová, K. Berka, V. Bazgier, S. Velankar, S. K. Burley, J. Koča, and A. S. Rose, Mol* Viewer: modern web app for 3D visualization and analysis of large biomolecular structures, Nucleic Acids Res. **49**, W431 (2021).

[14] E. Baranova, R. Fronzes, A. Garcia-Pino, N. Van Gerven, D. Papapostolou, G. Péhau-Arnaudet, E. Pardon, J. Steyaert, S. Howorka, and H. Remaut, SbsB structure and lattice reconstruction unveil $Ca^{2+}$ triggered S-layer assembly, Nature **487**, 119 (2012).

[15] A. Sogues, A. Fioravanti, W. Jonckheere, E. Pardon, J. Steyaert, and H. Remaut, Structure and function of the EA1 surface layer of *Bacillus anthracis*, Nat. Commun. **14**, 7051 (2023).

[16] M. Stewart, T. J. Beveridge, and T. J. Trust, Two patterns in the *Aeromonas salmonicida* A-layer may reflect a structural transformation that alters permeability, J. Bacteriol. **166**, 120 (1986).

[17] P. Delepelaire and C. Wandersman, Protein secretion in gram-negative bacteria. The extracellular metalloprotease B from *Erwinia chrysanthemi* contains a C-terminal secretion signal analogous to that of *Escherichia coli* alpha-hemolysin, J. Biol. Chem. **265**, 17118 (1990).

[18] S. A. Thompson, O. L. Shedd, K. C. Ray, M. H. Beins, J. P. Jorgensen, and M. J. Blaser, *Campylobacter fetus* surface layer proteins are transported by a type I secretion system, J. Bacteriol. **180**, 6450 (1998).

[19] T. A. M. Bharat, A. von Kügelgen, and V. Alva, Molecular logic of prokaryotic surface layer structures, Trends Microbiol. **29**, 405 (2021).

[20] M. Abdul-Halim, S. Schulze, A. DiLucido, F. Pfeiffer, A. W. Bisson Fijho, and M. Pohlschroder, Lipid anchoring of archaeosortase substrates and midcell growth in haloarchaea, mBio **11**, e00349 (2020).

[21] L. Gambelli, B. H. Meyer, M. McLaren, K. Sanders, T. E. F. Quax, V. Gold, S.-V. Albers, and B. Daum, Architecture and modular assembly of *Sulfolobus* S-layers revealed by electron cryotomography, Proc. Natl. Acad. Sci. USA **116**, 25278 (2019).

[22] A. von Kügelgen, S. van Dorst, K. Yamashita, D. L. Sexton, E. I. Tocheva, G. Murshudov, V. Alva, and T. A. M. Bharat, Interdigitated immunoglobulin arrays form the hyperstable surface layer of the extremophilic bacterium *Deinococcus radiodurans*, Proc. Natl. Acad. Sci. USA **120**, e2215808120 (2023).

[23] M. Taketani, M. S. Donia, A. N. Jacobson, J. D. Lambris, and M. A. Fischbach, A phase-variable surface layer from the gut symbiont *Bacteroides thetaiotaomicron*, mBio **6**, e01339 (2015).

[24] E. Couture-Tosi, H. Delacroix, T. Mignot, S. Mesnage, M. Chami, A. Fouet, and G. Mosser, Structural analysis and evidence for dynamic emergence of *Bacillus anthracis* S-layer networks, J. Bacteriol. **184**, 6448 (2002).

[25] M. Sára, D. Pum, S. Küpcü, P. Messner, and U. B. Sleytr, Isolation of two physiologically induced variant strains of *Bacillus stearothermophilus* NRS 2004/3a and characterization of their S-layer lattices, J. Bacteriol. **176**, 848 (1994).

[26] M. R. Tummuru and M. J. Blaser, Rearrangement of *sapA* homologs with conserved and variable regions in *Campylobacter fetus*, Proc. Natl. Acad. Sci. USA **90**, 7265 (1993).

[27] C. Mercier, D. Thies, L. Zhong, M. J. Raftery, and S. Erdmann, Characterization of an archaeal virus-host system reveals massive genomic rearrangements in a laboratory strain, Front. Microbiol. **14**, 1274068 (2023).

[28] L. Gambelli, M. McLaren, R. Conners, K. Sanders, M. C. Gaines, L. Clark, V. A. M. Gold, D. Kattnig, M. Sikora, C. Hanus, M. N. Isupov, and B. Daum, Structure of the two-component S-layer of the archaeon *Sulfolobus acidocaldarius*, eLife **13**, e84617 (2024).

[29] M. Sumper, E. Berg, R. Mengele, and I. Strobel, Primary structure and glycosylation of the S-layer protein of *Haloferax volcanii*, J. Bacteriol. **172**, 7111 (1990).

[30] M. Kessel, I. Wildhaber, S. Cohen, and W. Baumeister, Three-dimensional structure of the regular surface glycoprotein layer of *Halobacterium volcanii* from the Dead Sea, EMBO J. **7**, 1549 (1988).

[31] H. Wakai, S. Nakamura, H. Kawasaki, K. Takada, S. Mizutani, R. Aono, and K. Horikoshi, Cloning and sequencing of the gene encoding the cell surface glycoprotein of *Haloarcula japonica* strain TR-1, Extremophiles **1**, 29 (1997).

[32] Y. Nishiyama, T. Takashina, W. D. Grant, and K. Horikoshi, Ultrastructure of the cell wall of the triangular halophilic archaebacterium *Haloarcula japonica* strain TR-1, FEMS Microbiol. Lett. **99**, 43 (1992).

[33] J. Lechner and M. Sumper, The primary structure of a procaryotic glycoprotein: Cloning and sequencing of the cell surface glycoprotein gene of halobacteria, J. Biol. Chem. **262**, 9724 (1987).

[34] S. Trachtenberg, B. Pinnick, and M. Kessel, The cell surface glycoprotein layer of the extreme halophile *Halobacterium salinarum* and its relation to *Haloferax volcanii*: Cryo-electron tomography of freeze-substituted cells and projection studies of negatively stained envelopes, J. Struct. Biol. **130**, 10 (2000).

[35] P. G. Bolhuis, A. B. Martín-Cuadrado, R. Rosselli, L. Pašić, and F. Rodriguez-Varela, Transcriptome analysis of *Haloquadratum walsbyi*: vanity is but the surface, BMC Genom. **18**, 510 (2017).

[36] D. G. Burns, P. H. Janssen, T. Itoh, M. Kamekura, Z. Li, G. Jensen, F. Rodríguez-Valera, H. Bolhuis, and M. L. Dyall-Smith, *Haloquadratum walsbyi* gen. nov., sp. nov., the square haloarchaeon of Walsby, isolated from saltern crystallizers in Australia and Spain, Int. J. Syst. Evol. Microbiol. **57**, 387 (2007).

[37] T. J. Williams, Y. Liao, J. Ye, R. P. Kuchel, A. Poljak, M. J. Raferty, and R. Cavicchioli, Cold adaptation of the antarctic haloarchaea *Halohasta litchfieldiae* and *Halorubrum lacusprofundi*, Environ. Microbiol. **19**, 2210 (2017).

[38] H. Lu, Y. Lü, J. Ren, Z. Wang, Q. Wang, Y. Luo, J. Han, H. Xiang, Y. Du, and C. Jin, Identification of the S-layer glycoproteins and their covalently linked glycans in the halophilic archaeon *Haloarcula hispanica*, Glycobiology **25**, 1150 (2015).

[39] D. H. Haft, S. H. Payne, and J. D. Selengut, Archaeosortases and exosortases are widely distributed systems linking membrane transit with posttranslational modification, J. Bacteriol. **194**, 36 (2012).

[40] J. Kelly, E. Vinogradov, A. Robotham, L. Tessier, S. M. Logan, and K. F. Jarrell, Characterizing the N- and O-linked glycans of the PGF-CTERM sorting domain-containing S-layer protein of *Methanoculleus marisnigri*, Glycobiology **32**, 629 (2022).

[41] D. P. Bayler and S. F. Koval, Membrane association and isolation of the S-layer protein of *Methanoculleus marisnigri*, Can. J. Microbiol. **40**, 237 (1994).

[42] M. Firtel, G. Southam, G. Harauz, and T. J. Beveridge, Characterization of the cell wall of the sheathed methanogen *Methanospirillum hungatei* GP1 as an S layer, J. Bacteriol. **175**, 7550 (1993).

[43] G.-W. Cheong, D. Typke, and W. Baumeister, The surface protein layer of *Methanoplanus limicola*: Three-dimensional structure and chemical characterization, System Appl. Microbiol. **14**, 209 (1991).

[44] G.-W. Cheong, D. Typke, and W. Baumeister, Projection structure of the surface layer of *Methanoplanus limicola* at 10 Å resolution obtained by electron cryomicroscopy, J. Struct. Biol. **117**, 138 (1996).

[45] G. Zellner, E. Stackebrandt, B. J. Tindall, E. Conway de Macario, H. Kneifel, and U. B. Sleytr, *Methanocorpusculaceae* fam. nov., represented by *Methanocorpusculum parvum*, *Methanocorpusculum sinense* spec. nov. and

*Methanocorpusculum bavaricum* spec. nov., Arch. Microbiol. **151**, 381 (1989).

[46] G. Zellner, P. Messner, H. Kneifel, B. J. Tindall, J. Winter, and E. Stackebrandt, *Methanolacinia* gen. nov., incorporating *Methanomicrobium paynteri* as *Methanolacinia paynteri* comb. nov., J. Gen. Appl. Microbiol. **35**, 185 (1989).

[47] H. P. Zabel, H. König, and J. Winter, Isolation and characterization of a new coccoid methanogen, *Methanogenium tatii* spec. nov. from a solfataric field on Mount Tatio, Arch. Microbiol. **137**, 308 (1984).

[48] H. Huber, H. Jannasch, R. Rachel, T. Fuchs, and K. O. Stetter, *Archaeoglobus veneficus sp.* nov., a novel facultative chemolithoautotrophic hyperthermophilic sulfite reducer, isolated from abyssal black smokers, System Appl. Microbiol. **20**, 374 (1997).

[49] D. Hafenbradl, M. Keller, R. Dirmeier, R. Rachel, P. Roßnagel, S. Burggraf, H. Huber, and K. O. Stetter, *Ferroglobus placidus* gen. nov., sp. nov., a novel hyperthermophilic archaeum that oxidizes $Fe^{2+}$ at neutral pH under anoxic conditions of liquids and amorphous metals and alloys, Arch. Microbiol. **166**, 308 (1996).

[50] S. Sakai, R. Conrad, W. Liesack, and H. Imachi, *Methanocella arvoryzae* sp. nov., a hydrogenotrophic methanogen isolated from rice field soil, Int. J. Syst. Evol. Microbiol. **60**, 2918 (2010).

[51] D. R. Francoleon, P. Boontheung, Y. Yang, U. Kim, A. J. Ytterberg, P. A. Denny, P. C. Denny, J. A. Loo, R. P. Gunsalus, and R. R. Ogorzalek Loo, S-layer, surface-accessible, and Concanavalin A binding proteins of *Methanosarcina acetivorans* and *Methanosarcina mazei*, J. Proteome Res. **8**, 1972 (2009).

[52] M. A. Arbing, S. Chan, A. Shin, T. Phan, C. J. Ahn, L. Rohlin, and R. P. Gunsulas, Structure of the surface layer of the methanogenic archaean *Methanosarcina acetivorans*, Proc. Natl. Acad. Sci. USA **109**, 11812 (2012).

[53] G. Bröckl, M. Behr, S. Fabry, R. Hensel, H. Kaudewitz, E. Biendl, and H. König, Analysis and nucleotide sequence of the genes encoding the surface-layer glycoproteins of the hyperthermophilic methanogens *Methanothermus fervidus* and *Methanothermus sociabilis*, Eur. J. Biochem. **199**, 147 (1991).

[54] H. König, R. Rachel, and H. Claus, Proteinaceous surface layers of *Archaea*: Ultrastructure and biochemistry, in *Archaea: Molecular and Cellular Biology*, edited by R. Cavicchioli (ASM Press, 2007) pp. 315–340.

[55] J. Konisky, D. Lynn, M. Hoppert, F. Mayer, and P. Haney, Identification of the *Methanococcus voltae* S-layer structural gene, J. Bacteriol. **176**, 1790 (1994).

[56] S. F. Koval and K. F. Jarrell, Ultrastructure and biochemistry of the cell wall of *Methanococcus voltae*, J. Bacteriol. **169**, 1298 (1987).

[57] E. Akca, H. Claus, N. Schultz, G. Karbach, B. Schlott, T. Debaerdemaeker, J.-P. Declercq, and H. König, Genes and derived amino acid sequences of S-layer proteins from mesophilic, thermophilic, and extremely thermophilic methanococci, Extremophiles **6**, 351 (2002).

[58] E. Nußer and H. König, S layer studies on three species of *Methanococcus* living at different temperatures, Can. J. Microbiol. **33**, 256 (1987).

[59] A. S. Kostyukova, G. M. Gongadze, Y. Y. Polosina, E. A. Bonch-Osmolovskaya, M. L. Miroshnichenko, N. A. Chernyh, M. V. Obraztsova, V. A. Svetlichny, P. Messner, U. B. Sleytr, S. L'Haridon, C. Jeanthon, and D. Prieur, Investigation of structure and antigenic capacities of *Thermococcales* cell envelopes and reclassification of "*Caldococcus litoralis*" Z-1301 as *Thermococcus litoralis* Z-1301, Extremophiles **3**, 239 (1999).

[60] S. Goda, T. Koga, K. Yamashita, R. Kuriura, and T. Ueda, A novel carbohydrate-binding surface layer protein from the hyperthermophilic archaeon *Pyrococcus horikoshii*, Biosci. Biotechnol. Biochem. **82**, 1327 (2018).

[61] Y. Moalic, J. Hartunians, C. Dalmasso, D. Courtine, M. Georges, P. Oger, Z. Shao, M. Jebbar, and K. Alain, The piezo-hyperthermophilic archaeon *Thermococcus piezophilus* regulates its energy efficiency system to cope with large hydrostatic pressure variations, Front. Microbiol. **12**, 730231 (2021).

[62] N. Singhal, A. Garg, N. Singh, P. Gulati, M. Kumar, and M. Goel, Efficacy of signal peptide predictors in identifying signal peptides in the experimental secretome of *Picrophilous torridus*, a thermoacidophilic archaeon, PLoS One **16**, e0255826 (2021).

[63] C. Schlper, G. Puehler, I. Holz, A. Gambacorta, D. Janekovic, U. Santarius, H.-P. Klenk, and W. Zillig, *Picrophilus* gen. nov., fam. nov.: a novel aerobic, heterotrophic, thermoacidophilic genus and family comprising archaea capable of growth around pH 0, J. Bacteriol. **177**, 7050 (1995).

[64] W. Qin, S. A. Amin, R. A. Lundeen, K. R. Heal, W. Martens-Habbena, S. Turkarslan, H. Urakawa, K. C. Costa, E. L. Hendrickson, T. Wang, D. A. C. Beck, S. M. Tiquia-Arashiro, F. Taub, A. D. Holmes, N. Vajrala, P. M. Berube, T. M. Lowe, J. W. Moffett, A. H. Devol, N. S. Baliga, D. J. Arp, L. A. Sayavedra-Soto, M. Hackett, E. V. Armbrust, A. E. Ingalls, and D. A. Stahl, Stress response of a marine ammonia-oxidizing archaeon informs physiological status of environmental populations, ISME J. **12**, 508 (2018).

[65] W. Qin, K. R. Heal, R. Ramdasi, J. N. Kobelt, W. Martens-Habbena, A. D. Bertgnolli, S. A. Amin, C. B. Walker, H. Urakawa, M. Könneke, A. H. Devol, J. W. Moffett, E. V. Armbrust, G. J. Jensen, A. E. Ingalls, and D. A. Stahl, *Nitrosopumilus maritimus* gen. nov., sp. nov., *Nitrosopumilus cobalaminigenes* sp. nov., *Nitrosopumilus oxyclinae* sp. nov., and *Nitrosopumilus ureiphilus* sp. nov., four marine ammonia-oxidizing archaea of the phylum *Thaumarchaeota*, Int. J. Syst. Evol. Microbiol. **67**, 5067 (2017).

[66] M. Kerou, P. Offre, L. Valledor, S. S. Abby, M. Melcher, M. Nagler, W. Weckwerth, and C. Schleper, Proteomics and comparative genomics of *Nitrososphaera viennensis* reveal the core genome and adaptations of archaeal ammonia oxidizers, Proc. Natl. Acad. Sci. USA **113**, E7937 (2016).

[67] L. H. Hodgskiss, M. Melcher, M. Kerou, W. Chen, R. I. Ponce-Toledo, S. N. Savvides, S. Wienkoop, M. Hartl, and C. Schleper, Unexpected complexity of the ammonia monooxygenase in archaea, ISME J. **17**, 588 (2023).

[68] M. Stieglmeier, A. Klingl, R. J. E. Alves, S. K.-M. R. Rittmann, M. Melcher, N. Leisch, and C. Schleper, *Nitrososphaera viennensis* gen. nov., sp. nov., an aerobic and mesophilic, ammonia-oxidizing archaeon from soil and a member of the archaeal phylum *Thaumarchaeota*, ISME J. **64**, 2738 (2014).

[69] A. Klingl, C. Pickl, and J. Fleschsler, Archaeal cell walls, in *Bacterial Cell Walls and Membranes*, Subcellular Biochemistry, Vol. 92, edited by A. Kuhn (Springer, 2019) pp. 471–111.

[70] I. Wildhaber, U. Santarius, and W. Baumeister, Three-dimensional structure of the surface protein of *Desulfurococcus mobilis*, J. Bacteriol. **169**, 5563 (1987).

[71] J. Peters, W. Baumeister, and A. Lupas, Hyperthermostable surface layer protein tetrabrachion from the archaebacterium *Staphylothermus marinus*: Evidence for the presence of a right-handed coiled coil derived from the primary structure, J. Mol. Biol. **257**, 1031 (1996).

[72] J. Peters, M. Nitsch, B. Kühlmorgen, R. Golbik, A. Lupas, J. Kellermann, H. Engelhardt, J.-P. Pfander, S. Müller, K. Goldie, A. Engel, K.-O. Stetter, and W. Baumeister, Tetrabrachion: A filamentous archaebacterial surface protein assembly of unusual structure and extreme stability, J. Mol. Biol. **245**, 385 (1995).

[73] G. Palmieri, R. Cannio, I. Fiume, M. Rossi, and G. Pocsfalvi, Outside the unusual cell wall of the hyperthermophilic archaeon *Aeropyrum pernix* K1, Mol. Cell. Proteomics **8**, 2570 (2009).

[74] R. Rachel, Cell envelopes of crenarchaeota and nanoarchaeota, in *Prokaryotic cell wall compounds*, edited by H. König, H. Claus, and A. Varma (Springer-Verlag, Berlin, 2010) pp. 271–291.

[75] W. Baumeister, U. Santarius, S. Volker, G. Lembcke, and H. Engelhardt, The surface protein of *Hyperthermus butylicus*: Three dimensional structure and comparison with other archaebacterial surface proteins, System Appl. Microbiol. **13**, 105 (1990).

[76] R. Hegerl and W. Baumeister, Correlation averaging of a badly distorted lattice: The surface protein of *Pyrodictium occultum*, J. Electron Microsc. Tech. **8**, 413 (1988).

[77] A. Veith, A. Klingl, B. Zolghadr, K. Lauber, R. Mentele, F. Lottspeich, R. Rachel, S.-V. Albers, and A. Kletzin, *Acidianus*, *Sulfolobus* and *Metallosphaera* surface layers: structure, composition and gene expression, Mol. Microbiol. **73**, 58 (2009).

[78] S. Kato, Y. O. Tahara, T. Nishimura, K. Uematsu, T. Arai, D. Nakane, A. Ihara, T. Nishizaka, W. Iwasaki, T. Itoh, M. Miyata, and M. Ohkuma, Cell surface architecture of the cultivated DPANN archaeon *Nanobdella aerobiophila*, J. Bacteriol. **206**, e00351 (2024).

[79] H. Huber, C. Spinnler, A. Gambacorta, and K. O. Stetter, *Metallosphaera sedula* gen. and sp. nov. represents a new genus of aerobic, metal-mobilizing, thermoacidophilic archaebacteria, System Appl. Microbiol. **12**, 38 (1989).

[80] W. Baumeister, S. Volker, and U. Santarius, The three-dimensional structure of the surface protein of *Acidianus brierleyi* determined by electron crystallography, System Appl. Microbiol. **14**, 103 (1991).

[81] R. Prüschenk, W. Baumeister, and W. Zillig, Surface structure variants in different species of *Sulfolobus*, FEMS Microbiol. Lett. **43**, 327 (1987).

[82] K. A. Taylor, J. F. Deatherage, and L. A. Amos, Structure of the S-layer of *Sulfolobus acidocaldarius*, Nature **299**, 840 (1982).

[83] J. F. Deatherage, K. A. Taylor, and L. A. Amos, Three-dimensional arrangement of the cell wall protein of *Sulfolobus acidocaldarius*, J. Mol. Biol. **167**, 825 (1983).

[84] G. Lembcke, R. Dürr, R. Hegerl, and W. Baumeister, Image analysis and processing of an imperfect two-dimensional crystal: the surface layer of the archaebacterium *Sulfolobus acidocaldarius* re-investigated, J. Microsc. **161**, 263 (1991).

[85] W. Zillig, A. Gierl, G. Schreiber, S. Wunderl, D. Janekovic, K. O. Stetter, and H. P. Klenk, The archaebacterium *Thermofilum pendens* represents a novel genus of the thermophilic, anaerobic sulfur respiring *Thermoproteales*, System. Appl. Microbiol. **4**, 79 (1983).

[86] J. G. Elkins, M. Podar, D. E. Graham, K. S. Makarova, Y. Wolf, L. Randau, B. P. Hedlund, C. Brochier-Armanet, V. Kunin, A. Anderson, I. Lapidus, E. Goltsman, K. Barry, E. V. Koonin, P. Hugenholtz, N. Kyrpides, G. Wanner, P. Richardson, M. Keller, and K. O. Stetter, A korarchaeal genome reveals insights into the evolution of the Archaea, Proc. Natl. Acad. Sci. USA **105**, 8102 (2008).

[87] S. Gfrerer, D. Winkler, J. Novion Ducassou, Y. Counté, R. Rachel, and J. Gescher, A micrarchaeon isolate is covered by a proteinaceous S-layer, Appl. Environ. Microbiol. **88**, e01553 (2022).

[88] R. J. Giannone, H. Huber, T. Karpinets, T. Heimerl, U. Küper, R. Rachel, M. Keller, R. L. Hettich, and M. Podar, Proteomic characterization of cellular and molecular processes that enable the *Nanoarchaeum equitans-Ignicoccus hospitalis* relationship, PLoS ONE **6**, e22942 (2011).

[89] H. Huber, M. J. Hohn, K. O. Stetter, and R. Rachel, The phylum Nanoarcheota: Present knowledge and future perspectives of a unique form of life, Res. Microbiol. **154**, 165 (2003).

[90] V. La Cono, E. Messina, M. Rohde, E. Arcadi, S. Ciordia, F. Crisafi, R. Denaro, M. Ferrer, L. Giuliano, P. N. Golyshin, O. V. Golyshina, J. E. Hallsworth, G. La Spada, M. C. Mena, A. Y. Merkel, M. A. Shevchenko, F. Smedile, D. Y. Sorokin, S. V. Toshchakov, and M. M. Yakimov, Symbiosis between nanohaloarchaeon and haloarchaeon is based on utilization of different polysaccharides, Proc. Natl. Acad. Sci. USA **117**, 20223 (2020).

[91] S. Chu, S. Cavaignac, J. Feutrier, B. M. Phipps, M. Kostrzynska, W. W. Kay, and T. J. Trust, Structure of the tetragonal surface virulence array protein and gene of *Aeromonas salmonicida*, J. Biol. Chem. **266**, 15258 (1991).

[92] J. S. G. Dooley, H. Engelhardt, W. Baumeister, W. W. Kay, and T. J. Trust, Three-dimensional structure of an open form of the surface layer from the fish pathogen *Aeromonas salmonicida*, J. Bacteriol. **171**, 190 (1989).

[93] S. R. Thomas and T. J. Trust, Tyrosine phosphorylation of the tetragonal paracrystalline array of *Aeromonas hydrophila*: Molecular cloning and high-level expression of the S-layer protein gene, J. Mol. Biol. **245**, 568 (1995).

[94] S. Al-Karadaghi, D. N. Wang, and S. Hovmöller, Three-dimensional structure of the crystalline surface layer from *Aeromonas hydrophila*, J. Ultrastruct. Mol. Struct. Res. **101**, 92 (1988).

[95] R. G. E. Murray, J. S. G. Dooley, P. W. Whippey, and T. J. Trust, Structure of an S layer on a pathogenic strain of *Aeromonas hydrophila*, J. Bacteriol. **170**, 2625 (1988).

[96] P. W. Y. Liew, B. C. Jong, and N. Najimudin, Hypothetical protein Avin_16040 as the S-layer protein of

*Azotobacter vinelandii* and its involvement in plant root surface attachment, Appl. Environ. Microbiol. **81**, 7484 (2015).

[97] W. H. Bingle, H. Engelhardt, W. J. Page, and W. Baumeister, Three-dimensional structure of the regular tetragonal surface layer of *Azotobacter vinelandii*, J. Bacteriol. **169**, 5008 (1987).

[98] S. Ali, B. Jenkins, J. Cheng, B. Lobb, X. Wei, S. Egan, T. C. Charles, J. McConkey, B. J. Austin, and A. C. Doxey, Slr4, a newly identified S-layer protein from marine Gammaproteobacteria, is a major biofilm matrix component, Mol. Microbiol. **114**, 979 (2020).

[99] Y. Kawasaki, K. Kurosaki, D. Kan, I. Kazahaya Borges, A. Satake Atagui, M. Sato, K. Kondo, M. Katahira, I. Suzuki, and M. Takeda, Identification and characterization of the S-layer formed on the sheath of *Thiothrix nivea*, Arch. Microbiol. **200**, 1257 (2018).

[100] C. C. Remsen, S. W. Watson, J. B. Waterbury, and H. G. Trüper, Fine structure of *Ectothiorhodospira mobilis* Pelsh, J. Bacteriol. **95**, 2374 (1968).

[101] C. C. Remsen, S. W. Watson, and H. G. Trüper, Macromolecular subunits in the walls of marine photosynthetic bacteria, J. Bacteriol. **103**, 255 (1968).

[102] A. Vazquez Vital, *Surface (S)-layers from methanotrophic bacteria: Composition and function*, Master's thesis, San Diego State University (2022).

[103] V. N. Khmelenina, V. N. Shchukin, A. S. Reshetnikov, I. I. Mustakhimov, N. E. Suzina, B. T. Eshinimaev, and Y. A. Trotsenko, Structural and functional features of methanotrophs from hypersaline and alkaline lakes, Microbiology **79**, 472 (2010).

[104] V. N. Khmelenina, N. E. Suzina, and Y. A. Trotsenko, Surface layers of methanotrophic bacteria, Microbiology **82**, 529 (2013).

[105] E. Kawai, H. Akatsuka, A. Idei, E. Shibatani, and K. Omori, *Serratia marcescens* S-layer protein is secreted extracellularly via an ATP-binding cassette exporter, the Lip system, Mol. Microbiol. **27**, 941 (1998).

[106] T. J. Beveridge and R. G. E. Murray, Surface arrays on the cell wall of *Spirillum metamorphum*, J. Bacteriol. **124**, 1529 (1975).

[107] M. R. Dickson, K. H. Downing, W. H. Wu, and R. M. Glaeser, Three-dimensional structure of the surface layer protein of *Aquaspirillum serpens* VHA determined by electron crystallography, J. Bacteriol. **167**, 1025 (1986).

[108] T. J. Beveridge and R. G. E. Murray, Superficial cell-wall layers on *Spirillum* "Ordal" and their in vitro reassembly, Can. J. Microbiol. **22**, 567 (1976).

[109] A. Shetty, S. Chen, E. I. Tocheva, G. J. Jensen, and W. J. Hickey, Nanopods: A new bacterial structure and mechanism for deployment of outer membrane vesicles, PLoS ONE **6**, e20725 (2011).

[110] H. Engelhardt, S. Gerbi-Rieger, U. Santarius, and W. Baumeister, The three-dimensional structure of the regular surface protein of *Comamonas acidovorans* derived from native outer membranes and reconstituted two-dimensional crystals, Mol. Microbiol. **5**, 1695 (1991).

[111] S. Gajbhiye, E. D. Gonzales, D. B. Toso, N. A. Kirk, and W. J. Hickey, Identification of NpdA as the protein forming the surface layer in *Paracidovorax citrulli* and evidence of its occurrence as a surface layer protein in diverse genera of the betaproteobacteria and gammaproteobacteria, Access Microbiology **5**, 000685 (2023).

[112] A. Gilchrist, J. A. Fisher, and J. Smit, Nucleotide sequence analysis of the gene encoding the *Caulobacter crescentus* paracrystalline surface layer protein, Can. J. Microbiol. **38**, 193 (1992).

[113] M. R. J. Salton and R. C. Williams, Electron microscopy of the cell walls of *Bacillus megaterium* and *Rhodospirillum rubrum*, Biochim. Biophys. Acta **14**, 455 (1954).

[114] B. Wang, E. Kraig, and Kolodrubetz, A new member of the S-layer protein family: Characterization of the crs gene from *Campylobacter rectus*, Infect. Immun. **66**, 1521–1526 (1998).

[115] T. Dokland, I. Olsen, G. Farrants, and B. V. Johansen, Three-dimensional structure of the surface layer of *Wolinella recta*, Oral Microbiol. Immunol. **5**, 162 (1990).

[116] A. Sjögren, S. Hovmöller, G. Farrantis, H. Ranta, M. Haapasalo, K. Ranta, and K. Lounatmaa, The structure of crystalline bacterial surface layers, J. Bacteriol. **164**, 1278 (1985).

[117] C. M. Fletcher, M. J. Coyne, D. L. Bentley, O. F. Villa, and L. E. Comstock, Phase-variable expression of a family of glycoproteins imparts a dynamic surface to a symbiont in its human intestinal ecosystem, Proc. Natl. Acad. Sci. USA **104**, 2413 (2007).

[118] M. C. F. van Teeseling, N. M. de Almeida, A. Klingl, D. R. Speth, H. J. M. Op den Camp, R. Rachel, M. S. M. Jetten, and L. van Niftrik, A new addition to the cell plan of anammox bacteria: "*Candidatus* Kuenenia stuttgartiensis" has a protein surface layer as the outermost layer of the cell, J. Bacteriol. **196**, 80 (2014).

[119] C. Ding and L. Adrian, Comparative genomics in "*Candidatus* Kuenenia stuttgartiensis" reveal high genomic plasticity in the overall genome structure, CRISPR loci and surface proteins, BMC Genom. **21**, 851 (2020).

[120] L. Gambelli, R. Mesman, W. Versantvoort, C. A. Diebolder, A. Engel, W. Evers, M. S. M. Jetten, M. Pabst, B. Daum, and L. van Niftrik, The polygonal cell shape and surface protein layer of anaerobic methane-oxidizing *Methylomirabilis lanthanidiphila* bacteria, Front. Microbiol. **12**, 766527 (2021).

[121] P. Messner, K. Steiner, K. Zarschler, and C. Schäffer, S-layer nanoglycobiology of bacteria, Carbohydr. Res. **343**, 1934 (2008).

[122] K. Meier-Stauffer, H.-J. Busse, F. A. Rainey, J. Burghardt, A. Scheberl, F. Hollaus, B. Kuen, A. Makristathis, U. B. Sleytr, and P. Messner, Description of *Bacillus thermoaerophilus* sp. nov., to include sugar beet isolates and *Bacillus brevis* ATCC 12990, Int. J. Syst. Bacteriol. **46**, 532 (1996).

[123] S. K. Burley and R. G. E. Murray, Structure of the regular surface layer of *Bacillus polymyxa*, Can. J. Microbiol. **29**, 775 (1983).

[124] H. Engelhardt, Z. Cejka, and W. Baumeister, Three-dimensional structure of surface layers from various *Bacillus* and *Clostridium* species, in *Crystalline Bacterial Cell Surface Layers*, edited by U. B. Sleytr, P. Messner, and M. Sára (Springer, Berlin, Heidelberg, 1988) pp. 87–91.

[125] A. Schenk and M. Aragno, *Bacillus schlegelii*, a new species of thermophilic, facultatively chemolithoautotrophic bacterium oxidizing molecular hydrogen, J. Gen. Microbiol. **115**, 333 (1979).

[126] N. Ilk, C. Völlenkle, E. M. Egelseer, A. Breitweiser, U. B. Sleytr, and M. Sára, Molecular characterization of the S-layer gene, *sbpA*, of *Bacillus sphaericus* CCM 2177 and production of a functional S-layer fusion protein with the ability to recrystallize in a defined orientation while presenting the fused allergen, Appl. Environ. Microbiol. **68**, 3251 (2002).

[127] J. E. Norville, D. F. Kelly, T. F. Knight Jr., A. M. Belcher, and T. Walz, 7 Å projection map of the S-layer protein sbpA obtained with trehalose-embedded monolayer crystals, J. Struct. Biol. **160**, 313 (2007).

[128] U. B. Sleytr, C. Huber, N. Ilk, D. Pum, B. Schuster, and E. M. Egelseer, S-layers as a tool kit for nanobiotechnological applications, FEMS Microbiol. Lett. **267**, 131 (2007).

[129] R. D. Bowditch, P. Baumann, and A. A. Yousten, Cloning and sequencing of the gene encoding a 125-kilodalton surface-layer protein from *Bacillus sphaericus* 2362 and of a related cryptic gene, J. Bacteriol. **171**, 4178 (1989).

[130] N. K. Li, H. S. Kim, J. A. Nash, M. Lim, and Y. G. Yingling, Nanoscale mono- and multi-layer cylinder structures formed by recombinant S-layer proteins of mosquitocidal *Bacillus sphaericus* C3-41, Appl. Microbiol. Biotechnol. **97**, 7275 (2013).

[131] F. L. Lederer, U. Weinert, T. J. Günther, J. Raff, S. Weiß, and K. Pollmann, Identification of multiple putative S-layer genes partly expressed by *Lysinibacillus sphaericus* JG-B53, Microbiology **159**, 1097 (2013).

[132] S. C. Holt and E. R. Leadbetter, Comparative ultrastructure of selected aerobic spore-forming bacteria: a freeze-etching study, Bacteriol. Rev. **33**, 346 (1969).

[133] P. M. Ryzhkov, K. Ostermann, and G. Rödel, Isolation, gene structure, and comparative analysis of the S-layer gene sslA of *Sporosarcina ureae* ATCC 13881, Genetica **131**, 255 (2007).

[134] H. Engelhardt, W. O. Saxton, and W. Baumeister, Three-dimensional structure of the tetragonal surface layer of *Sporosarcina ureae*, J. Bacteriol. **168**, 309 (1986).

[135] M. Suhr, F. L. Lederer, T. J. Günther, J. Raff, and K. Pollmann, Characterization of three different unusual S-layer proteins from *Viridibacillus arvi* JG-B58 that exhibits two super-imposed S-layer proteins, PLoS ONE **11**, e0156785 (2016).

[136] A. Tsuboi, R. Uchihi, T. Adachi, T. Sasaki, S. Hayakawa, H. Yamagata, N. Tsukagoshi, and S. Udaka, Characterization of the genes for the hexagonally arranged surface layer proteins in protein-producing *Bacillus brevis* 47: complete nucleotide sequence of the middle wall protein gene, J. Bacteriol. **170**, 935 (1988).

[137] A. Tsuboi, H. Engelhardt, U. Santarius, N. Tsukagoshi, S. Udaka, and W. Baumeister, Three-dimensional structure of the surface protein layer (MW layer) of *Bacillus brevis* 47, J. Ultrastruct. Mol. Struct. Res. **102**, 178 (1989).

[138] A. Tsuboi, R. Uchihi, H. Engelhardt, H. Hattori, S. Shimuzu, N. Tsukagoshi, and S. Udaka, In vitro reconstitution of a hexagonal array with a surface layer protein synthesized by *Bacillus subtilis* harboring the surface layer protein gene from *Bacillus brevis* 47, J. Bacteriol. **171**, 6747 (1989).

[139] H. J. Boot, C. P. Kolen, J. M. van Noort, and P. H. Pouwels, S-layer protein of *Lactobacillus acidophilus* ATCC 4356: purification, expression in *Escherichia coli*, and nucleotide sequence of the corresponding gene, J. Bacteriol. **175**, 6089 (1993).

[140] E. Smit, F. Oling, R. Demel, B. Martinez, and P. H. Pouwels, The S-layer protein of *Lactobacillus acidophilus* ATCC 4356: Identification and characterisation of domains responsible for S-protein assembly and cell wall binding, J. Mol. Biol. **305**, 245 (2001).

[141] T. Sagmeister, N. Gubensäk, C. Buhlheller, C. Grininger, M. Eder, A. Dordić, C. Millán, A. Medina, P. Sánchez-Murcia, F. Berni, U. Hynönen, D. Vejzović, E. Damisch, N. Kulminskaya, L. Petrowitsch, M. Oberer, A. Palva, N. Malanović, J. Codée, W. Keller, I. Usón, and T. Pavkov-Keller, The molecular architecture of Lactobacillus S-layer: Assembly and attachment to teichoic acids, ResearchSquare.

[142] B. Kuen, U. B. Sleytr, and W. Lubitz, Sequence analysis of the sbsA gene encoding the 130-kDa surface-layer protein of *Bacillus stearothermophilus* strain PV72, Gene **145**, 115 (1994).

[143] U. B. Sleytr, M. Sára, Z. Küpcü, and P. Messner, Structural and chemical characterization of S-layers of selected strains of *Bacillus stearothermophilus* and *Desulfotomaculum nigrificans*, Arch. Microbiol. **146**, 19 (1986).

[144] K. C. Schuster, H. F. Mayer, R. Kieweg, W. A. Hampel, and M. Sára, A synthetic medium for continuous culture of the S-layer carrying *Bacillus stearothermophilus* PV 72 and studies on the influence of growth conditions on cell wall properties, Biotechnol. Bioeng. **48**, 66 (1995).

[145] M. Jarosch, E. M. Egelseer, D. Mattanovich, U. B. Sleytr, and M. Sára, S-layer gene sbsC of *Bacillus stearothermophilus* ATCC 12980: molecular characterization and heterologous expression in *Escherichia coli*, Microbiology **146**, 273 (2000).

[146] E. M. Egelseer, K. Leitner, M. Jarosch, C. Hotzy, S. Zayni, U. B. Sleytr, and M. Sára, The S-layer proteins of two *Bacillus stearothermophilus* wild-type strains are bound via their N-terminal region to a secondary cell wall polymer of identical chemical composition, J. Bacteriol. **180**, 1488 (1998).

[147] T. Pavkov, E. M. Egelseer, M. Tesarz, D. I. Svergun, U. B. Sleytr, and W. Keller, The structure and binding behavior of the bacterial cell surface layer protein SbsC, Structure **16**, 1226 (2008).

[148] C. Schäffer, T. Wugedtisch, H. Kählig, A. Scheberl, S. Zayni, and P. Messner, The surface layer (S-layer) glycoprotein of *Geobacillus stearothermophilus* NRS 2004/3a, J. Biol. Chem. **277**, 6230 (2002).

[149] I. Etienne-Toumelin, J.-C. Sirard, E. Duflot, M. Mock, and A. Fouet, Characterization of the *Bacillus anthracis* S-layer: Cloning and sequencing of the structural gene, J. Bacteriol. **177**, 614 (1995).

[150] A. Fioravanti, F. Van Hauwermeiren, S. E. Van der Verren, W. Jonckheere, A. Goncalves, E. Pardon, J. Steyaert, H. De Greve, M. Lamkanfi, and H. Remaut, Structure of S-layer protein Sap reveals a mechanism for therapeutic intervention in anthrax, Nat. Microbiol. **4**, 1805 (2019).

[151] S. Mesnage, E. Tosi-Couture, M. Mock, P. Gounon, and A. Fouet, Molecular characterization of the *Bacillus anthracis* main S-layer component: evidence that it is the

major cell-associated antigen, Mol. Microbiol. **23**, 1147 (1997).

[152] X.-Y. Wang, D.-B. Wang, Z.-P. Zhang, L.-J. Bi, J.-B. Zhang, W. Ding, and X.-E. Zhang, A S-layer protein of *Bacillus anthracis* as a building block for functional protein arrays by in vitro self-assembly, Small **11**, 5826 (2015).

[153] L. Poppinga, B. Janesch, A. Funfhaus, E. Sekot, G. aand Garcia-Gonzalez, G. Hertlein, K. Hedtke, C. Schäffer, and E. Genersch, Identification and functional analysis of the S-layer protein SplA of *Paenibacillus larvae*, the causative agent of American foulbrood of honey bees, PLoS Pathog. **8**, e1002716 (2012).

[154] A. Slobodkin, A.-L. Reysenbach, F. Mayer, and J. Wiegel, Isolation and characterization of the homoacetogenic thermophilic bacterium *Moorella glycerini* sp. nov., Int. J. Syst. Bacteriol. **47**, 969 (1997).

[155] C. Holliger, D. Hahn, H. Harmsen, W. Ludwig, B. Shumacher, W.and Tindall, F. Vazquez, N. Weiss, and A. J. B. Zehnder, *Dehalobacter restrictus* gen. nov. and sp. nov., a strictly anaerobic bacterium that reductively dechlorinates tetra- and trichloroethene in an anaerobic respiration, Arch. Microbiol. **169**, 313 (1998).

[156] C. L. Woodside, H. Engelhardt, and W. Baumeister, The tetragonal surface layer of *Clostridium aceticum*: three-dimensional structure and comparison with the hexagonal layer of *Clostridium thermohydrosulfuricum*, Eur. J. Cell Biol. **42**, 211 (1986).

[157] M. J. Thornley, A. M. Glauert, and U. B. Sleytr, Structure and assembly of bacterial surface layers composed of regular arrays of subunits, Phil. Trans. R. Soc. Lond. B. **268**, 147 (1974).

[158] R. A. Crowther and U. B. Sleytr, An analysis of the fine structure of the surface layers from two strains of *Clostridia*, including correction for distorted images, J. Ultrastruct. Res. **58**, 41 (1977).

[159] Z. Cejka and W. Baumeister, Three-dimensional structure of the surface protein of *Clostridium thermosaccharolyticum*, FEMS Microbiol. Lett. **44**, 13 (1987).

[160] A. Sjögren, D. N. Wang, S. Hovmöller, M. Haapasalo, H. Ranta, E. Kerosuo, H. Jousimies-Somer, and K. Lounatmaa, The three-dimensional structures of S-layers of two novel *Eubacterium* species isolated from inflammatory human processes, Mol. Microbiol. **2**, 81 (1988).

[161] R. Ristl, K. Steiner, K. Zarschler, S. Zayni, P. Messner, and C. Schäffer, The S-layer glycome—adding to the sugar coat of bacteria, Int. J. Microbiol. **2011**, 127870 (2011).

[162] Z. Cejka, R. Hegerl, and W. Baumeister, Three-dimensional structure of the surface layer protein of *Clostridium thermohydrosulfuricum*, J. Ultrastruct. Res. **96**, 1 (1986).

[163] J. Peters, M. Peters, F. Lottspeich, , and W. Baumeister, S-layer protein gene of *Acetogenium kivui*: Cloning and expression in *Escherichia coli* and determination of the nucleotide sequence, J. Bacteriol. **171**, 6307 (1989).

[164] A. Lupas, H. Engelhardt, J. Peters, U. Santarius, S. Volker, and W. Baumeister, Domain structure of the *Acetogenium kivui* surface layer revealed by electron crystallography and sequence analysis, J. Bacteriol. **176**, 1224 (1994).

[165] D. H. Currie, A. M. Guss, C. D. Herring, R. J. Giannone, C. M. Johnson, P. K. Lankford, S. D. Brown, R. L. Hettich, and L. R. Lynd, Profile of secreted hydrolases, associated proteins, and SlpA in *Thermoanaerobacterium saccharolyticum* during the degradation of hemicellulose, Appl. Environ. Microbiol. **80**, 5001 (2014).

[166] M. Lemaire, I. Miras, P. Gounon, and P. Béguin, Identification of a region responsible for binding to the cell wall within the S-layer protein of *Clostridium thermocellum*, Microbiology **144**, 211 (1998).

[167] J. L. Peyret, N. Bayan, G. Joliff, T. Gulik-Krzywicki, L. Mathieu, E. Schechter, and G. Leblon, Characterization of the cspB gene encoding PS2, an ordered surface-layer protein in *Corynebacterium glutamicum*, Mol. Microbiol **9**, 97 (1993).

[168] S. Scheuring, H. Stahlberg, M. Chami, C. Houssin, J.-L. Rigaud, and A. Engel, Charting and unzipping the surface layer of *Corynebacterium glutamicum* with the atomic force microscope, Mol. Microbiol. **44**, 675 (2002).

[169] N. Hansmeier, F. W. Bartels, R. Ros, D. Anselmetti, A. Tauch, A. Pühler, and J. Kalinowski, Classification of hyper-variable *Corynebacterium glutamicum* surface-layer proteins by sequence analyses and atomic force microscopy, J. Biotechnol. **112**, 177 (2004).

[170] W. Baumeister, F. Karrenberg, R. Rachel, A. Engel, B. Ten Heggeler, and W. O. Saxton, The major cell envelope protein of *Micrococcus radiodurans* (R1), Eur. J. Biochem. **125**, 535 (1982).

[171] J. R. Castón, J. Berenguer, E. Kocsis, and J. L. Carrascosa, Three-dimensional structure of different aggregates built up by the s-layer protein of *Thermus thermophilus*, J. Struct. Biol. **113**, 164 (1994).

[172] R. M. Morris, S. Sowell, D. Barorsky, S. Zinder, and R. Richardson, Transcription and mass-spectroscopic proteomic studies of electron transport oxidoreductases in *Dehalococcoides ethenogenes*, Environ. Microbiol. **8**, 1499 (2006).

[173] D. L. Sexton, G. Chen, F. K. Murdoch, A. Hashimi, F. E. Löffler, and E. I. Tocheva, Ultrastructure of organohalide-respiring *Dehalococcoidia* revealed by cryo-electron tomography, Appl. Environ. Microbiol. **88**, e01906 (2022).

[174] M. Palmer, J. K. Covington, E.-M. Zhou, S. C. Thomas, N. Habib, C. O. Seymour, D. Lai, J. Johnston, A. Hashimi, J.-Y. Jiao, A. R. Muok, L. Liu, W.-D. Xian, X.-Y. Zhi, M.-M. Li, L. P. Silva, B. P. Bowen, K. Louie, A. Briegel, J. Pett-Ridge, P. K. Weber, E. I. Tocheva, T. Woyke, T. R. Northern, X. Mayali, W.-J. Li, and B. P. Hedlund, Thermophilic *Dehalococcoidia* with unusual traits shed light on an unexpected past, ISME J. **17**, 952 (2023).

[175] W.-W. Li, M.-S. Xia, W.-B. Li, L.-W. Liu, Y. Yang, and J.-H. Li, Characterization of a putative S-layer protein of a colonial *Microcystis* strain, Curr. Microbiol. **75**, 173 (2018).

[176] J. McCarren, J. Heuser, R. Roth, N. Yamada, M. Martone, and B. Brahamsha, Inactivation of *swmA* results in the loss of an outer cell layer in a swimming *Synechococcus* strain, J. Bacteriol. **187**, 224 (2005).

[177] C. Trautner and W. F. J. Vermaas, The *sll1951* gene encodes the surface layer protein of *Synechocystis* sp. strain PCC 6803, J. Bacteriol. **195**, 5370 (2013).

[178] K. Kuroda, M. Nakajima, R. Nakai, Y. Hirakata, S. Kagemasa, K. Kubota, T. Q. P. Noguchi, K. Yamamoto, H. Satoh, M. K. Nobu, and T. Nirihiro, Microscopic and metatranscriptomic analyses revealed

unique cross-domain parasitism between phylum *Candidatus* Patescibacteria/candidate phyla radiation and methanogenic archaea in anaerobic ecosystems, mBio **15**, e03102 (2024).

[179] X. Chen, O. Molenda, C. T. Brown, C. R. A. Toth, S. Guo, F. Luo, J. Howe, C. L. Nesbø, C. He, E. A. Montabana, J. H. D. Cate, J. F. Banfield, and E. A. Edwards, "*Candidatus* Nealsonbacteria" are likely biomass recycling ectosymbionts of methanogenic archaea in a stable benzene-degrading enrichment culture, Appl. Environ. Microbiol. **89**, e00025 (2023).

[180] D. H. Parks, M. Chuvochina, C. Rinke, A. J. Mussig, P.-A. Chaumeil, and P. Hugenholtz, GTDB: an ongoing census of bacterial and archaeal diversity through a phylogenetically consistent, rank normalized and complete genome-based taxonomy, Nucleic Acids Res. **50**, D785 (2022).

[181] I. Letunic and P. Bork, Interactive Tree of Life (iTOL) v6: recent updates to the phylogenetic tree display and annotation tool, Nucleic Acids Res. , gkae268 (2024).

[182] UniProt: the Universal Protein Knowledgebase in 2023, Nucleic Acids Res. **51**, D523 (2023).

[183] C. Camacho, G. Coulouris, V. Avagyan, N. Ma, J. Papadopoulos, K. Bealer, and T. L. Madden, BLAST+: architecture and applications, BMC Bioinformatics **10**, 1 (2009).

[184] T. Frickey and A. Lupas, CLANS: a Java application for visualizing protein families based on pairwise similarity, Bioinformatics **20**, 3702 (2004).

[185] L. Zimmermann, A. Stephens, S.-Z. Nam, D. Rau, J. Kübler, M. Lozajic, F. Gabler, J. Söding, A. N. Lupas, and V. Alva, A completely reimplemented MPI bioinformatics toolkit with a new HHpred server at its core, J. Mol. Biol. **430**, 2237 (2018).

[186] T. Paysan-Lafosse, M. Blum, S. Chuguransky, T. Grego, B. L. Pinto, G. A. Salazar, M. L. Bileschi, P. Bork, A. Bridge, L. Colwell, *et al.*, InterPro in 2022, Nucleic Acids Res. **51**, D418 (2023).

[187] M. Steinegger, M. Meier, M. Mirdita, H. Vöhringer, S. J. Haunsberger, and J. Söding, HH-suite3 for fast remote homology detection and deep protein annotation, BMC Bioinformatics **20**, 1 (2019).

[188] The Protein Data Bank, http://www.rcsb.org/pdb/.

[189] Evolutionary Classification of Protein Domains, http://prodata.swmed.edu/ecod/.

[190] J. Mistry, S. Chuguransky, L. Williams, M. Qureshi, G. A. Salazar, E. L. Sonnhammer, S. C. Tosatto, L. Paladin, S. Raj, L. J. Richardson, *et al.*, Pfam: The protein families database in 2021, Nucleic Acids Res. **49**, D412 (2021).

[191] F. Teufel, J. J. Almagro Armenteros, A. R. Johansen, M. H. Gíslason, S. I. Pihl, K. D. Tsirigos, O. Winther, S. Brunak, G. von Heijne, and H. Nielsen, SignalP 6.0 predicts all five types of signal peptides using protein language models, Nat. Biotechnol. **40**, 1023 (2022).

[192] A. Punjani, J. L. Rubinstein, D. J. Fleet, and M. A. Brubaker, cryoSPARC: algorithms for rapid unsupervised cryo-EM structure determination, Nat. Methods **14**, 290 (2017).

[193] E. C. Meng, T. D. Goddard, E. F. Pettersen, G. S. Couch, Z. J. Pearson, J. H. Morris, and T. E. Ferrin, USCF ChimeraX: Tools for structure building and analysis, Protein Sci. **32**, 34792 (2023).

[194] A. von Kügelgen, H. Tang, G. G. Hardy, D. Kureisaite-Ciziene, Y. V. Brun, P. J. Stansfield, C. V. Robinson, and T. A. M. Bharat, *In situ* structure of an intact lipopolysaccharide-bound bacterial surface layer, Cell **180**, 348 (2020).

[195] D. N. Mastronarde, SerialEM: A program for automated tilt series acquisition on Tecnai microscopes using prediction of specimen position, Microsc. Microanal. **9**, 1182 (2003).

[196] J. R. Kremer, D. N. Mastronarde, and J. R. McIntosh, Computer visualization of three-dimensional image data using IMOD, J. Struct. Biol. **116**, 71 (1996).

[197] A. Durán-Viseras, C. Sánchez-Porro, D. Viver, K. T. Konstantinidis, and A. Ventosa, Discovery of the streamlined haloarchaeon *Halorutilus salinus*, comprising a new order widespread in hypersaline environments across the world, mSystems **8**, e01198 (2023).

[198] W. Zillig, K. O. Stetter, W. Schäfer, D. Janekovic, S. Wunderl, I. Holz, and P. Palm, Thermoproteales: A novel type of extremely thermoacidophilic anaerobic archaebacteria isolated from Icelandic solfataras, Zbl. Bakt. Hyg., I. Abt. Orig. C **2**, 205 (1981).

[199] A. K. Perras, B. Daum, C. Ziegler, L. K. Takahashi, M. Ahmed, G. Wanner, A. Klingl, G. Leitinger, D. Kolb-Lenz, S. Gribaldo, A. Auerbach, M. Mora, A. J. Probst, A. Bellack, and C. Moissl-Eichinger, S-layers at second glance? Altiarchaeal grappling hooks (hami) resemble archaeal S-layer proteins in structure and sequence, Front. Microbiol. **6**, 543 (2015).

[200] C. Moissy, R. Rachel, A. Briegel, H. Engelhardt, and R. Huber, The unique structure of archaeal 'hami', highly complex cell appendages with nano-grappling hooks, Molec. Microbiol. **56**, 361 (2005).

[201] A.-L. Reysenbach, Y. Liu, A. B. Banta, T. J. Beveridge, J. D. Kirshtein, S. Schouten, M. K. Tivey, K. L. Von Damm, and M. A. Voytek, A ubiquitous thermoacidophilic archaeon from deep-sea hydrothermal vents, Nature **442**, 444 (2006).

[202] M. A. Khomyakova, A. Y. Merkel, D. D. Mamiy, A. A. Klyukina, and A. I. Slobodkin, Phenotypic and genomic characterization of *Bathyarchaeum tardum* gen. nov., sp. nov., a cultivated representative of the archaeal class *Bathyarchaeia*, Front. Microbiol. **14**, 1214631 (2023).

[203] R. Hatzenpichler, A. Kohtz, V. Krukenberg, N. Petrosian, Z. Jay, and M. Pilhofer, Cultivation and visualization of a methanogen of the phylum thermoproteota, ResearchSquare (2023).

[204] E. Blöchl, R. Rachel, S. Burggraf, D. Hafenbradl, H. W. Jannasch, and K. O. Stetter, *Pyrolobus fumarii*, gen. and sp. nov., represents a novel group of archaea, extending the upper temperature limit for life to 113oc, Extremophiles **1**, 14 (1997).

[205] C. M. Debieux, E. J. Dridge, C. M. Mueller, P. Splatt, K. Paskiewicz, I. Knight, H. Florance, J. Love, R. W. Titball, R. J. Lewis, D. J. Richardson, and C. S. Butler, A bacterial process for seleniumnanosphere assembly, Proc. Natl. Acad. Sci. USA **108**, 13480 (2011).

[206] J. W. Austin, A. Engel, R. G. E. Murray, and U. Aebi, Structural analysis of the S-layer of *Lampropedia hyalina*, J. Ultrastruct. Mol. Struct. Res. **102**, 255 (1989).

[207] E. M. Egelseer, T. Danhorn, M. Pleschberger, C. Hotzy, U. B. Sleytr, and M. Sára, Characterization of an S-layer glycoprotein produced in the course of S-layer varia-

tion of *Bacillus stearothermophilus* ATCC 12980 and sequencing and cloning of the sbsD gene encoding the protein moiety, Arch. Microbiol. **177**, 70 (2001).

[208] M. M. Faraldo, M. A. de Pedro, and J. Berenguer, Sequence of the S-layer gene of *Thermus thermophilus* HB8 and functionality of its promoter in *Escherichia coli*, J. Bacteriol. **174**, 7458 (1992).

[209] G. Olabarría, L. A. Fernández-Herrero, J. L. Carrascosa, and J. Berenguer, *slpM*, a gene coding for an "S-layer-like array" overexpressed in S-layer mutants of *Thermus thermophilus* HB8, J. Bacteriol. **178**, 357 (1996).

[210] V. A. Gaisin, R. Kooger, D. S. Grouzdev, V. M. Gorlenko, and M. Pilhofer, Cryo-electron tomography reveals the complex ultrastructural organization of multicellular filamentous *Chloroflexota* (*Chloroflexi*) bacteria, Front. Microbiol. **11**, 1373 (2020).

[211] R. Rachel, D. Pum, J. Šmarda, D. Šmajs, J. Komrska, V. Krzyžánek, G. Rieger, and K. O. Stetter, II. Fine structure of S-layers, FEMS Microbiol. Rev. **20**, 13 (1997).

[212] J. Šmarda, D. Šmajs, J. Komrska, and V. Krzyžánek, S-layers on cell walls of cyanobacteria, Micron **33**, 252 (2002).

[213] V. Kryžánek, Analysis of continuously distorted quasiperiodic images: two-dimensional reconstruction of S layers of cyanobacteria, Opt. Eng. **39**, 872 (2000).

[214] C. He, R. Keren, M. L. Whittaker, I. F. Farag, J. A. Doudna, J. H. D. Cate, and J. F. Banfield, Genome-resolved metagenomics reveals site-specific diversity of episymbiotic CPR bacteria and DPANN archaea in groundwater ecosystems, Nat. Microbiol. **6**, 354 (2021).

[215] B. K. Ganser-Pornillos, A. Cheng, and M. Yeager, Structure of full-length HIV-1 CA: A model for the mature capsid lattice, Cell **131**, 70 (2007).

[216] K. A. Dryden, C. S. Crowley, S. Tanaka, T. O. Yeates, and M. Yeager, Two-dimensional crystals of carboxysome shell proteins recapitulate the hexagonal packing of three-dimensional crystals, Protein Sci. **18**, 2629 (2009).

[217] M. Sutter, M. Faulkner, C. Aussignargues, B. C. Paasch, S. a. Barrett, C. A. Kerfeld, and L.-N. Lui, Visualization of bacterial microcompartment facet assembly using high-speed atomic force microscopy, Nano Lett. **16**, 1590 (2016).

[218] C. A. Kerfeld, C. Aussignargues, J. Zarzycki, F. Cai, and M. Sutter, Bacterial microcompartments, Nat. Rev. Microbiol. **16**, 277 (2018).

[219] A. M. Stewart, K. L. Stewart, T. O. Yeates, and T. A. Bobik, Advances in the world of bacterial microcompartments, Trends Biochem. Sci. **46**, 406 (2021).

[220] T. W. Giessen, Encapsulins, Annu. Rev. Biochem. **91**, 353 (2022).

[221] L. Kailas, C. Terry, N. Abbott, R. Taylor, N. Mullin, S. B. Tzokov, S. J. Todd, B. A. Wallace, J. K. Hobbs, A. Moir, and P. A. Bullough, Surface architecture of endospores of the *Bacillus cereus/anthracis/ thuringiensis* family at the subnanometer scale, Proc. Natl. Acad. Sci. USA **108**, 16014 (2011).

[222] C. Terry, S. Jiang, D. S. Radford, Q. Wan, S. Tzokov, A. Moir, and P. A. Bullough, Molecular tiling on the surface of a bacterial spore – the exosporium of the *Bacillus anthracis/cereus/thuringiensis* group, Mol. Microbiol. **104**, 539 (2017).

[223] T. K. Janganan, N. Mullin, A. Dafis-Sagarmendi, J. Brunt, S. B. Tzokov, S. Stringer, A. Moir, R. R. Chaudhuri, R. P. Fagan, J. K. Hobbs, and P. A. Bullough, Architecture and self-assembly of *Clostridium sporogenes* and *Clostridium botulinum* spore surfaces illustrate a general protective strategy across spore formers, mSphere **5**, e00424 (2020).

[224] O. Pornillos, B. K. Ganser-Pornillos, B. N. Kelly, Y. Hua, F. G. Whitby, C. D. Stout, W. I. Sundquist, C. P. Hill, and M. Yeager, X-ray structures of the hexameric building block of the HIV capsid, Cell **137**, 1282 (2009).

[225] G. Zhao, J. R. Perilla, E. L. Yufenyuy, X. Meng, B. Chen, J. Ning, J. Ahn, A. M. Gronenborn, K. Schulten, C. Aiken, and P. Zhang, Mature HIV-1 capsid structure by cryo-electron microscopy and all-atom molecular dynamics, Nature **497**, 643 (2013).

[226] M. Nandhagopal, A. A. Simpson, J. R. Gurnon, X. Yan, T. S. Baker, M. V. Graves, J. L. Van Etten, and M. G. Rossmann, The structure and evolution of the major capsid protein of a large, lipid-containing DNA virus, Proc. Natl. Acad. Sci. USA **99**, 14758 (2002).

[227] Q. Fang, D. Zhu, I. Agarkova, J. Adhikari, T. Klose, Y. Liu, Z. Chen, Y. Sun, M. L. Gross, J. L. Van Etten, X. Zhang, and M. G. Rossmann, Near-atomic structure of a giant virus, Nat. Commun. **10**, 388 (2019).

[228] J. Hyun, H. Matsunami, T. G. Kim, and M. Wolf, Assembly mechanism of the pleomorphic immature poxvirus scaffold, Nat. Commun. **13**, 1704 (2022).

[229] S. A. Wynne, R. A. Crowther, and A. G. W. Leslie, The crystal structure of the human hepatitis B virus capsid, Mol. Cell **3**, 771 (1999).

[230] R. Ouyang, A. R. Costa, C. K. Cassidy, A. Otwinowksa, V. C. J. Williams, A. Latka, P. J. Stansfield, Z. Drulis-Kawa, Y. Briers, D. M. Pelt, S. J. J. Brouns, and A. Briegel, High-resolution reconstruction of a jumbo-bacteriophage infecting capsulated bacteria using hyperbranched tail fibers, Nat. Commun. **13**, 7241 (2015).

[231] F. Li, C.-F. D. Hou, R. K. Lokareddy, R. Yang, F. Forti, F. Briani, and G. Cingolani, High-resolution cryo-EM structure of the *Pseudomonas* bacteriophage E217, Nat. Commun. **14**, 4052 (2023).

[232] Z. Cui, K. V. Gorzelnik, J.-Y. Chang, C. Langlais, J. Jakana, R. Young, and J. Zhang, Structures of Q$\beta$ virions, virus-like particles, and the Q$\beta$-MurA complex reveal internal coat proteins and the mechanism of host lysis, Proc. Natl. Acad. Sci. USA **114**, 11697 (2017).

[233] K. Valegård, L. Liljas, K. Fridborg, and T. Unge, The three-dimensional structure of the bacterial virus MS2, Nature **345**, 36 (1990).

[234] Q. Xie and M. S. Chapman, Canine parvovirus capsid structure, analyzed at 2.9 Å resolution, J. Mol. Biol. **264**, 497 (1996).

[235] T. G. Laughlin, A. Deep, A. M. Prichard, C. Seitz, Y. Gu, E. Erustun, S. Suslov, K. Khanna, E. A. Birkholz, E. Armbruster, J. A. McCammon, R. E. Amaro, J. Pogliano, K. D. Corbett, and E. Villa, Architecture and self-assembly of the jumbo bacteriophage nuclear shell, Nature **608**, 429 (2022).

[236] L. F. B. Christensen, L. M. Hansen, K. Finster, G. Christiansen, P. H. Nielsen, D. E. Otzen, and M. S. Dueholm, The sheaths of *Methanospirillum* are made of a new type of amyloid protein, Front. Microbiol. **9**, 2729 (2018).

[237] M. S. Dueholm, P. Larsen, K. Finster, M. R. Stenvang, G. Christiansen, B. S. Vad, A. Bøggild, D. E. Otzen, and P. H. Nielsen, The tubular sheaths encasing *Methanosaeta thermophila* filaments are functional amyloids, J. Biol. Chem. **290**, 20590 (2015).

[238] G. B. Patel, G. D. Sprout, R. W. Humphrey, and T. J. Beveridge, Comparative analyses of the sheath structures of *Methanothrix concilii* GP6 and *Methanospirillum hungatei* strains GP1 and JF1, Can. J. Microbiol. **32**, 623 (1986).

[239] T. J. Beveridge, M. Stewart, R. J. Doyle, and G. D. Sprott, Unusual stability of the *Methanospirillum hungatei* sheath, J. Bacteriol. **162**, 728 (1985).

[240] H. Wang, J. Zhang, D. Toso, S. Liao, F. Sedighian, R. Gunsalus, and Z. H. Zhou, Hierarchical organization and assembly of the archaeal cell sheath from an amyloid-like protein, Nat. Comm. **14**, 6720 (2023).

[241] R. J. Blackler, A. López-Guzán, F. F. Hager, B. Janesch, G. Martinz, S. M. L. Gagnon, O. Haji-Ghassemi, P. Kosma, P. Messner, C. Schäffer, and S. V. Evans, Structural basis of cell wall anchoring by SLH domains in *Paenibacillus alvei*, Nat. Commun. **9**, 3120 (2018).
